# Inferring long-term effective population size with Mutation-Selection models

**DOI:** 10.1101/2021.01.13.426421

**Authors:** T. Latrille, V. Lanore, N. Lartillot

## Abstract

Mutation-selection phylogenetic codon models are grounded on population genetics first principles and represent a principled approach for investigating the intricate interplay between mutation, selection and drift. In their current form, mutation-selection codon models are entirely characterized by the collection of site-specific amino-acid fitness profiles. However, thus far, they have relied on the assumption of a constant genetic drift, translating into a unique effective population size (*N*_e_) across the phylogeny, clearly an unreasonable hypothesis. This assumption can be alleviated by introducing variation in *N*_e_ between lineages. In addition to *N*_e_, the mutation rate (*μ*) is susceptible to vary between lineages, and both should co-vary with life-history traits (LHTs). This suggests that the model should more globally account for the joint evolutionary process followed by all of these lineage-specific variables (*N*_e_, *μ*, and LHTs). In this direction, we introduce an extended mutation-selection model jointly reconstructing in a Bayesian Monte Carlo framework the fitness landscape across sites and long-term trends in *N*_e_, *μ* and LHTs along the phylogeny, from an alignment of DNA coding sequences and a matrix of observed LHTs in extant species. The model was tested against simulated data and applied to empirical data in mammals, isopods and primates. The reconstructed history of *N*_e_ in these groups appears to correlate with LHTs or ecological variables in a way that suggests that the reconstruction is reasonable, at least in its global trends. On the other hand, the range of variation in Ne inferred across species is surprisingly narrow. This last point suggests that some of the assumptions of the model, in particular concerning the assumed absence of epistatic interactions between sites, are potentially problematic.

## 1 Introduction

Since the realization, by Zuckerkandl and Pauling (1965) that genetic sequences are informative about the evolutionary history of the species, molecular phylogenetics has developed into a mature and very active field. A broad array of models and inference methods have been developed, using DNA sequences for reconstructing the phylogenetic relationships among species (Felsenstein, 1981), for estimating divergence times (Thorne and Kishino, 2002), or for reconstructing the genetic sequences of remote ancestors (Liberles, 2007). However, genetic sequences might contain information about other aspects of the evolutionary history and, in particular, about past population-genetic regimes.

Interspecific divergence is the long-term outcome of population-genetic processes, in which point mutations at the level of individuals are then subjected to selection and genetic drift, leading to substitutions at the level of the population. As a result, the substitution patterns that can be reconstructed along phylogenies are modulated by the underlying population-genetic parameters (mutation biases, selective landscapes, effective population size), suggesting the possibility to infer the past variation of these parameters over the phylogeny. Independently, ecological properties such as phenotypic characters or life-history traits can be observed in extinct or in present-day species. Using the comparative method (Felsenstein, 1985), these traits can be reconstructed for the unobserved ancestral species. Combined together, genetic and phenotypic ancestral reconstructions can then be used to unravel the interplay between evolutionary and ecological mechanisms.

Practically, in order to disentangle mutation, selection and genetic drift, we need to classify individual substitutions into different categories, differing in the strength of mutation, selection or genetic drift. In protein-coding DNA sequences, the mutational process occurs at the nucleotide level. Assuming that synonymous mutations are selectively neutral and that selection mostly acts at the protein level, synonymous substitutions can be used to infer the patterns of mutation, without any interference contributed by selection. Then, by comparing the non-synonymous substitution rate relative to the synonymous substitution rate (the ratio *d*_*N*_*/d*_*S*_), one can estimate the global strength of selection acting on proteins. This idea was formalized using phylogenetic codon models (Muse and Gaut, 1994; Goldman and Yang, 1994). This led to a broad range of applications, either to detect proteins under adaptive selection (Kosiol *et al.*, 2008), or to measure the modulations of the strength of purifying selection between sites (Echave *et al.*, 2016), genes (Zhang and Yang, 2015), or lineages (Lartillot and Poujol, 2011).

Concerning variation in *d*_*N*_*/d*_*S*_ between lineages, and in a context mostly characterized by purifying selection, the nearly-neutral theory predicts that changes in the global strength of selection (measured as *d*_*N*_*/d*_*S*_) is related to changes in the relative strength of genetic drift, which is in turn mediated by changes in effective population size (*N*_e_) (Ohta, 1992). Mechanistically, populations with high *N*_e_ are characterized by more efficient purifying selection against mildly deleterious mutations, resulting in lower *d*_*N*_*/d*_*S*_ (Kimura, 1979; Welch *et al.*, 2008).

Codon models have been used to empirically measure such changes in the efficacy of purifying selection along phylogenies, either by allowing for different *d*_*N*_*/d*_*S*_ values in different parts of the tree (Dutheil *et al.*, 2012), or by estimating *d*_*N*_*/d*_*S*_ independently for every branch of the tree (Popadin *et al.*, 2007). Alternatively, *d*_*N*_*/d*_*S*_ can be modelled as a continuous trait, varying along the phylogeny as a stochastic process, splitting at each node of the tree into independent processes (Seo *et al.*, 2004). Once empirical estimates of the variation in *d*_*N*_*/d*_*S*_ between lineages or groups has been obtained, these can be compared to changes in *N*_e_ across lineages, so as to test the validity of the predictions of the nearly-neutral theory. Independent empirical estimation of *N*_e_ is usually done vie proxies, such as the neutral diversity within species (Galtier, 2016), or life-history traits. For instance, animal species characterized by a large body size or an extended longevity are typically expected to also have a low *N*_e_ (Romiguier *et al.*, 2014). Alternatively, a Bayesian integrative framework has been proposed (Lartillot and Poujol, 2011), extending the approach of Seo *et al.* (2004), in which the joint variation in *d_S_*, *d*_*N*_*/d*_*S*_ and in life-history traits or other proxies of *N*_e_ is modelled as a multivariate Brownian process, with a variance-covariance matrix capturing the signal of their correlated evolution.

Analyses using these approaches and these proxies of *N*_e_ have suggested a negative correlation between *d*_*N*_*/d*_*S*_ and *N*_e_ (Popadin *et al.*, 2007; Lanfear *et al.*, 2010; Lartillot and Poujol, 2011; Lartillot and Delsuc, 2012; Romiguier *et al.*, 2014; Figuet *et al.*, 2017), thus confirming the theoretical prediction of the nearly-neutral theory. However, the universality and robustness of the correlation between *d*_*N*_*/d*_*S*_ and *N*_e_ is still debated (Nabholz *et al.*, 2013; Lanfear *et al.*, 2014; Figuet *et al.*, 2016; Bolívar *et al.*, 2019), and further investigation might be required. Moreover, these analyses do not explicitly formalize the quantitative relationship between *N*_e_ and *d*_*N*_*/d*_*S*_. This relation is in principle dependent on the underlying fitness landscape (Welch *et al.*, 2008; Cherry, 1998; Goldstein, 2011), and can show complicated behavior due to non-equilibrium properties (Jones *et al.*, 2016). These questions could be addressed in the context of a mechanistic modelling approach.

As an alternative to classical *d*_*N*_*/d*_*S*_ - based codon models, mechanistic codon models explicitly introduce population genetic equations into the codon substitution process (Halpern and Bruno, 1998). Specifically, these so-called mutation-selection codon models explicitly assign a fitness parameter to each amino acid. As a result, the substitution rate between each pair of codons can be predicted, as the product of the mutation rate and the fixation probability of the new codon, which is in turn dependent on the fitness of the initial and the final codons. Since the strength of selection is typically not homogeneous along the protein sequence, and depends on the local physicochemical requirements (Echave *et al.*, 2016; Goldstein and Pollock, 2016, 2017), local changes in selective strength are usually taken into account by allowing for site-specific amino-acid fitness profiles. Site-specific amino-acid preferences are typically estimated either by penalized maximum likelihood (Tamuri and Goldstein, 2012; Tamuri *et al.*, 2014), or in a Bayesian context, using an infinite mixture based on a Dirichlet process prior (Rodrigue *et al.*, 2010; Rodrigue and Lartillot, 2014). This second approach is further considered below.

Although not directly expressed in terms of this variable, the mutation-selection formalism induces an equilibrium *d*_*N*_*/d*_*S*_, which is theoretically lower than 1, thus explicitly modelling purifying selection (Spielman and Wilke, 2015; Dos Reis, 2015). As a result, the mutation-selection codon framework proved to be a valuable null (nearly-neutral) model, against which to compare the observed *d*_*N*_*/d*_*S*_ by classical codon models, so as to test for the presence of adaptation (Rodrigue and Lartillot, 2016; Bloom, 2017).

However, these mutation-selection methods have so far assumed the strength of genetic drift, or equivalently *N*_e_, to be constant across the phylogeny. This assumption is clearly not realistic, as attested by the empirically measured variation in *d*_*N*_*/d*_*S*_ between lineages using classical codon models or, more directly, by the broad range of synonymous neutral diversity observed across species (Galtier, 2016). The impact of this assumption on the estimation of the fitness landscape across sites (Tamuri *et al.*, 2014; Rodrigue and Lartillot, 2014), or on the tests for the presence of adaptation (Rodrigue and Lartillot, 2016; Bloom, 2017) is totally unknown. Relaxing this assumption of a constant *N*_e_ is thus necessary.

Conversely, since the mutation-selection formalism explicitly incorporates *N*_e_ as a parameter of the model, extending the model so as to let *N*_e_ vary across lineages is relatively straightforward, at least conceptually. Doing this would then provide an occasion to address several important questions: do we have enough signal in empirical sequence alignments, to estimate the evolutionary history of *N*_e_ along a phylogeny? Can we more generally revisit the question of the empirical correlations between *N*_e_ and ecological life-history traits (longevity, maturity, weight, size, …), previously explored using classical *d*_*N*_*/d*_*S*_ based models, but now in the context of this mechanistic framework?

## 2 New approaches

To address these questions, here we introduce a variant of the mutation-selection codon model, in which selection is modulated along the sequence (using site-specific amino-acid profiles), while the mutation rate (*μ*), the effective population size (*N*_e_) and life-history traits are allowed to vary along the phylogeny (figure 1). Methodologically, our model is fundamentally an integration between the Bayesian non-parametric version of the Halpern and Bruno (1998) mutation-selection model (Rodrigue and Lartillot, 2014), and the molecular comparative framework modelling the joint evolution of life-history and molecular traits (Lartillot and Poujol, 2011).

**Figure 1:**
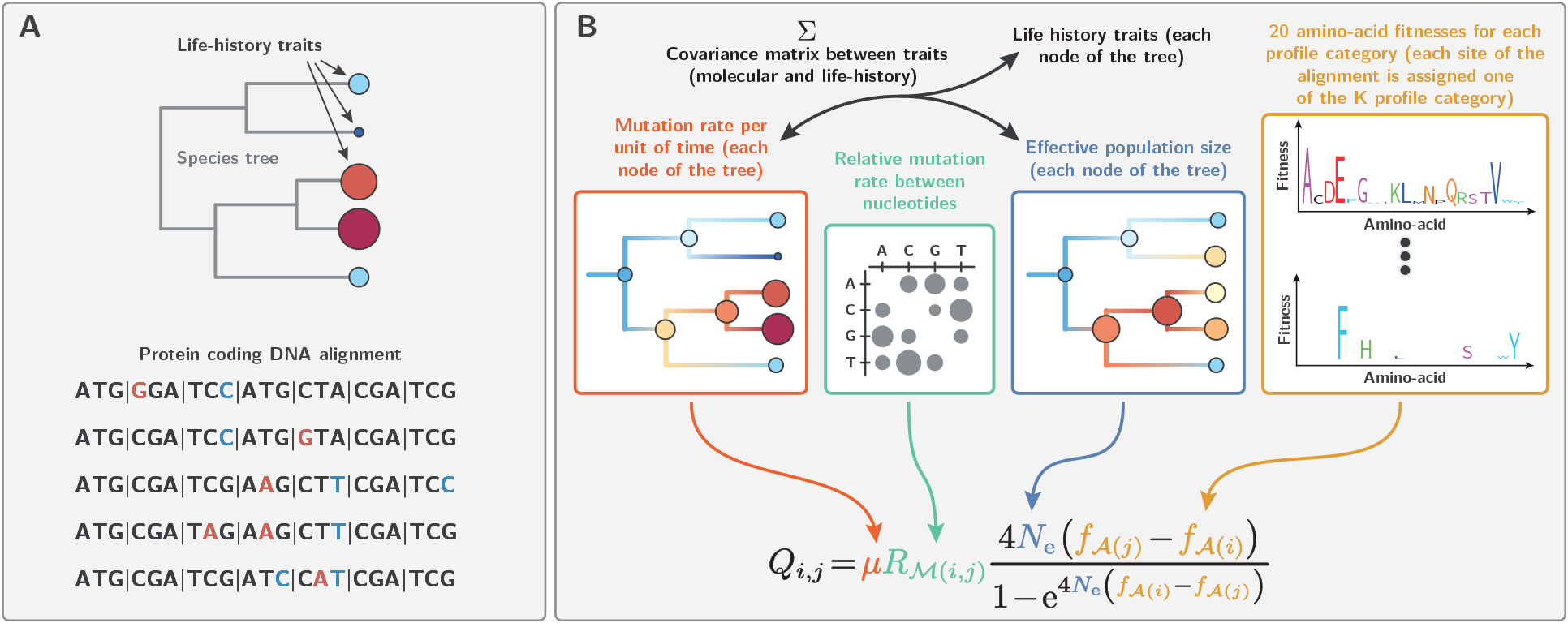
Model summary. Panel A. Our method requires a (given) rooted tree topology, an alignment of protein-coding DNA and (optionally) quantitative life-history trait for the extant species. Panel B. Relying on a codon model based on the mutation-selection formalism, assuming an auto-correlated log-Brownian process for the variation through time in effective population size (*N*_e_), mutation rate (*μ*) and life-history traits, our Bayesian inference method estimates amino-acid fitness profiles across sites, variation in mutation rate and effective population size along the tree, as well as the node ages and the nucleotide mutation rates.

Formally, the substitution rate (per unit of time) from codon *i* to *j*, denoted *Q*_*i,j*_, is equal to the total rate of mutation (per unit of time) at the level of the population (2*N*_e_*μ*_*i,j*_) multiplied by the probability of fixation of the mutation 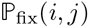:

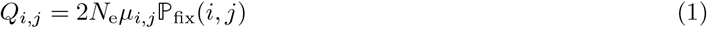

In the case of synonymous mutations, which we assumed are neutral, the probability of fixation is independent of the original and target codon, and equals 1/2*N*_e_, such that *Q*_*i,j*_ simplifies to:

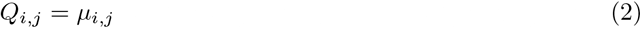

In the case of non-synonymous mutations, the probability of fixation depends on the difference in fitness between the amino acid encoded by the initial and final codons:

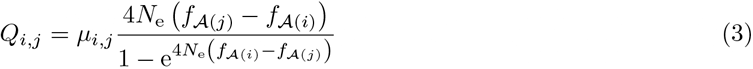

where ***f*** is a 20-dimensional vector specifying the log-fitness for each amino acid, and 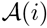 is the amino acid encoded by codon *i*.

In the model introduced here, *N*_e_ and *μ* are allowed to vary between species (across branches) as a multivariate log-Brownian process, but are assumed constant along the DNA sequence. Conversely, amino-acid fitness profiles ***f*** are considered constant along the tree but are assumed to vary across sites, being modelled as independent and identically distributed random-effects from an unknown distribution estimated using a Dirichlet process prior.

This model was implemented in a Markov chain Monte Carlo framework, allowing for joint inference of site-specific selection profiles and reconstruction of life-history traits and population-genetic regimes along the phylogeny. After validating our model and our inference framework against simulated data, we apply it to several cases of interest across metazoans (placental mammals, primates and isopods), for which some proxies of *N*_e_ are available.

## 3 Results

### 3.1 Validation using simulations

The inference framework was first tested on independently simulated multiple sequence alignments (see methods). With the aim of applying the inference method to empirical datasets, the simulation parameters were chosen so as to match an empirically relevant empirical regime. Thus, the tree topology and the branch lengths were chosen based on a tree estimated on the mammalian dataset further considered below. The other aspects of the simulation model (fitness landscape, variation in *N*_e_) were then varied along a gradient of increasing complexity, so as to test the inference framework under increasingly challenging conditions.

A first series of simulations was meant to test the soundness of our inference framework, by simulating essentially under the model used for inference, although with an independently developed software. Thus, the mutation-selection approximation was assumed to be valid, and sites were simulated under different fitness profiles empirically determined (Bloom, 2017), and finally, *N*_e_ was assumed to undergo discrete shifts at the tree nodes but otherwise to remain constant along each branch. In this context, branch lengths and branch-specific values of *N*_e_ were accurately estimated by our inference method (figure 2, panel A & D). Concerning *N*_e_, the slope of the linear regression between true and estimated branch-specific *N*_e_ is 0.794 (*r*^2^ = 0.915)

**Figure 2:**
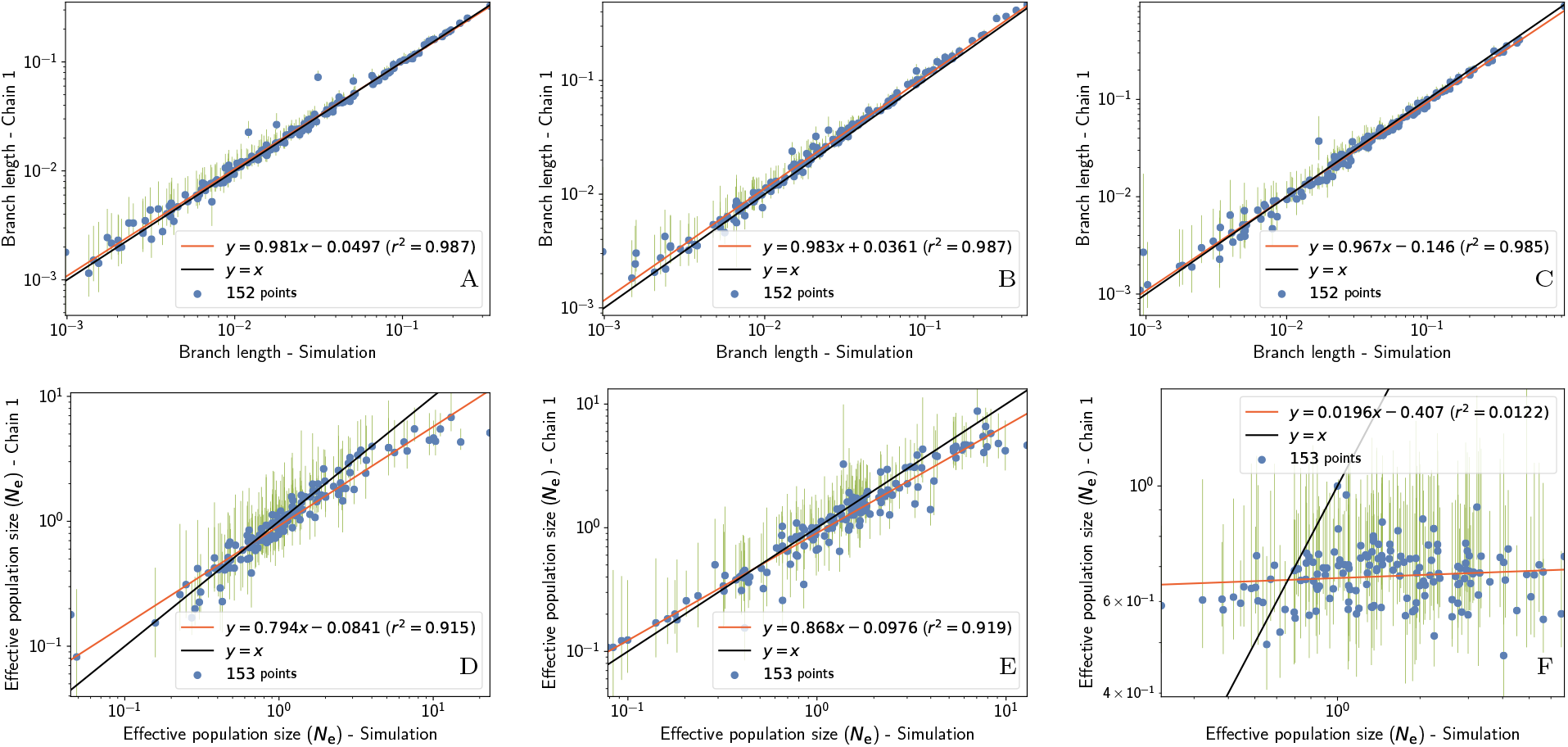
A-C: branch lengths in expected number of substitutions per site. D-F: *N*_e_ values across nodes (including the leaves) relative to *N*_e_ at the root. From left to right: simulation under the mutation-selection approximation (A,D), under a Wright-Fisher model accounting for small population size effects (5000 individuals at the root), site linkage and short term fluctuation of *N*_e_ (B,E) and accounting for site epistasis in the context of selection for protein stability. The tree root is 150 million years old, where the initial population start with a mutation rate of 1*e*^−8^ per site per generation, and generation time of 10 years. These experiments confirm that signal in the placental mammalian tree can allow to reliably infer the direction of change in *N*_e_, even if linkage disequilibrium, short term fluctuation of *N*_e_ and finite population size effects are not accounted for in the inference framework. However, the presence of epistasis between sites is a serious threat to the inference of *N*_e_.

However, the assumptions made for this first round of simulations are almost certainly violated in practice. First, *N*_e_ is expected to undergo continuous changes along the lineages of the phylogeny. Second, the diffusion approximation for the probability of fixation (equation 3) may not hold in small finite populations. Third, assuming a separate substitution process for each site is equivalent to assuming no linkage between sites (free recombination). In practice, however, there is limited recombination, at least within exons, and this could induce deviations from the mutation-selection approximation, due to Hill-Robertson effects.

The finite population was now modelled explicitly, using a Wright-Fisher simulator, tracking the frequency of each allele at the gene level and at each generation along the phylogeny. No recombination was implemented within genes. These more complex simulation settings account for small population size effects, for hitchhiking of weakly deleterious mutations during selective sweep and for background selection due to linkage disequilibrium. In addition, the effective population size *N*_e_ and the mutation rate were allowed to fluctuate continuously along the branches of the tree (changing by a small amount after each generation of the underlying Wright-Fisher process). Finally, short-term fluctuations of *N*_e_, of the order of 20% per generation, were accounted for by adding a random noise to the Brownian process describing the long-term evolution of *N*_e_. In spite of these deviations between the simulation and the inference models, branch lengths and branch-specific effective population sizes could again be robustly recovered by the inference framework (slope of 0.868, *r*^2^ = 0.919, figure 2, panel B & E).

These results are encouraging. However, they still rely on the assumption of a site-independent fitness landscape, which is equivalent to assuming no epistasis. Yet this assumption is almost certainly violated in practice (Pollock and Goldstein, 2014; Shah *et al.*, 2015). Accordingly, we implemented a more complex, site-dependent fitness landscape accounting for the selective interactions between sites induced by the 3-dimensional structure of protein. In this model, the conformational stability of the protein determines its probability of being in the folded state, which is in turn taken as a proxy for fitness (Williams *et al.*, 2006; Goldstein, 2011; Pollock *et al.*, 2012). Under this evolutionary model, and at any gven time, the fitness landscape at a particular codon site is dependent on the amino acids that are currently present at those sites that are in the vicinity of the focal site in 3D space (see supplementary). When applied to data simulated using this model, our inference framework could accurately recover the simulated branch lengths (figure 2, panel D). On the other hand, the distribution of *N*_e_ across the tree could not be accurately recovered (slope of 0.0196, *r*^2^ = 0.0122, figure 2, panel F). In fact, no meaningful variation in *N*_e_ is detected, and the little variation in *N*_e_ that is inferred shows no correlation with the true branch-specific mean *N*_e_ values. This effect can be explained by the predicted independence of *d*_*N*_*/d*_*S*_, and more generally of the scaled selection coefficients associated with non-synonymous mutations, to changes in *N*_e_ in this specific model of protein stability, as shown theoretically by Goldstein (2013).

As an alternative model of epistasis between sites, a Fisher geometric model was also considered for the simulations (see supplementary). The results under this model are intermediate between simulations without epistasis and simulations under the biophysically-inspired model considered above. More specifically, under data simulated using Fisher’s geometric model, the true and estimated branch-specific *N*_e_ are strongly correlated with each other (*r*^2^ = 0.73). On the other hand, the slope of the correlation is substantially less than 1 (0.571). In other words, the trends in *N*_e_ across the tree are correctly recovered, but the range of the variation in effective population size over the tree is substantially under-estimated. As for the branch lengths, they are again correctly estimated. In summary, our simulation experiments show that our inference framework is reliable in the absence of model mis-specification and is robust to violations concerning short-versus long-term variation in *N*_e_ or to the presence of empirically reasonable levels of Hill-Robertson interference. On the other hand, and very importantly, epistasis, which is ignored by the inference model, appears to lead to a general underestimation of the true variation in Ne, to an extent that depends on the exact epistatic model but can go as far as completely obliterating any signal about the true variation in *N*_e_ across the tree in the most extreme situations.

### 3.2 Empirical experiments

We next applied our inference framework to a series of 4 empirical datasets spanning different taxnonomic groups within metazoans. As a first empirical case, we considered a dataset of 77 placental mammals, for which complete genome sequences and information about life-history traits is available. Placental mammals offer an interesting example, for which effective population size is likely to show substantial variation across lineages. This variation in *N*_e_ is expected to covary with life-history traits (LHTs), such that large-bodied species are expected to have smaller effective population sizes, compared to small-bodied species.

For computational reasons, we restricted our analyses to small concatenates made of 18 randomly sampled alignments of orthologous genes. Since the mutation-selection model considered here assumes a mostly nearly-neutral regime, genes for which positive selection was detected using a site codon model were excluded. To assess the reproducibility of our inference and check that the signal about variation in *N*_e_ is not driven by particular genes, we analysed 4 concatenated random samples of 18 genes. The different concatenate showed similar trends in the change of *μ* (*r*^2^ = [0.92, 0.95]) and *N*_e_ (*r*^2^ = [0.51, 0.68]) between pairs of experiments (see supplementary).

The reconstructed long-term changes in effective population size (*N*_e_) is displayed in figure 3. We visually observe a global trend of increasing *N*_e_ throughout the tree around 90 and 60 My. We also observe *N*_e_ to be lower in some clades, such as Cetacea and Camelidae, while being higher in other clades, such as Rodentia and Pecora. In some cases, a decrease in *N*_e_ can be observed along an isolated branch of the tree, for example on the branches leading to the Alpaca (*Vicugna pacos*) or the cheetah (*Acinonyx jubatus*).

**Figure 3:**
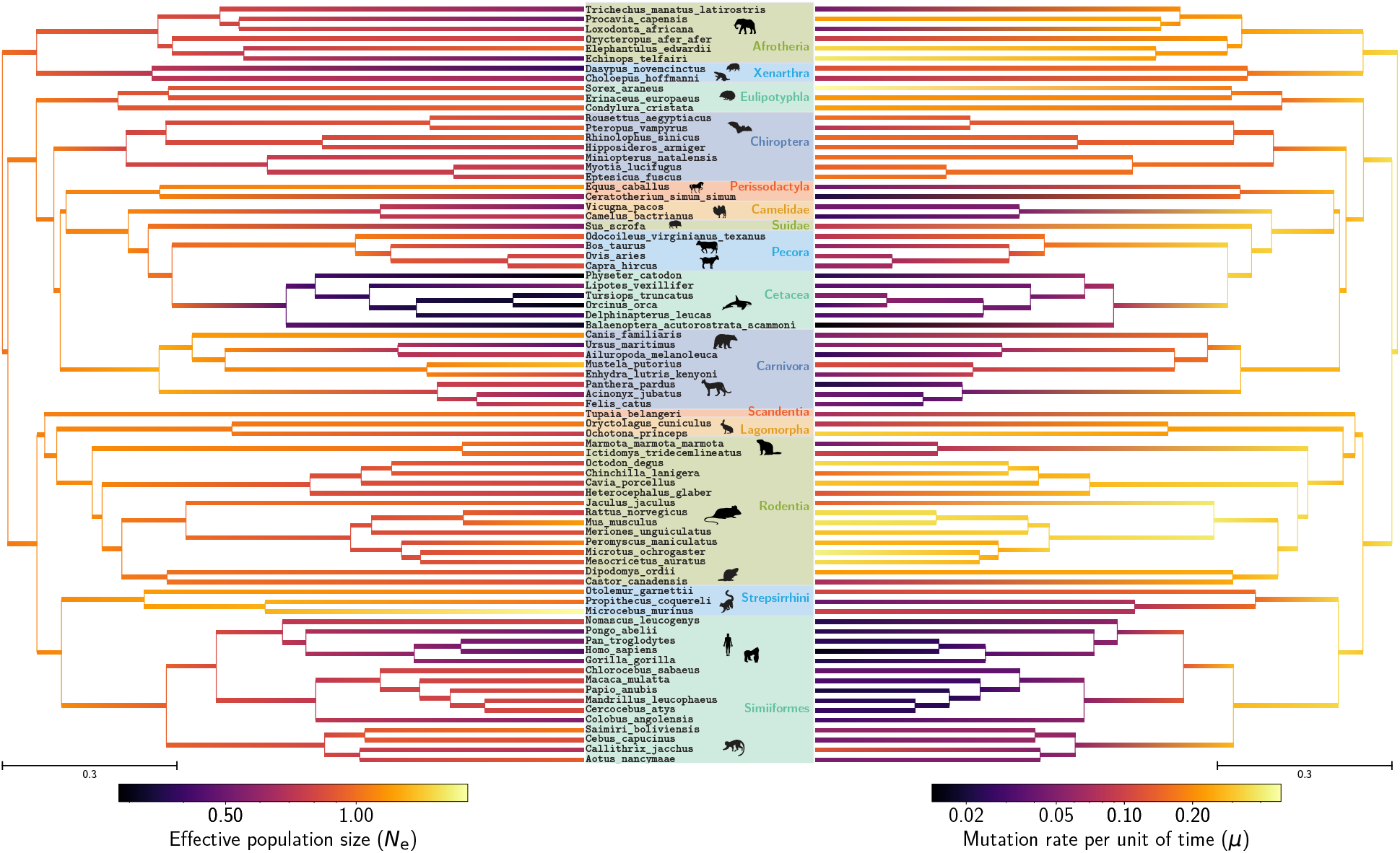
Inferred phylogenetic history of *N*_e_ (left) and *μ* (right) across placental mammals. Inference was conducted on a randomly chosen set of 18 out of 226 highly conserved CDS (¡ 1% of gaps). Only highly conserved CDS were retained such that the assumption of constant fitness landscape is not incautiously broken by protein with changing function and/or adaptive selection. *N*_e_ values are relative to the root, which is arbitrarily set to one. Mean values of MCMC (after burn-in) are obtained at each node of the tree, hence a gradient can be extrapolated along each branch. *μ* spanned almost 2 order of magnitude, and if we assume the root to be 105My old (Kumar *et al.*, 2017), the rescaled mutation rate per site per year in extant species is between 1.1*e*^−10^ and 7.8*e*^−9^. *N*_e_ at the root of the tree is arbitrarily set to 1, and all values are relative to the root, which spans at most an order of magnitude.

The estimated covariance matrix (table 1) gives a global synthetic picture about the patterns of covariation between the mutation rate per unit of time *μ*, the effective population size *N*_e_ and the three LHTs. First, the variation in *μ* across species is negatively correlated with variation in body mass, age at sexual maturity and longevity (*ρ* = [−0.84, −0.83], table 1). These correlations, which were previously reported (Lartillot and Delsuc, 2012; Nabholz *et al.*, 2013) probably reflect generation time effects (Lanfear *et al.*, 2010; Gao *et al.*, 2016). Similarly, and more interestingly in the present context, the variation in *N*_e_ between species is also negatively correlated with LHTs (*ρ* = [−0.54, −0.47], table 1). This is consistent with the expectation that small-sized and short-lived species tend to be characterized by larger effective population sizes (Romiguier *et al.*, 2014). Of note, these results mirror previous findings, based on classical codon models, showing that *d*_*N*_*/d*_*S*_ tends to be positively correlated with LHTs (Lartillot and Delsuc, 2012; Nabholz *et al.*, 2013; Figuet *et al.*, 2017). Result which was also recovered on the present dataset, using a classical *d*_*N*_*/d*_*S*_ based codon model (supplementary materials). Interestingly, the correlation of *d*_*N*_*/d*_*S*_ with LHTs is weaker than that of our inferred *N*_e_ with LHTs, as expected if the variation in *d*_*N*_*/d*_*S*_ indirectly (and imperfectly) reflects the underlying variation in Ne. Finally, *N*_e_ and *μ* are positively correlated in their variation (*ρ* = 0.44), which might simply reflect the fact that both negatively correlate with LHTs. The partial-correlation coefficients (see supplementary) between *N*_e_ and LHTs are not significantly different from 0. However, this might simply be due to the very strong correlation between the three LHTs considered here (*ρ* = [0.81, 0.85]), such that controlling for any one of them removes most of the signal contributed by the available empirical variation between species.

**Table 1:**
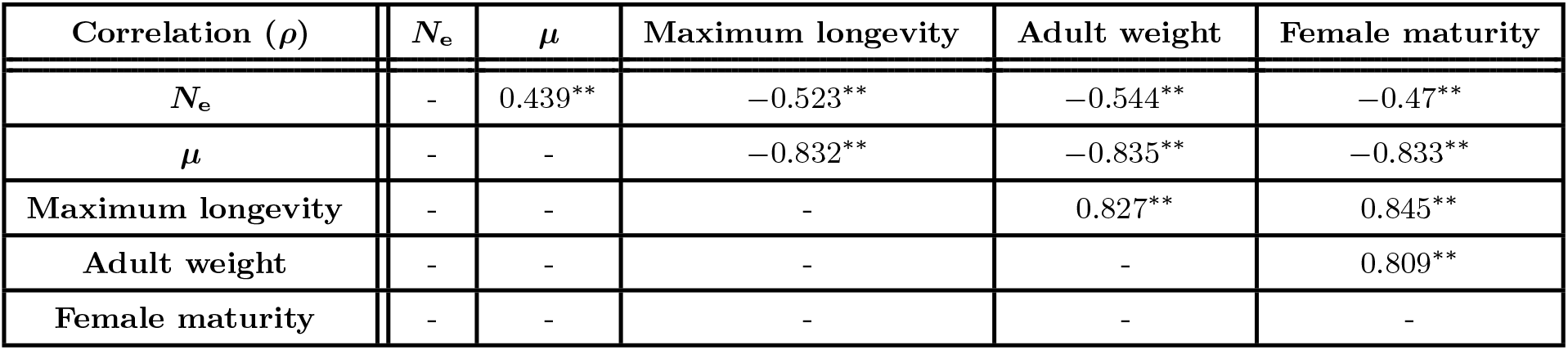
Correlation coefficient between effective population size (*N*_e_), mutation rate per site per unit of time (*μ*), and life-history traits (Maximum longevity, adult weight and female maturity). Asterisks indicate strength of support of the posterior probability to be different than 0 (pp) as \**pp* > 0.95 and \*\**pp* > 0.975. Observed correlations are compatible with the interpretation that large populations are composed of small, short-lived individuals. Moreover if the mutation rate per generation is considered constant in first approximation, the mutation rate per unit of time is positively correlated to generation rate, hence to population size.

Thus, altogether, the inferred trends in *N*_e_ across species appear to be as expected, based on considerations about life-history evolution. On the other hand, the total range of the inferred variation in *N*_e_ across the entire extant taxa is surprisingly narrow, with one order of magnitude (9.2) at most between high and low *N*_e_ (see supplementary). This almost certainly represents an underestimate of the true range of variation across placental mammals.

As another case study, we analysed a group of isopod species that have made multiple independent transitions to subterranean environments. The transition from a terrestrial to a subterranean lifestyle is typically associated with a global life-history and ecological syndrome characterized by a loss of vision, longer generation times and, most interestingly, smaller population sizes, due to a lower carrying capacity of the subterranean environment (Capderrey *et al.*, 2013). Protein coding DNA sequence alignments and qualitative life-history traits such as habitat (surface or underground), pigmentation (depigmented, partially depigmented or pigmented) and ocular structure (anophthalmia, microphthalmia, or ocular) are available for these species (Eme *et al.*, 2013; Saclier *et al.*, 2018). The assumption of a Brownian auto-correlated process for describing the changes in *N*_e_ along the tree may not be so well adapted to the present case, since the changes in *N*_e_ associated with the transition to a subterranean environment are likely to correspond to relatively sudden shifts, rather than continuous variation, and the ecological correlate (subterranean versus terrestrial) is not a quantitative trait. However, the dataset considered here contains independent transitions to a subterranean lifestyle, thus offering an opportunity to test for a potential correlation between inferred *N*_e_ variation and terrestrial versus subterranean lifestyles over the terminal branches. In our analysis across 4 concatenated random samples of 12 genes, we observe a reproducible (see supplementary) and statistically significant reduction in *N*_e_ for underground or depigmented species, or for species with visual impairment (see figure 4). Of note, the species that dit not undergo a transition to subterranean environments feature a relative *N*_e_ close to 1, meaning that *N*_e_ has not changed much along the lineages (since the root of the tree). Again, the total range of the inferred variation in *N*_e_ across the entire extant taxa is surprisingly narrow, with ratio of 3.3 at most between high and low *N*_e_ (see supplementary).

**Figure 4:**
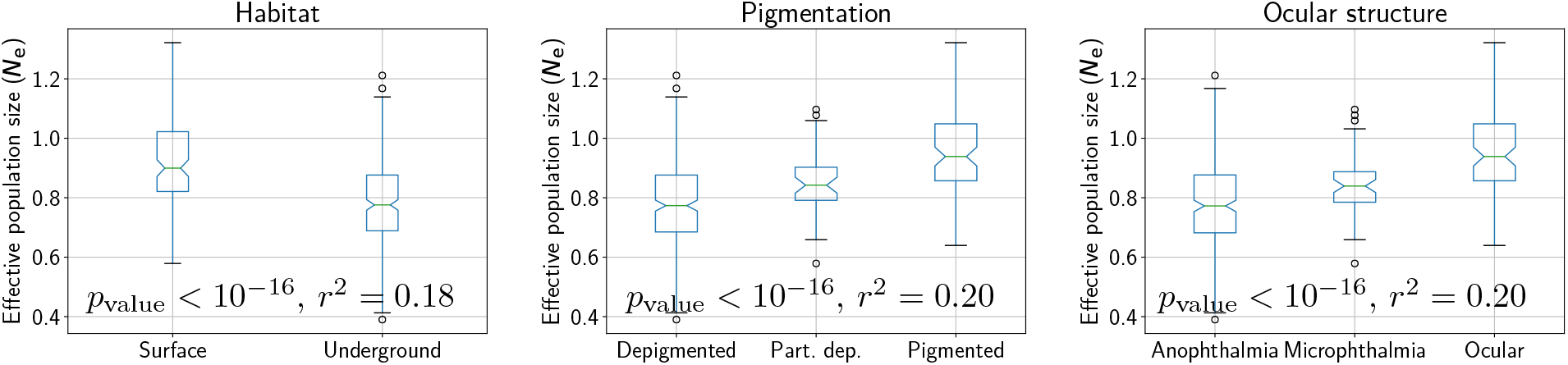
*N*_e_ estimation for extant isopods species, sorted according to their habitat (left), pigmentation (middle), and ocular structure (right). All three qualitative trait statistically correlates with changes in *N*_e_. Underground, or depigmented species, or species with visual impairment are characteristic of low *N*_e_ species.

Next, our empirical framework was also applied on a set of genes sampled across primates, taken from Perelman *et al.* (2011) and reanalysed in Brevet and Lartillot (2019). In addition to LHTs (mass, female maturity, generation time and longevity), information about nuclear synonymous diversity (*π*_*S*_) and non-synonymous over synonymous diversity (*π*_*N*_ */π*_*S*_), are available for 10 species across the dataset and are expected to correlate with *N*_e_ according to population genetics (Eyre-walker and Keightley, 2007; Galtier, 2016). However, the correlation coefficient between our inferred *N*_e_ and *π*_*S*_ or *π*_*N*_ */π*_*S*_ and LHTs are not statistically significant, nor with LHTs (see supplementary). Again, the total range of the inferred variation in *N*_e_ across the entire tree is narrow, with ratio of 6.4 at most between high and low *N*_e_. This results contrasts with the finding of Brevet and Lartillot (2019) on the same dataset based on *d*_*N*_*/d*_*S*_ - based codon models, where the estimated *N*_e_ was found to span several orders of magnitude, and correlated positively with *π*_*S*_.

## 4 Discussion

Mechanistic phylogenetic codon models express the substitution rates between codons as a function of the mutation rates at the nucleotide level, selection over amino-acid sequences and effective population size. Thus far, the development of mutation-selection models of the HB family (Rodrigue *et al.*, 2010; Tamuri and Goldstein, 2012) has mostly focused on the question of fully accounting for the fine-scale modulations of selection between amino-acids and across sites (Rodrigue *et al.*, 2010; Tamuri and Goldstein, 2012). However, the issue of the variation in the global population-genetic regime between species has received much less attention. In particular, effective population size (*N*_e_) is expected to vary substantially over the species of a given clade, yet current mutation-selection models all invariably assume *N*_e_ to be constant across the phylogeny.

Here, we have introduced an extension of the mutation-selection model that accounts for this variation. When applied to an alignment of protein coding sequences, this mechanistic model returns an estimate of the modulations of amino-acid preferences across sites. Simultaneously, it reconstructs the joint evolution of life-history traits and molecular and population-genetic parameters (mutation rate *μ* and effective population size *N*_e_) along the phylogeny, while estimating the correlation matrix between these variables, intrinsically accounting for phylogenetic inertia.

### 4.1 Reliability of the inference of the phylogenetic history of *N*_e_

The reconstructions obtained on several empirical datasets, in particular in mammals and in isopods, suggest that the method is able to correctly infer the directional trends of the changes in *N*_e_ across species. In particular, in mammals, the inferred variation in *N*_e_ correlates negatively with body size and, more generally, with life-history traits, as expected under the reasonable assumption that large-bodied mammals would tend to have smaller effective population sizes Popadin *et al.* (2007); Lartillot and Delsuc (2012); Nabholz *et al.* (2013); Figuet *et al.* (2017). Similarly, in isopods, smaller effective population sizes are inferred in subterranean species, again, as expected (Capderrey *et al.*, 2013).

However, if the trends are in right direction, the magnitude of the changes inferred across the phylogeny is surprisingly narrow and does not match independent empirical estimates of the variation in those clades. In particular, in mammals, synonymous diversity varies by a factor at least 10 between species (Galtier, 2016). In animals, the synonymous diversity roughly spans two orders of magnitude, whereas *N*_e_ varies considerably more across species, by a factor of 10^3^ (Galtier and Rousselle, 2020). For instance, effective population sizes estimated based on population genomic data are of the order of 10 000 in humans (Li and Durbin, 2011), and 100 000 in mice (Geraldes *et al.*, 2008). Thus, clearly, our approach underestimates the true variation. Different mechanisms not accounted for by the model could explain this result.

First, genetic hitchhiking, Hill-Robertson interference, and short-term fluctuations of *N*_e_ could generate this effect. However, inference conducted on alignments simulated under a Wright-Fisher model accounting for linkage and for short-term variation in *N*_e_ suggests that empirically reasonable levels of Hill-Robertson interferences are not strong enough to explain this observation, at least in the regimes explored. Second, *μ* and *N*_e_ could also be fluctuating along the genome (Gossmann *et al.*, 2011; Ellegren *et al.*, 2003; Eyre-Walker and Eyre-Walker, 2014). This assumption needs to be tested, though we expect that relaxing this assumption would not change drastically the magnitude of inferred *N*_e_ since some of this fluctuation should be absorbed by the inferred site-specific fitness profiles. Third, the DNA sequences could also be misaligned at some sites. However we observe the same magnitude of inferred *N*_e_ for different sets of genes indicating this might not be the primary reason. Fourth, the genes selected in our alignments could be under adaptive evolution, or their function could have changed. However, at least in mammals, the impact of this potential problem was minimized by the use of genes for which no positive selection was detected using standard phylogenetic codon site models.

Finally, one key assumption of the mutation-selection model that is likely to be violated in practice is the assumption of site-independence. In reality, epistasis might be prevalent in protein coding sequence evolution (Pollock and Goldstein, 2014; Shah *et al.*, 2015). Our simulations under an epistatic landscape point to epistasis being a major factor to be investigated. Indeed, *N*_e_ could not be appropriately estimated under these simulation settings, although the outcome more specifically depends on the exact model for the fitness landscape. An extreme case is obtained using a biophysically-inspired model, assuming purifying selection for conformational stability. This model was previously explored using simulations and theoretical developments Goldstein (2013), and it was shown that, under this model, *d*_*N*_*/d*_*S*_ and more generally the substitution process is virtually insensitive to *N*_e_. This is confirmed by our experiments, showing that the mutation-selection approach explored here cannot infer the true variation in *N*_e_ under this model.

A less extreme outcome is obtained under an alternative model also implementing epistatic interactions between sites via Fisher’s geometric model (Tenaillon, 2014; Blanquart and Bataillon, 2016). Interestingly, under this model, our inference framework is able to infer the correct trends of *N*_e_, although with a substantially underestimated range of inferred variation, thus mirroring the results obtained on placental mammals. Of note, these results do not necessarily imply that models based on biophysics are empirically less relevant than Fisher’s geometric model. Instead, they might just betray that the response of the substitution process to changes in *N*_e_ may be sensitive to the exact quantitative details of the underlying fitness landscape. More work is probably needed here to characterize these exact conditions. Nevertheless, our simulation experiments suggest a global pattern: epistatic interactions induce a buffering of the response of the substitution process to changes in *N*_e_. The meaningful correlation patterns observed with LHTs in the case of placental mammals suggest that this buffering is not complete. Nevertheless, ignoring epistatic interactions at the inference level appears to result in a substantial underestimation of the range over which *N*_e_ varies across species.

Interestingly, the magnitude of the inferred range of *N*_e_ variation is similar for the placental and the primate datasets (with a 9-fold and 6-fold variation in mammals and primates, respectively), whereas one would have expected a much larger range of variation over the broader phylogenetic scale of placental mammals, compared to primates. An explanation could be that the effects of epistasis are more apparent at longer time-scales. Indeed, the total number of substitutions from root to leaves is greater, and as a result, the local environment, and therefore the fitness landscape at the level of each site, has been less stable across the phylogeny.

Although modelling epistasis in an inference framework is a complex biological, mathematical and computational problem, our work points to a potential signal of epistasis that could be retrieved in a phylogenetic context. More specifically, since the slope of the response of the substitution process to changes in *N*_e_ appears to be informative about the epistatic regime, then, conversely, by relying on independent estimates of *N*_e_ (e.g. using polymorphism), this effect could be used to leverage a quantitative estimate of the statistical distribution of epistatic effects.

Other methods have recently been developed to reconstruct phylogenetic changes in *N*_e_. For example, a method recently developed uses polymorphism and generation time for some present-day species to reconstruct *N*_e_ along the phylogeny, based on a classical (*d*_*N*_*/d*_*S*_ - based) codon model (Brevet and Lartillot, 2019). This method implicitly relies on a nearly-neutral model, assuming a fixed and gamma-shaped distribution of fitness effects across non-synonymous mutations. The approach is calibrated using fossils, and as a result, returns estimates of the absolute value of *N*_e_ and of its phylogenetic variation. Here, in contrast, our method requires neither generation times nor polymorphism data, and the fitness effects are not constrained to a specific distribution. On the other hand, the inferred effective population sizes are only relative. In addition, the empirical fitting of the model requires more computing resources.

### 4.2 Potential applications and future developments

Apart from reconstructing the phylogenetic history of *N*_e_ and investigating its causes and covariates, another potentially interesting application of our approach is in detecting adaptation. In this direction, mutation-selection models represent a useful null nearly-neutral model, explicitly modelling the background of purifying selection acting over protein coding genes. Adaptation can then be detected by measuring the deviation from this null model (Rodrigue and Lartillot, 2016; Bloom, 2017).

However, by assuming a constant *N*_e_ along a phylogeny, the statistical power of this approach to detect sites under adaptive evolution may not be optimal. In particular, the site-specific fitness profiles inferred by the model are averaged along the phylogeny and are seemingly more diffuse than those estimated profiles under our present framework (see supplementary materials). Thus, our method should provide a better null model of purifying selection against which to test for the presence of adaptive evolution.

This approach can be further extended in other directions. First, currently, our model also assumes no selection on codon usage. In the case of primates or placental mammals, this assumption is probably reasonable (Yang and Nielsen, 2008), although it is more questionable for other groups, in particular Drosophila (Duret and Mouchiroud, 1999; Plotkin and Kudla, 2011). In principle, this assumption can be relaxed by implementing selective codon preferences that are shared across all sites. Such an implementation would provide the advantage of estimating codon usage biases, while simultaneously accounting for its confounding effect when estimating selection on amino-acids and inter-specific variation in *N*_e_.

Second, the Bayesian analysis conducted here was based on relatively small alignments (20 000 sites at most), and with strong limits on the parametrization of the underlying mixture model (allowing for at most 50 distinct profile categories). Profiling of the program (not shown) shows that the number of components of the profile mixture is the limiting step of the computation. Yet, a larger number of components might be required, in order to achieve more accurate inference of the site-specific profiles. One possible development, leading to statistically more stable genome-wide estimates of *N*_e_, would be to develop a multi-gene parallelized version of the model, in which each coding sequence would have its own mixture model, and would run on a separate thread, while the history of *N*_e_ would be shared by all computing processes.

Finally, estimating *N*_e_ in a mutation-selection phylogenetic model relies on the relation between *N*_e_ and the relative strength of drift, in a context where, ultimately, the signal about the intensity of drift comes from the relative rate of non-synonymous substitutions. However, this purely phylogenetic approach does not leverage a second aspect of *N*_e_ at the population level, namely, the fact that *N*_e_ also determines the levels of neutral genetic diversity that can be maintained (*π* = 4*N*_e_*u*, where *u* is the mutation rate per generation). Hence, neutral diversity yields an independent empirical estimate of *N*_e_. In principle, our mechanistic model could be extended so as to incorporate polymorphism data within species at the tips of the phylogeny. A similar method has been previously pioneered in the case of 3 species and using a distribution of fitness effect(Wilson *et al.*, 2011). More generally, the nearly-neutral theory of evolution defines a long-term *N*_e_, which might be different from the short-term definition of *N*_e_ (Platt *et al.*, 2018). Thus we could ask if empirical independent estimations of *N*_e_ from within species (based on genetic diversity) and between species (based on the substitution process) are congruent, and if not, what are the mechanisms responsible for this discrepancy.

Notwithstanding theoretical considerations on the nearly-neutral theory of evolution, empirical clues about the long-term trends in the modulations of the intensity of genetic drift opens up a large diversity of ecological and evolutionary questions. Spatial and temporal changes of genetic drift along ecological niches and events can now be investigated, so as to disentangle the underlying evolutionary and ecological pressures.

## 5 Materials and Methods

In the model presented here, *N*_e_ and *μ* (and quantitative traits) are allowed to vary between species (across branches) as a multivariate log-Brownian process, but assumed constant along the DNA sequence. Conversely, amino-acid fitness profiles are assumed to vary across sites, but are considered constant along the tree. The model makes several assumptions about the evolutionary process generating the observed alignment. First, the species tree topology is supposed to be known, and each gene should match the species tree, meaning genes are strict orthologs (no paralogs and no horizontal transfers). Second, there is no epistasis (interaction between sites), such that any position of the sequence has its own independent evolutionary process and a substitution at one position does not affect the substitution process at other positions. Third, from a population genetics perspective, we assumed sites of the protein to be unlinked, or equivalently the mutation rate is low enough such that there is no Hill-Robertson interference nor genetic hitchhiking. Fourth, polymorphism is ignored in extant species.

The parameterization of the models is described as a Bayesian hierarchical model, including the prior distributions and the parameters of the model. This hierarchical model is formally represented as directed acyclic graph, depicted in figure 5.

**Figure 5:**
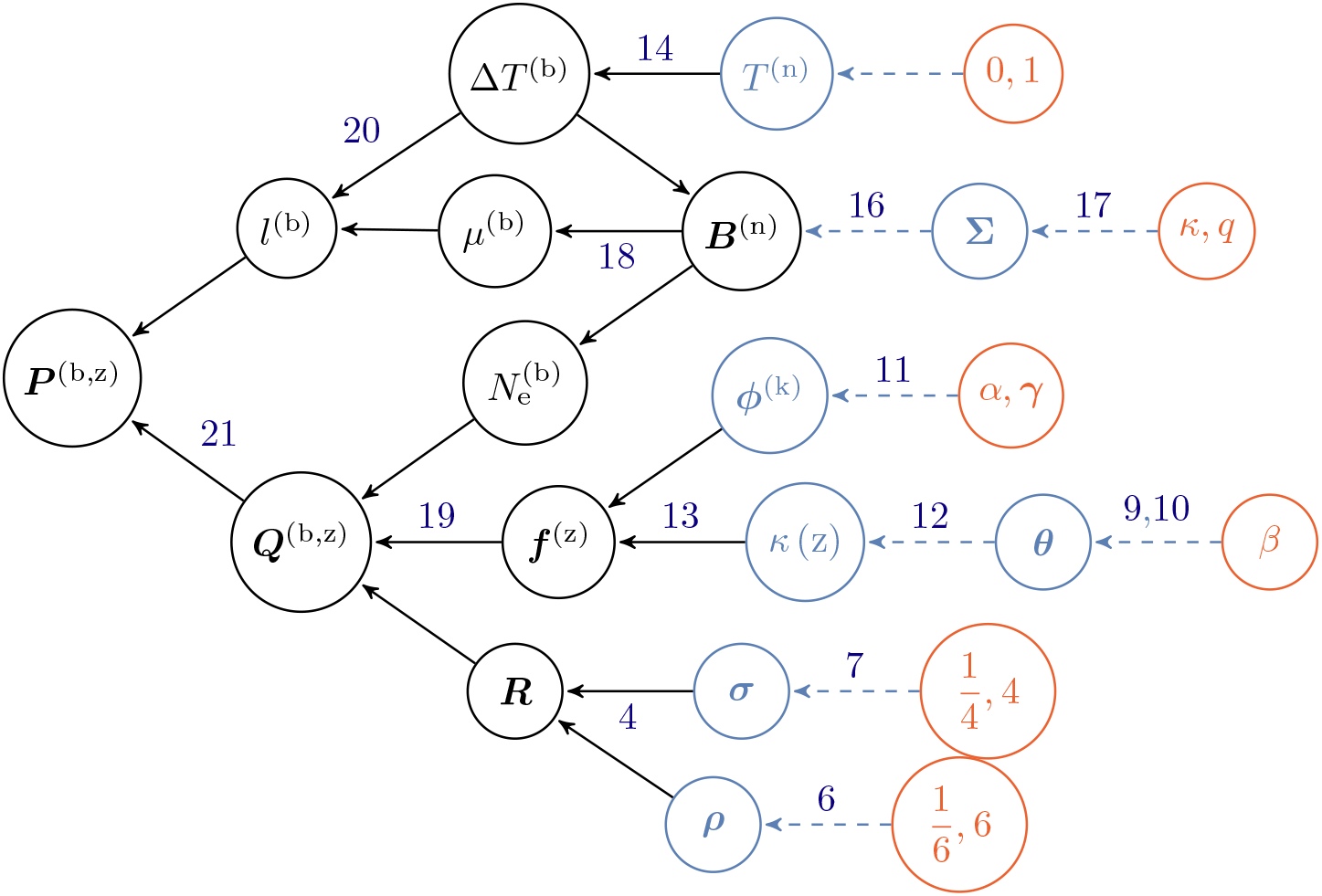
Directed acyclic graph (DAG) of dependencies between variables. Nodes of the directed acyclic graph are the variables, and edges are the functions. Hyper-parameters are depicted in 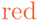 circles, random variables in 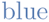 circles, and transformed variables in black. 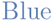 dashed line denotes a drawing from a random distribution, and black solid lines denote a function. For a given node, all the nodes pointing toward him (upstream) are its dependencies which determines its distribution. The other way around, following the arrows in the DAG (downstream), simple prior distributions are combined together to form more complex joint prior distribution which ultimately defines the prior distribution of the model.

### 5.1 Nucleotide mutation rates

The generalized time-reversible nucleotide mutation rate matrix ***R*** is a function of the nucleotide frequencies ***σ*** and the symmetric exchangeability rates ***ρ*** (Tavaré, 1986). ***σ*** = (*σ*_*A*_, *σ*_*C*_, *σ*_*G*_, *σ*_*T*_) is the equilibrium base fre-quency vector, giving the frequency at which each base occurs at each site. ***ρ*** = (*ρ*_*AC*_, *ρ*_*AG*_, *ρ*_*AT*_, *ρ*_*CG*_, *ρ*_*CT*_, *ρ*_*GT*_) is the vector of exchangeabilities between nucleotides. Altogether, the rate matrix is:

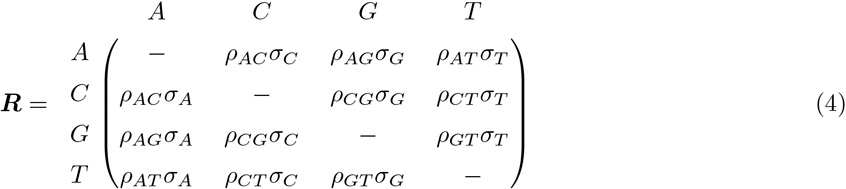

By definition, the sum of the entries in each row of the nucleotide rate matrix ***R*** is equal to 0, giving the diagonal entries:

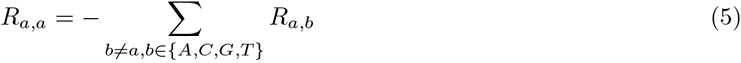

The prior on the exchangeabilities ***ρ*** is a uniform Dirichlet distribution of dimension 6:

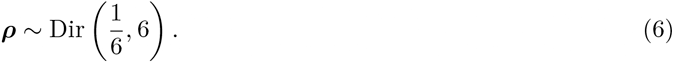

The prior on the equilibrium base frequencies ***σ*** is a uniform Dirichlet distribution of dimension 4:

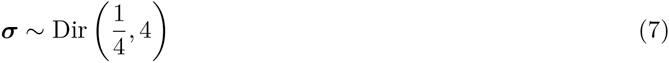

The general time-reversible nucleotide matrix is normalized such that the total flow equals to 1:

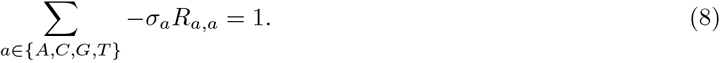

### 5.2 Site-dependent selection

Site-specific amino-acid fitness profiles are assumed i.i.d. from a mixture model, itself endowed with a truncated Dirichlet process prior. Specifically, the mixture has K components (K = 50 by default). The prior on component weights (***θ***) is modeled using a stick-breaking process, truncated at K and of parameter *β*:

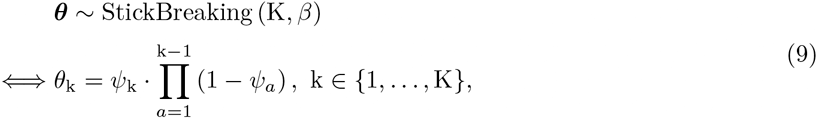

where *ψ*_k_ are i.i.d. from a beta distribution

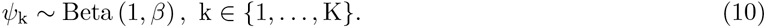

Of note, the weights decrease geometrically in expectation, at rate *β*, such that lower values of *β* induce more heterogeneous distributions of weights.

Each component of the mixture defines a 20-dimensional fitness profile ***ϕ***^(k)^ (summing to 1), for k ∈ {1,…, K}. These fitness profiles are i.i.d. from a Dirichlet of center ***γ*** and concentration *α*:

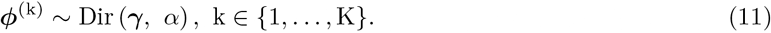

Site allocations to the mixture components *κ* (z) ∈ {1,…, K}, for z ∈ {1,…, Z} running over the Z sites of the alignment, are i.i.d. multinomial of parameter ***θ***:

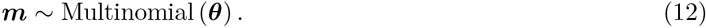

For a given parameter configuration for the mixture, the Malthusian fitness selection coefficients ***f*** ^(z)^ at site z, are obtained by taking the logarithm of the fitness profile assigned to this site:

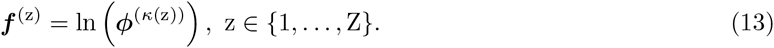

### 5.3 Dated tree

The topology of the rooted phylogenetic tree is supposed to be known and is not estimated by the model. The model estimates the dates at which branches split, thus the dated tree requires *P* − 2 internal node ages that are free parameters, where *P* is the number of extant taxa (leaves of the tree). By definition, leaf ages are all set to 0. The root age is set arbitrarily to 1, but if fossils data are also available the dated tree can be rescaled into absolute time using cross-multiplication. A uniform prior is assumed over internal node ages *T* ^(n)^, n ∈ {*P* + 1,…, 2*P* − 2}.

The duration Δ*T* ^(b)^ represented by a given branch b, for b ∈ {1,…, 2*P* − 2} is defined as the difference in ages between the oldest node at the tip of the branch *T* ^(b*↑*)^, and the youngest node *T* ^(b*↓*)^:

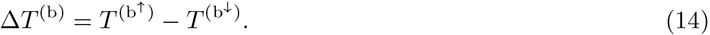

### 5.4 Branch dependent traits

The effective population size *N*_e_ and mutation rate per unit of time *μ* are assumed to evolve along the phylogeny, and to be correlated. If quantitative life-history traits (LHTs) are also available for some nodes of the tree (leaves and/or internal nodes), they are also assumed to evolve along the phylogeny and to be correlated between them, and with *N*_e_ and *μ*. The total number of traits is noted *L*, when counting *N*_e_, *μ* and all user-defined LHT (denoted ***X***). Their variation through time is modelled by an *L*-dimensional log-Brownian process ***B***. By convention, the first component of the log-brownian corresponds to *N*_e_, and the second component to *μ*. Thus:

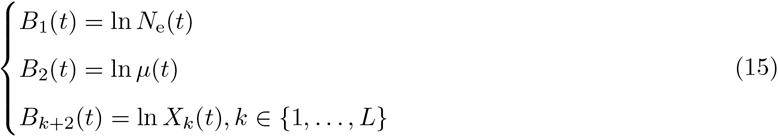

The effective population size at the root is set to 1 for identifiability of the fitness profiles.

Along a branch b ∈ {1,…, 2*P* − 2} of the tree, a log-Brownian process starts at the oldest node at the tip of the branch (b^↑^), and ends at the youngest node (b^↓^). The rate of change of the log-Brownian process per unit of time is constant and determined by the positive semi-definite and symmetric covariance matrix **Σ**. Thus the distribution at node b^↓^ of ***B***^(b*↓*)^ is multivariate Gaussian, with mean equals to the Brownian process sampled at the oldest node ***B***^(b*↑*)^, and variance Δ*T* ^(b)^**Σ**:

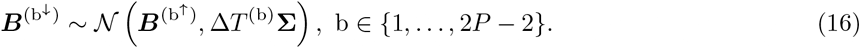

The Brownian process at the root of the tree is uniformly distributed, except for the first component fixed to 0 for identifiability (see above). The prior on the covariance matrix is an inverse Wishart distribution, parameterized by *κ* = 1 and with *q* = *L* + 1 degrees of freedom:

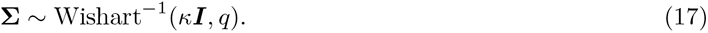

We are interested in approximating the expected substitution rates between codons over the branch. Ideally, under the Brownian process just described, the rates of substitution between codons are continuously changing through time. Also, even conditional on the value of *N*_e_ at both ends, the Brownian path along the branch entails a random component, leading to complicated integral expressions for substitution rates (Horvilleur and Lartillot, 2014). Here, a branchwise approximation is used (Lartillot and Poujol, 2011), which consists of first deriving an approximation for the mean *N*_e_ along the branch, conditional on the values of *N*_e_ at both ends, and then using this mean branchwise *N*_e_ to define the codon substitution rates.

In the case of log-Brownian process, the most likely path (or geodesic) from ***B***^(b*↑*)^ to ***B***^(b*↓*)^ is the straight line, and therefore, it would make sense to take the mean value of 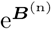 along this geodesic. We then have 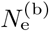 and *μ*^(b)^ for each branch b ∈ {1,…, 2*P* − 2} of the tree:

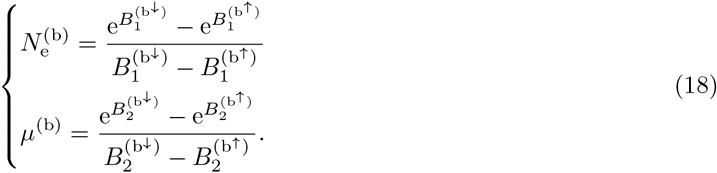

### 5.5 Codon substitution rates

The mutation rate between codons *i* and *j*, denoted *μ*_*i,j*_ depends on the underlying nucleotide change between the codons. First, if codons *i* and *j* are not nearest-neighbours, *μ*_*i,j*_ is equal to 0. Second, if codons *i* and *j* are only one mutation away, 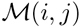 denotes the nucleotide change (e.g. 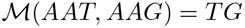), and *μ*_*i,j*_ is given by the underlying nucleotide relative rate 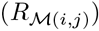 scaled by the mutation rate per time (*μ*). Technically, the 4-dimensional nucleotide relative rate matrix (***R***) is normalized such that we expect 1 substitution per unit of time, hence the scaling by *μ*.

For a given branch b and a given site z, the codon substitution rate (per unit of time) matrix ***Q***^(b,z)^ is given by:

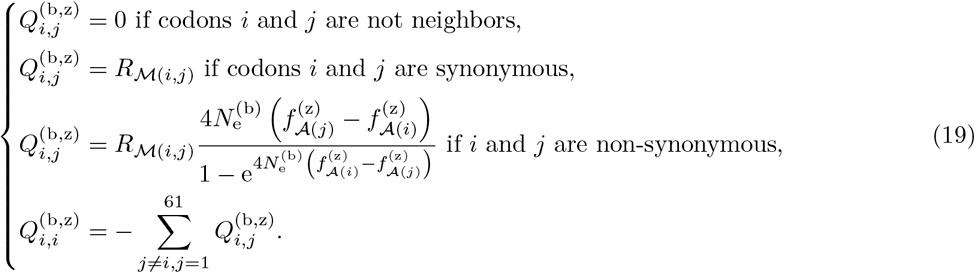

We see from this equation that, *f* and *N*_e_ are confounded, such that increasing the effective population size while decreasing the fitnesses by the same factor leads to the same substitution rate.

The branch lengths *l*^(b)^ are defined as the expected number of neutral substitutions per DNA site along a branch:

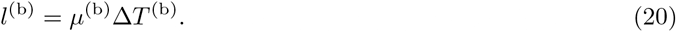

Together, the probability of transition between codons for a given branch b and site z is:

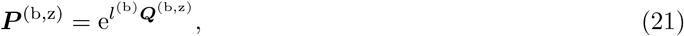

which are the matrices necessary to compute the likelihood of the data (*D*) given the parameters of the model using the pruning algorithm.

### 5.6 Bayesian implementation

Bayesian inference was conducted using Markov Chain Monte Carlo (MCMC). Most phylogenetic MCMC samplers target the distribution over the model parameters given the sequence alignment, which means that they have to repeatedly invoke the pruning algorithm to recalculate the likelihood which is most often the limiting step of the MCMC. An alternative, which is used here, is to do the MCMC conditionally on the detailed substitution history 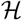, thus doing the MCMC over the augmented configuration 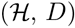, under the target distribution obtained by combining the mapping-based likelihood with the prior over model parameters.

The key idea that makes this strategy efficient is that the mapping-based likelihood depends on compact summary statistics of 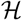, leading to very fast evaluation of the likelihood. On the other hand, this requires to implement more complex MCMC procedures that have to alternate between:

1. sampling 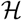 conditionally on the data and the current parameter configuration.
2. re-sampling the parameters conditionally on 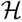.

To implement the mapping-based MCMC sampling strategy, we first sample the detailed substitution history 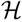 for all sites along the tree. Several methods exist for doing this (Nielsen, 2002; Rodrigue *et al.*, 2008), which are used here in combination (first trying the accept-reject method of Nielsen, then switching to the uniformization approach of Rodrigue *et al* if the first round has failed).

Then, we write down the probability of 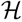 given the parameters, and finally, we collect all factors that depend on some parameter of interest and make some simplifications. This ultimately leads to relatively compact sufficient statistics (see supplementary) allowing for fast numerical evaluation of the likelihood (Irvahn and Minin, 2014; Davydov *et al.*, 2016). As an example, making an MCMC move on the *N*_e_ at a given node of the tree is faster since only the mapping-based likelihood (using path sufficient statistics) at the neighbouring branches of the node is necessary, instead of computing the likelihood for the entire tree.

Markov chain Monte Carlo (MCMC) are run for 4000 points and the first 1000 points are discarded as burn-in. Convergence is then assessed (see supplementary) by comparing two independent chains, checking that both site-specific fitness and branch *N*_e_ have the same posterior mean.

### 5.7 Correlation between traits

The correlation between trait *a* and trait *b* ∈ {1*,…, L*} can be obtained from the covariance matrix Σ:

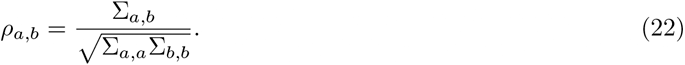

This correlation coefficient is then averaged over the posterior distribution, and statistical support is assessed based on the posterior probability of having a positive (or negative) value for the coefficient.

### 5.8 Simulations

To test the robustness of the model, four parameterized simulators were developed: `SimuDiv, SimuPoly, SimuFold & SimuGeo`. All four simulators use a log-Brownian multivariate process to model the changes in the mutation rate per generation, the generation time and *N*_e_ along the lineages. `SimuDiv, SimuFold & SimuGeo` all simulate point substitutions along the phylogenetic tree. The simulator starts from an initial sequence at equilibrium. The change in fitness is computed for all possible mutant, hence computing all strictly positive substitution rates. At each point, the next substitution is chosen proportional to these rates using in Gillespie’s algorithm (Gillespie, 1977). At each node, the process is split, and finally stopped at the leaves of the tree. `SimuPoly` simulates explicitly each generation along the phylogeny under a Wright-Fisher population, consisting of three steps: mutation, selection and genetic drift of currently segregating alleles. Mutations are drawn randomly based on mutation rates. Drift is induced by the multinomial resampling of the currently segregating alleles. We assume that the DNA sequence is composed of exons, with no linkage between exons, and total linkage of sites within an exon. Moreover, in `SimuPoly`, the instant value of log-*N*_e_ can also be modelled as a sum of a log-Brownian process and an Ornstein-Uhlenbeck process. The log-Brownian motion accounts for long-term fluctuations, while the Ornstein-Uhlenbeck introduces short-term fluctuations. In `SimuDiv` and `SimuPoly`, each codon site contributes independently to the fitness depending on the encoded amino acids, through site-specific amino-acid fitness profiles experimentally determined (Bloom, 2017). In `SimuFold`, the fitness of a sequence is computed as the probability of the protein to be in the folded state. `SimuFold` is a `C++` adaptation of a Java code previously published (Goldstein and Pollock, 2016, 2017), where we also allow for changes in *N*_e_ and *μ* along a phylogenetic tree. Supplementary materials describe the models in more details, as well as performance of the inference model against them.

### 5.9 Empirical data

For placental mammals, alignments were extracted from OrthoMam database (Ranwez *et al.*, 2007; Scornavacca *et al.*, 2019). Only highly conserved coding sequences are kept for the analysis, representing 226 CDS with ≤ 1% of gaps in the alignment. Life-history traits (LHTs) for longevity, age at maturity and weight were obtained from AnAge database (De Magalhaãs and Costa, 2009; Tacutu *et al.*, 2012). We focused our analysis on 77 taxa for which information is available for at least one LHT.

## 6 Reproducibility - Supplementary Materials

The simulators written in `C++` are publicly available under MIT license at https://github.com/ ThibaultLatrille/SimuEvol. The Bayesian inference model, written in `C++` in the component based (Lanore, 2019) software `BayesCode`, is publicly available at https://github.com/ThibaultLatrille/bayescode. Supplementary materials and figures are available in appendix supplementary materials. The scripts and instructions necessary to reproduce the simulated and empirical experiments are available at https://github.com/ThibaultLatrille/MutationSelectionDrift.

## 7 Author contributions

TL gathered and formatted the data, developed the new models in `BayesCode` and `SimuEvol` and conducted all analyses, in the context of a PhD work (Ecole Normale Superieure de Lyon). VL restructured and refactored the code sustaining the branch and site heterogeneous Bayesian Monte Carlo in `BayesCode`. TL and NL both contributed to the writing of the manuscript.

## 8 Acknowledgements

We wish to thank Tristan Lefébure for sharing the isopods phylogeny, alignments and life-history traits. We thank Philippe Veber for insightful discussion on mutation-selection models and software development. We gratefully also acknowledge the help of Nicolas Rodrigue, Laurent Gueguen, Benoit Nahbolz and Laurent Duret for their advice and review concerning this manuscript. This work was performed using the computing facilities of the CC LBBE/PRABI. Funding: French National Research Agency, Grant ANR-15-CE12-0010-01 / DASIRE.

## Notes

### Competing Interest Statement

The authors have declared no competing interest.

