## Supplementary materials for "Inferring long-term effective population size with Mutation-Selection models"

#### Contents


1 Summary statistics 1
1.1 Partial correlation coefficient 1
1.2 Fitness profile entropy 1

2 Simulations 2
2.1 Site-specific fitness profiles (`SimuDiv`) 2
2.2 Wright-Fisher with polymorphism (`SimuPoly`) 4
2.3 Fisher geometric landscape (`SimuGeo`) 5
2.4 Protein folding probability (`SimuFold`) 7

3 Empirical data in mammals 8
3.1 Chain convergence 8
3.2 Traits estimation & correlation (replicate 1, chain 1) 9
3.3 Repeatability of experiments 14
3.4 Amino-acid preferences entropy 19
3.5 Traits estimation with branch $\omega$ (replicate 1, chain 1) 19

4 Empirical data in Isopods 21
4.1 Traits estimation (replicate 1, chain 1) 21
4.2 Repeatability of experiments 23

5 Empirical data in Primates 29
5.1 Chain convergence 29
5.2 Traits estimation (chain 1) 29
5.3 Amino-acid preferences entropy 38
5.4 Traits estimation with branch $\omega$ (chain 1) 38

6 Sufficient statistics 41
6.1 Path sufficient statistics 41
6.2 Length sufficient statistics 42
6.3 Scatter sufficient statistics 42


#### 1 Summary statistics

##### 1.1 Partial correlation coefficient

The correlation coefficient $\rho_{a,b}$ give the total regression between two variables. Partial-correlation coefficient account for the entire covariance matrix, and measure the correlation between 2 traits, knowing the values of all the other traits:

$$\rho_{a,b|c\in\{1,\dots,L\}\setminus\{a,b\}} = -\frac{\Omega_{a,b}}{\sqrt{\Omega_{a,a}\Omega_{b,b}}}, \tag{1}$$

where the precision matrix $\mathbf{\Omega}$ is the inverse of the covariance matrix:

$$\mathbf{\Omega} = \mathbf{\Sigma}^{-1} \tag{2}$$

##### 1.2 Fitness profile entropy

For a category k, the Shannon's entropy ($\Omega$) of the fitness profile ($\boldsymbol{\phi}$) is defined as:

$$\Omega^{(\mathrm{k})} = -\sum_{\mathrm{a}=1}^{20} \phi_{\mathrm{a}}^{(\mathrm{k})} \ln\left(\phi_{\mathrm{a}}^{(\mathrm{k})}\right) \tag{3}$$

The Shannon's entropy measures the flatness of the fitness profile, with a value of 0 corresponding to a single peak fitness landscape (only one amino acid is present), and a value of $log(20) \simeq 3$ corresponding to a neutral landscape, where each amino acid has the same fitness.

The Shannon's entropy can be averaged over all sites as:

$$\langle \Omega \rangle = \frac{1}{\mathrm{Z}} \sum_{\mathrm{z}=1}^{\mathrm{Z}} \Omega^{\kappa(\mathrm{z})} \tag{4}$$

### 2 Simulations

#### 2.1 Site-specific fitness profiles (`SimuDiv`)

For simulations under a site-independent fitness landscape, with site-specific fitness profiles, the protein log-fitness is computed as the sum of amino-acid log-fitness coefficients along the sequence. In this model, each codon site z has its own fitness profile, denoted $\boldsymbol{\phi}^{(\mathrm{z})} = \{\phi_{\mathrm{a}}^{(\mathrm{z})},\ 1 \leq \mathrm{a} \leq 20\}$, a vector of 20 amino-acid scaled (Wrightian) fitness coefficients. Since $\mathbb{S}[\mathrm{z}]$ is the codon at site z, the encoded amino acid is $\mathcal{A}(\mathbb{S}[\mathrm{z}])$, hence the fitness at site z is $\boldsymbol{\phi}^{(\mathrm{z})}_{\mathcal{A}(\mathbb{S}[\mathrm{z}])}$. Altogether, the selection coefficient of the mutant $\mathbb{S}'$ is:

$$s\left(\mathbb{S}, \mathbb{S}'\right) = \sum_{\mathrm{z}=1}^{\mathrm{Z}} \ln \left( \frac{\boldsymbol{\phi}^{(\mathrm{z})}_{\mathcal{A}(\mathbb{S}'[\mathrm{z}])}}{\boldsymbol{\phi}^{(\mathrm{z})}_{\mathcal{A}(\mathbb{S}[\mathrm{z}])}} \right), \tag{5}$$

The fitness vectors $\boldsymbol{\phi}^{(\mathrm{z})}$ used in this study are extracted from Bloom (2017). They were experimentally determined by deep mutational scanning.

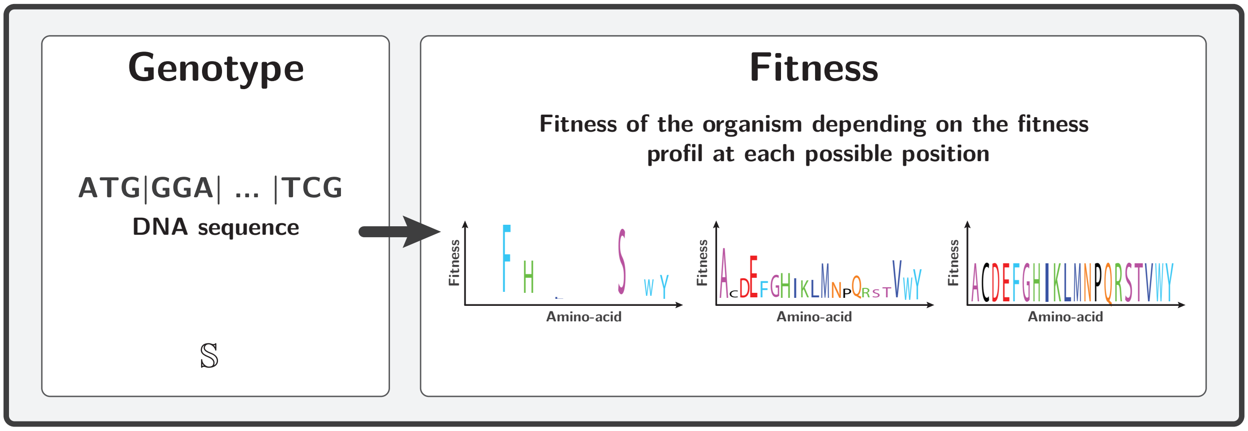


The next change in the protein coding DNA and the time to next the event is chosen using Gillespie's algorithm (Gillespie, 1977), according to the rates of substitution between codons:

$$Q_{i,j} = \mu_{i,j} \frac{4N_{\mathrm{e}} s\left(\mathbb{S}^t, \mathbb{S}^{t+1}\right)}{1 - \mathrm{e}^{-4N_{\mathrm{e}} s\left(\mathbb{S}^t, \mathbb{S}^{t+1}\right)}}, \tag{6}$$

where $Q_{i,j} = \mu_{i,j}$ in the case of synonymous substitutions.

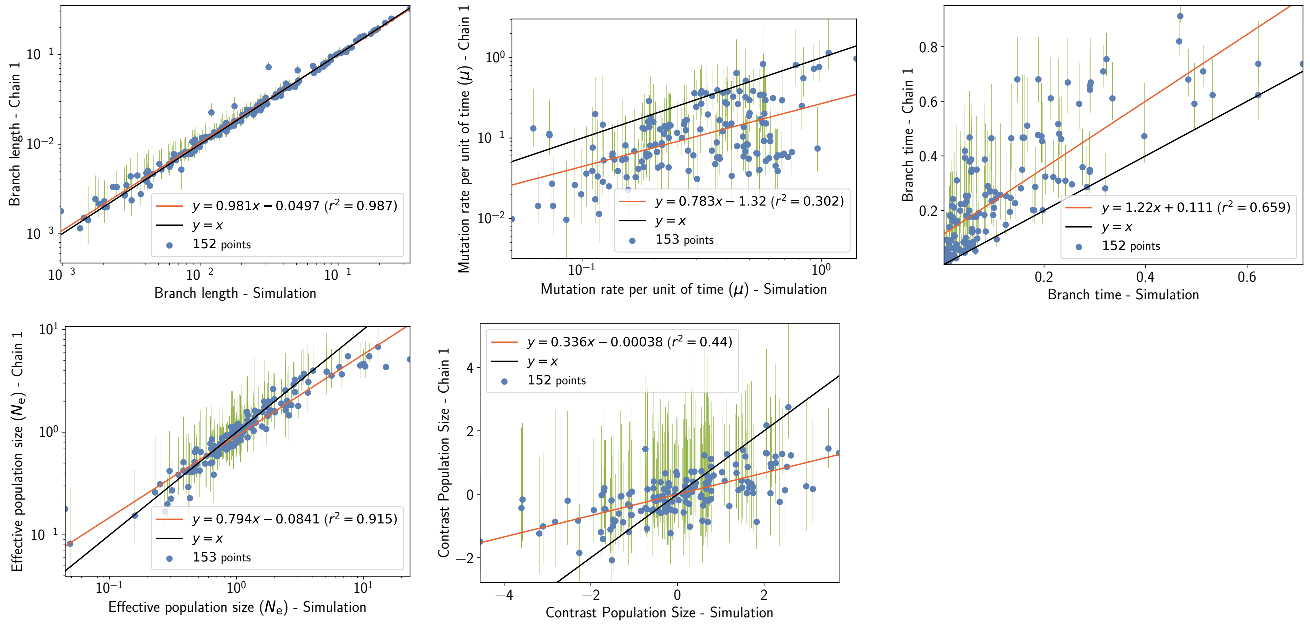


Figure 1: Inferred branch parameters under simulations accounting for site-specific amino-acid profiles, long term fluctuation of $N_\mathrm{e}$, mutation rate per generation and generation time. Estimation is obtained with the mechanistic inference model developed in this paper of site-specific amino-acid fitness profiles and log-Brownian process for $N_\mathrm{e}$, $\mu$ and life-history traits.

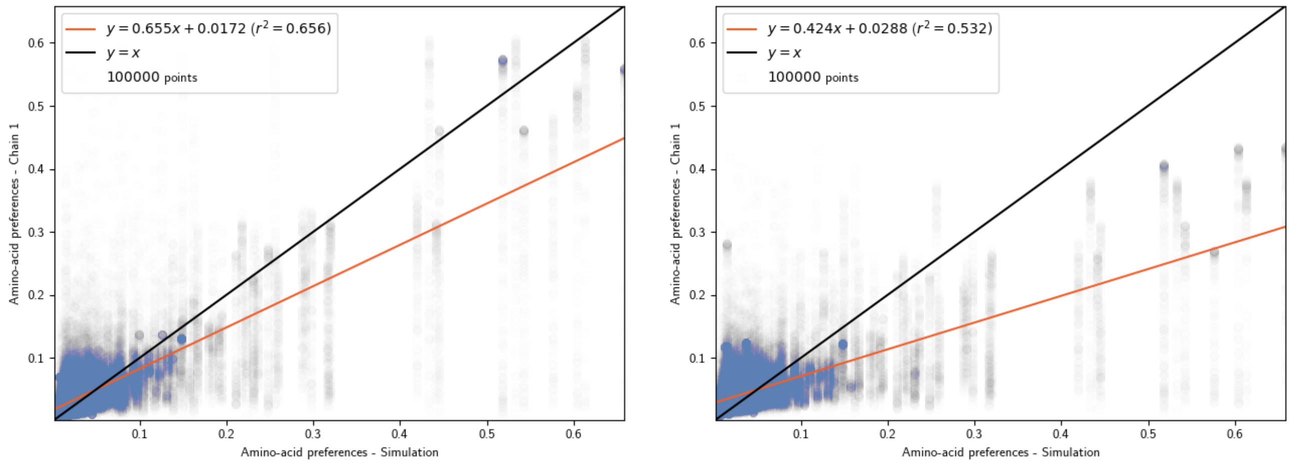


Figure 2: Inferred and simulated site-specific amino-acid profiles under simulation accounting for long term fluctuation of $N_\mathrm{e}$, mutation rate per generation and generation time. Estimation is obtained with the mechanistic inference model developed in this paper of site-specific amino-acid fitness profiles and log-Brownian process for $N_\mathrm{e}$, $\mu$ and life-history traits (in the left panel), or under the assumption of constant $N_\mathrm{e}$ (in the right panel).

| **Experiment** | $\langle\Omega\rangle$ **(branch** $N_\mathrm{e}$**)** | $\langle\Omega\rangle$ **(constant** $N_\mathrm{e}$**)** |
|---|---|---|
| `SimuDiv`, chain 1 | $2.30 \pm 0.04$ | $2.45 \pm 0.02$ |
| `SimuDiv`, chain 2 | $2.30 \pm 0.04$ | $2.45 \pm 0.02$ |

Table 1: Estimated amino-acid entropy under simulations accounting for long term fluctuation of $N_\mathrm{e}$, mutation rate per generation and generation time. Estimation is obtained with the mechanistic inference model developed in this paper of site-specific amino-acid fitness profiles and log-Brownian process for $N_\mathrm{e}$, $\mu$ and life-history traits (in the left column), or under the assumption of constant $N_\mathrm{e}$ (in the right column).

#### 2.2 Wright-Fisher with polymorphism (SimuPoly)

The evolutionary dynamics was formalized as a Wright-Fisher model with mutation, selection and drift The population is assumed to be panmictic, with effective population size $N_{\mathrm{e}}$ and with non-overlapping generations.

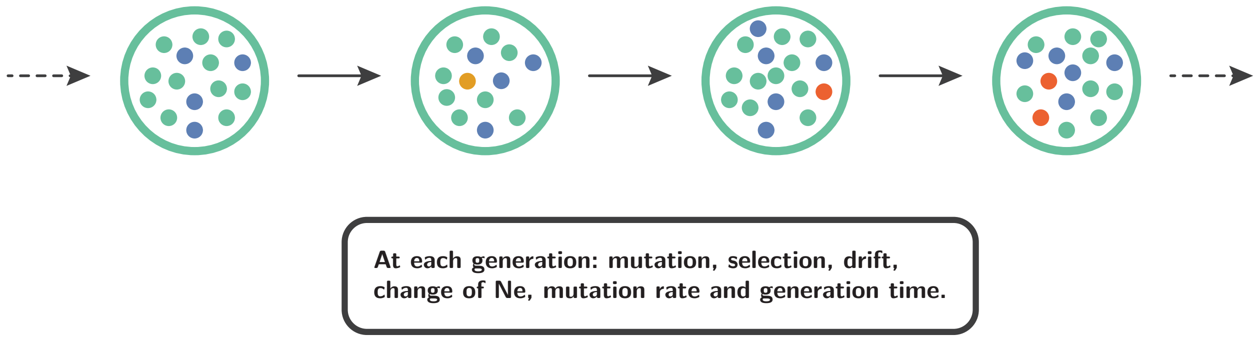


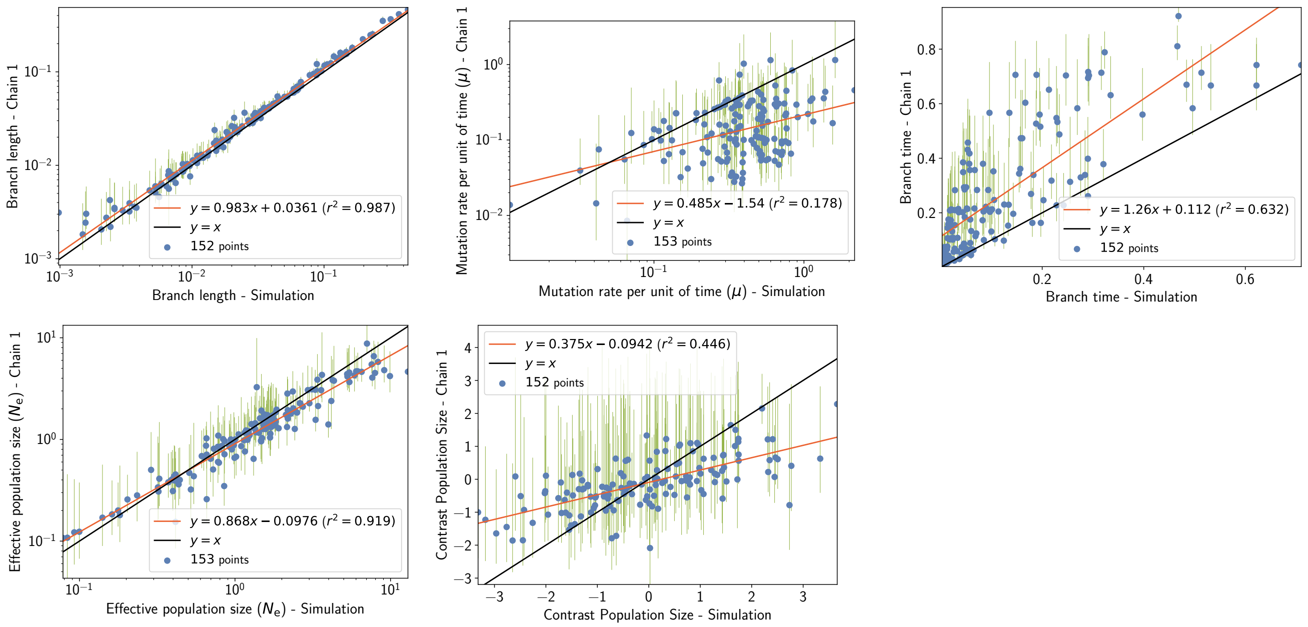


Figure 3: Inferred branch parameters under simulation accounting for finite population effects, site linkage and short term fluctuation of $N_{\mathrm{e}}$. Estimation is obtained with the mechanistic inference model developed in this paper of site-specific amino-acid fitness profiles and log-Brownian process for $N_{\mathrm{e}}$, $\mu$ and life-history traits.

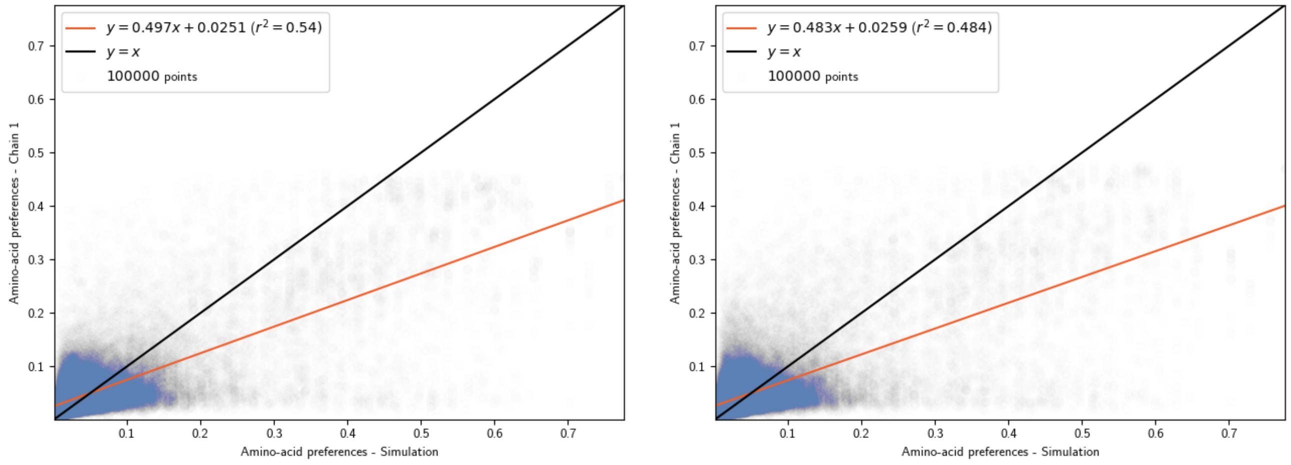


Figure 4: Inferred and simulated site-specific amino-acid profiles under simulation accounting for finite population effects, site linkage and short term fluctuation of $N_{\mathrm{e}}$. Estimation is obtained with the mechanistic inference model developed in this paper of site-specific amino-acid fitness profiles and log-Brownian process for $N_{\mathrm{e}}$, $\mu$ and life-history traits (in the left panel), or under the assumption of constant $N_{\mathrm{e}}$ (in the right panel).

| **Experiment** | $\langle \Omega \rangle$ **(branch** $N_{\mathrm{e}}$**)** | $\langle \Omega \rangle$ **(constant** $N_{\mathrm{e}}$**)** |
|---|---|---|
| `SimuPoly`, chain 1 | $2.47 \pm 0.03$ | $2.37 \pm 0.02$ |
| `SimuPoly`, chain 2 | $2.47 \pm 0.03$ | $2.37 \pm 0.02$ |

Table 2: Estimated amino-acid entropy under simulation accounting for finite population effects, site linkage and short term fluctuation of $N_{\mathrm{e}}$. Estimation is obtained with the mechanistic inference model developed in this paper of site-specific amino-acid fitness profiles and log-Brownian process for $N_{\mathrm{e}}$, $\mu$ and life-history traits (in the left column), or under the assumption of constant $N_{\mathrm{e}}$ (in the right column).

#### 2.3 Fisher geometric landscape (`SimuGeo`)

We simulated substitutions in a protein using an adaptation of Fisher's geometric landscape (Tenaillon, 2014; Blanquart and Bataillon, 2016). In the original context, the phenotype is a vector ($\boldsymbol{P}$) in a multidimensional space, where the number of dimensions is often termed complexity. From a phenotype, the fitness is a monotonously decreasing function of the phenotype distance to 0. The exact functional phenotype-fitness map depends on 2 external parameters controlling for strength ($\alpha$) and epistasis ($\beta$). If the phenotype-fitness map is explicit, the genotype-phenotype map is more pervasive. Mutations are seen has displacement of the phenotype in the multidimensional space. Beneficial mutations are moving the phenotype closer to 0, whereas deleterious mutations are moving the phenotype further away. In such original context, the distribution of mutational effects is not dependent on the current genotype, but this can be relaxed using a genotype-phenotype map.

In a protein context, the genotype-phenotype map can be defined by assigning to each of the 20 amino acid a vector in the multidimensional space. Since different sites of the protein do not have the same physico-chemical properties, we can define a specific genotype-phenotype map for each position of the sequence. Overall, the protein phenotype is computed as the sum of site-specific multidimensional vectors, obtained by accessing the amino acid present at each site of the protein. From a DNA sequence $\mathbb{S}^t$ after $t$ substitutions, the protein's phenotype is given by:

$$\boldsymbol{P}\left(\mathbb{S}^t\right) = \sum_{\mathrm{z}=1}^{\mathrm{Z}} \boldsymbol{P}_{\mathrm{z}}\left(\mathbb{S}^t(\mathrm{z})\right), \tag{7}$$

where $\boldsymbol{P}_{\mathrm{z}}$ is the genotype-phenotype map at site z.

And the Wrightian fitness of $\mathbb{S}^t$ is :

$$W\left(\boldsymbol{P}\left(\mathbb{S}^t\right)\right) = e^{-\alpha \left|\boldsymbol{P}\left(\mathbb{S}^t\right)\right|^{\beta}}, \tag{8}$$

where strength ($\alpha > 0$) and epistasis ($\beta$) are parameters of the fitness function.

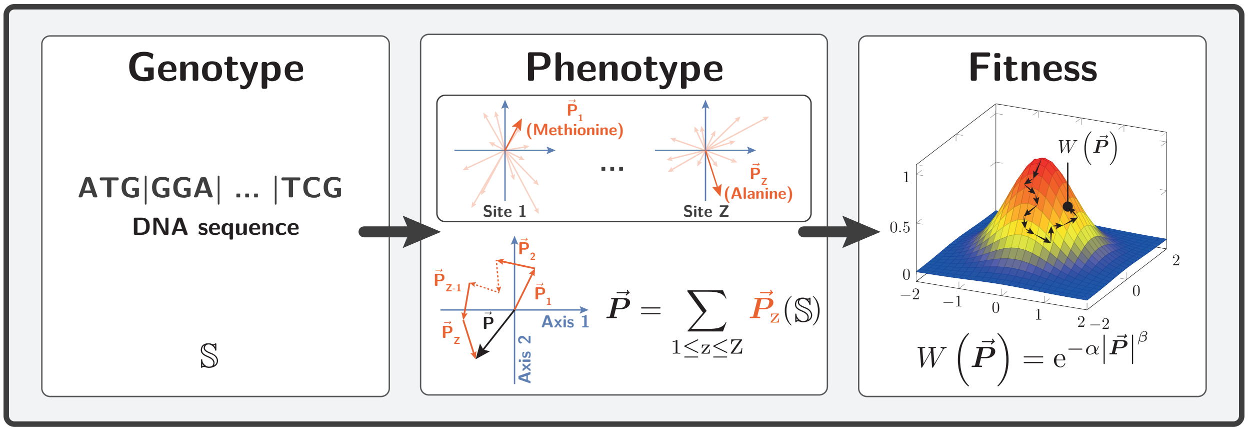


For each possible mutant (at time $t+1$ substitutions), we compute $\boldsymbol{P}\left(\mathbb{S}^{t+1}\right)$ from the updated sequence $\mathbb{S}^{t+1}$, and subsequently the selection coefficient of the mutant:

$$s\left(\mathbb{S}^{t}, \mathbb{S}^{t+1}\right) = \frac{W\left(\boldsymbol{P}\left(\mathbb{S}^{t+1}\right)\right) - W\left(\boldsymbol{P}\left(\mathbb{S}^{t}\right)\right)}{W\left(\boldsymbol{P}\left(\mathbb{S}^{t}\right)\right)}. \tag{9}$$

The next change in the protein coding DNA and the time to next the event is chosen using Gillespie's algorithm (Gillespie, 1977), according to the rates of substitution between codons:

$$Q_{i,j} = \mu_{i,j} \frac{4N_{\mathrm{e}} s\left(\mathbb{S}^{t}, \mathbb{S}^{t+1}\right)}{1 - \mathrm{e}^{-4N_{\mathrm{e}} s\left(\mathbb{S}^{t}, \mathbb{S}^{t+1}\right)}}, \tag{10}$$

where $Q_{i,j} = \mu_{i,j}$ in the case of synonymous substitutions.

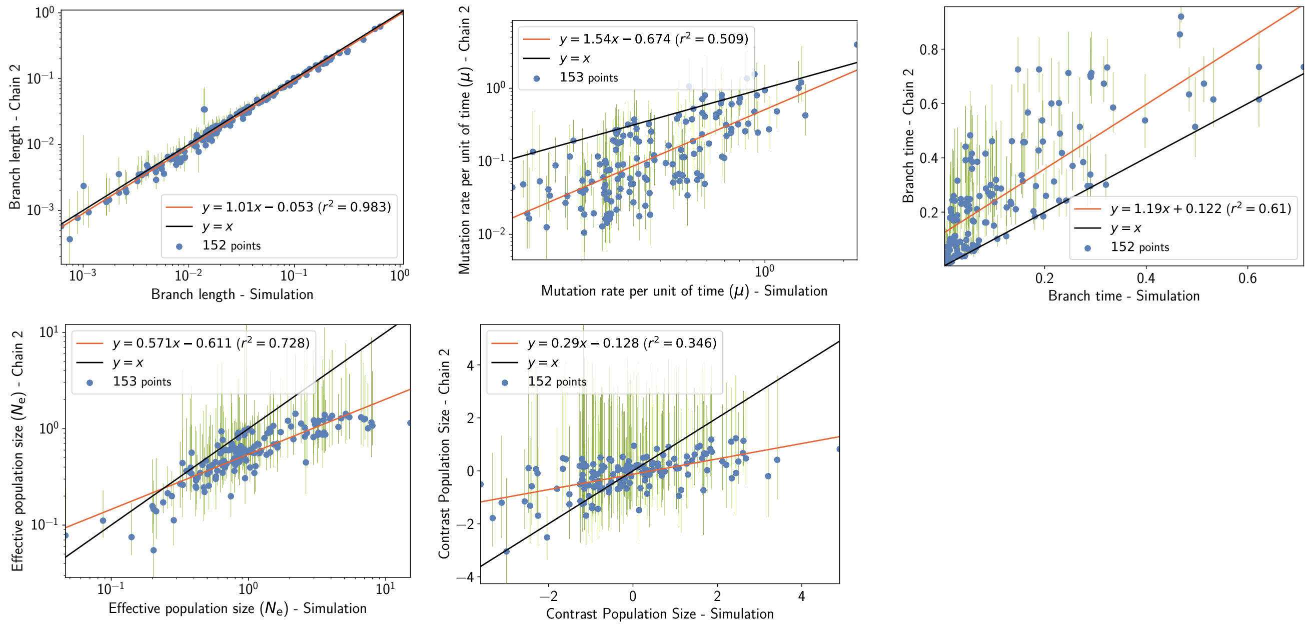


Figure 5: Inferred branch parameters under simulation accounting for site epistasis in geometric landscape, thus fluctuation of the selection coefficient along the phylogeny. Estimation is obtained with the mechanistic inference model developed in this paper of site-specific amino-acid fitness profiles and log-Brownian process for $N_{\mathrm{e}}$, $\mu$ and life-history traits.

| Experiment | $\langle\Omega\rangle$ (branch $N_e$) | $\langle\Omega\rangle$ (constant $N_e$) |
|---|---|---|
| SimuGeo, chain 1 | $2.27 \pm 0.02$ | $2.46 \pm 0.02$ |
| SimuGeo, chain 2 | $2.23 \pm 0.04$ | $2.46 \pm 0.02$ |

Table 3: Estimated amino-acid entropy under simulation accounting for site epistasis (geometric landscape), thus fluctuation of the selection coefficient along the phylogeny. Estimation is obtained with the mechanistic inference model developed in this paper of site-specific amino-acid fitness profiles and log-Brownian process for $N_e$, $\mu$ and life-history traits (in the left column), or under the assumption of constant $N_e$ (in the right column).

#### 2.4 Protein folding probability (SimuFold)

We simulated substitutions in the protein phosphatase (Z = 300 codon sites) as in Goldstein and Pollock (2017). From a DNA sequence $\mathbb{S}^t$ after $t$ substitutions, we compute the free energy of the folded state $G_F(\mathbb{S}^t)$, using the 3-dimensional structure of the folded state and pair-wise contact energies between neighboring amino-acid residues:

$$G_F\left(\mathbb{S}^t\right) = \sum_{z=1}^{Z} \sum_{r \in \mathcal{V}(z)} I\left(\mathbb{S}^t(z), \mathbb{S}^t(r)\right), \tag{11}$$

where $I(a, b)$ is the pair-wise contact energies between amino acid $a$ and $b$, using contact potentials estimated by Miyazawa and Jernigan (1985), and $\mathcal{V}(z)$ are the neighbor residues of site z (closer than 7Å) in the 3D structure.

The free energy of unfolded states $G_U(\mathbb{S}^t)$ is approximated using 55 decoy 3D structures that supposedly represent a sample of possible unfolded states:

$$G_U\left(\mathbb{S}^t\right) = \left\langle G\left(\mathbb{S}^t\right)\right\rangle - kT\ln(1.0\mathrm{E}^{160}) - \frac{2\left[\left\langle G\left(\mathbb{S}^t\right)^2\right\rangle - \left\langle G\left(\mathbb{S}^t\right)\right\rangle^2\right]}{kT} \tag{12}$$

where the average $\langle . \rangle$ runs other the 55 decoy 3D structures, and $k$ is the Boltzmann constant and $T$ the temperature in Kelvin.

From the energy of folded and unfolded states, we can compute the difference in free energy between the states:

$$\Delta G\left(\mathbb{S}^t\right) = G_F\left(\mathbb{S}^t\right) - G_U\left(\mathbb{S}^t\right) \tag{13}$$

Wrightian fitness is defined as the probability of our protein to be in the folded state:

$$W\left(\Delta G\left(\mathbb{S}^t\right)\right) = \mathbb{P}_F\left(\mathbb{S}^t\right) = \frac{e^{-\beta G_F\left(\mathbb{S}^t\right)}}{e^{-\beta G_F\left(\mathbb{S}^t\right)} + e^{-\beta G_U\left(\mathbb{S}^t\right)}} = \frac{1}{1 + e^{\beta \Delta G\left(\mathbb{S}^t\right)}}, \tag{14}$$

where $\beta$ is the inverse of the temperature ($\beta = 1/kT$).

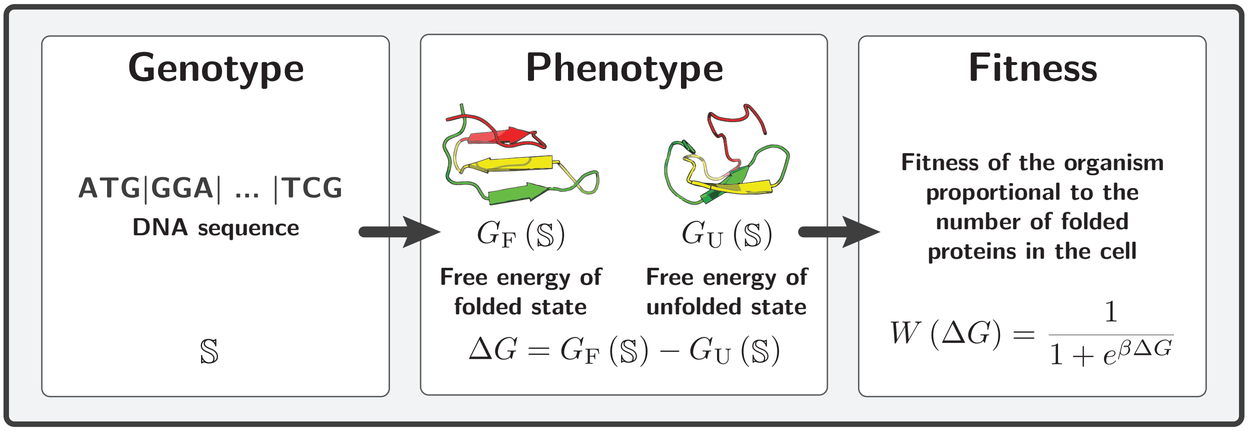

For each possible mutant (at time $t+1$ substitutions), we compute $\Delta G^{t+1}$ from the updated sequence $\mathbb{S}^{t+1}$, and subsequently the selection coefficient of the mutant:

$$s\left(\mathbb{S}^t, \mathbb{S}^{t+1}\right) = \frac{W\left(\Delta G\left(\mathbb{S}^{t+1}\right)\right) - W\left(\Delta G\left(\mathbb{S}^{t}\right)\right)}{W\left(\Delta G\left(\mathbb{S}^{t}\right)\right)}. \tag{15}$$

The next change in the protein coding DNA and the time to next the event is chosen using Gillespie's algorithm (Gillespie, 1977), according to the rates of substitution between codons:

$$Q_{i,j} = \mu_{i,j} \frac{4N_{\mathrm{e}} s\left(\mathbb{S}^t, \mathbb{S}^{t+1}\right)}{1 - \mathrm{e}^{-4N_{\mathrm{e}} s\left(\mathbb{S}^t, \mathbb{S}^{t+1}\right)}}, \tag{16}$$

where $Q_{i,j} = \mu_{i,j}$ in the case of synonymous substitutions.

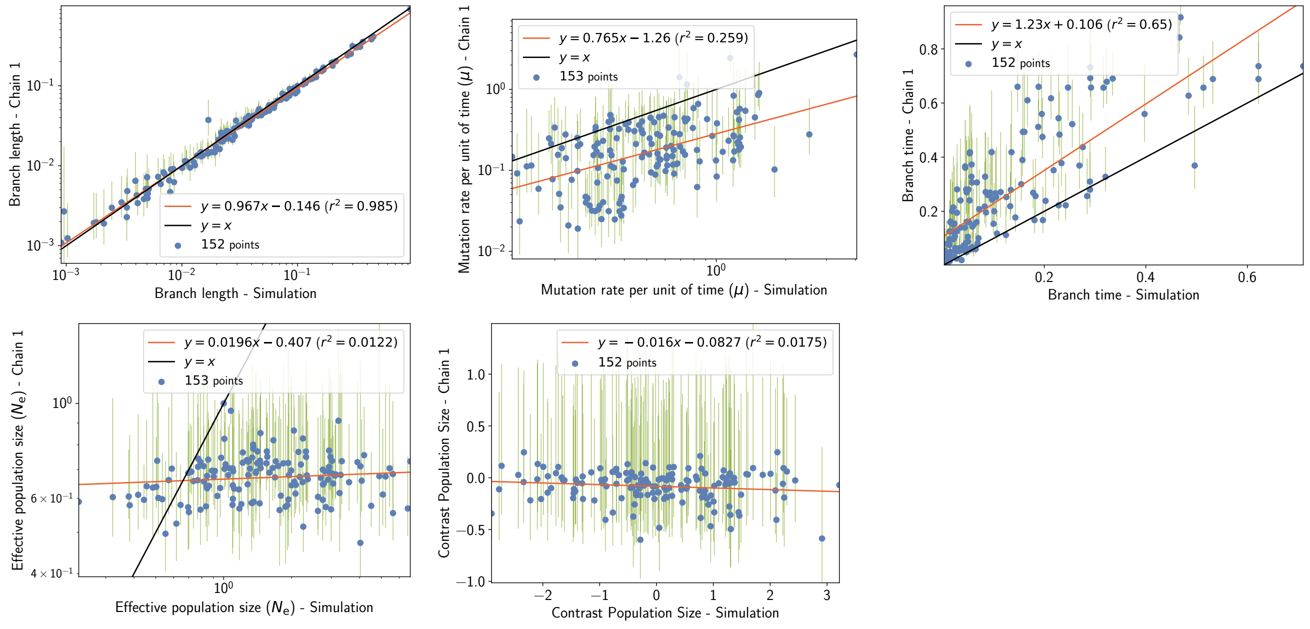


Figure 6: Inferred branch parameters under simulation accounting for site epistasis (folding stability model), thus fluctuation of the selection coefficient along the phylogeny. Estimation is obtained with the mechanistic inference model developed in this paper of site-specific amino-acid fitness profiles and log-Brownian process for $N_{\mathrm{e}}$, $\mu$ and life-history traits.

| **Experiment** | $\langle\Omega\rangle$ **(branch** $N_{\mathrm{e}}$**)** | $\langle\Omega\rangle$ **(constant** $N_{\mathrm{e}}$**)** |
|---|---|---|
| `SimuFold`, chain 1 | $1.31 \pm 0.05$ | $1.61 \pm 0.03$ |
| `SimuFold`, chain 2 | $1.30 \pm 0.04$ | $1.60 \pm 0.03$ |

Table 4: Estimated amino-acid entropy under simulation accounting for site epistasis (folding stability model), thus fluctuation of the selection coefficient along the phylogeny. Obtained with the mechanistic inference model developed in this paper of site-specific amino-acid fitness profiles and log-Brownian process for $N_{\mathrm{e}}$, $\mu$ and life-history traits (in the left column), or under the assumption of constant $N_{\mathrm{e}}$ (in the right column).

### 3 Empirical data in mammals

#### 3.1 Chain convergence

Obtained with the mechanistic inference model developed in this paper of site-specific amino-acid fitness profiles and log-Brownian process for $N_{\mathrm{e}}$, $\mu$ and life-history traits.

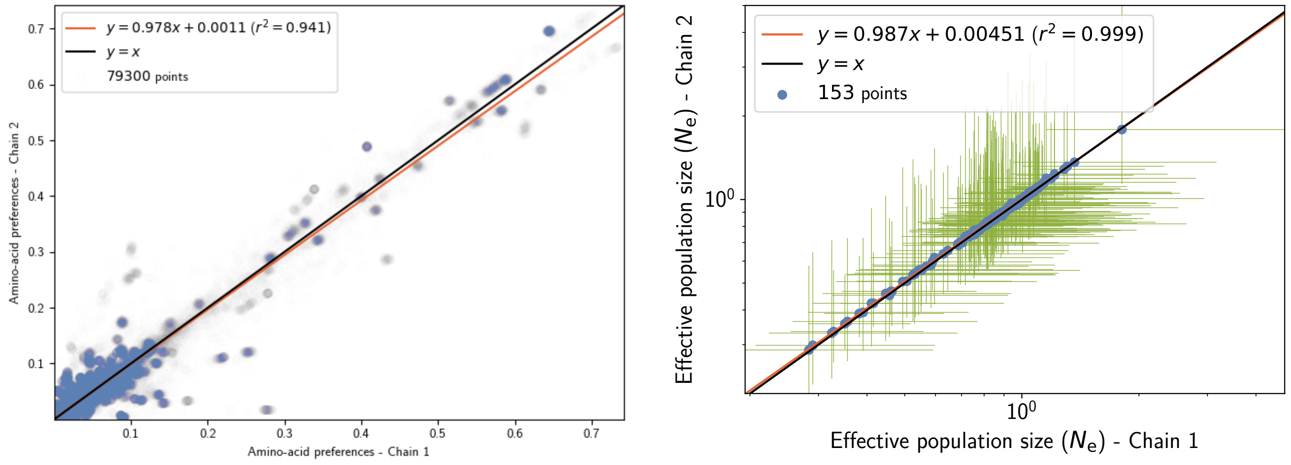


Figure 7: Chain convergence of site amino-acid preferences (left panel) and branch $N_e$ (right panel).

#### 3.2 Traits estimation & correlation (replicate 1, chain 1)

Obtained with the mechanistic inference model developed in this paper of site-specific amino-acid fitness profiles and log-Brownian process for $N_e$, $\mu$ and life-history traits.

| **Covariance ($\Sigma$)** | $N_e$ | $\mu$ | **Maximum longevity** | **Adult weight** | **Female maturity** |
|---|---|---|---|---|---|
| $N_e$ | 0.281** | 0.324** | −0.268** | −1.29** | −0.308** |
| $\mu$ | - | 1.93** | −1.12** | −5.19** | −1.43** |
| **Maximum longevity** | - | - | 0.934** | 3.58** | 1.01** |
| **Adult weight** | - | - | - | 19.9** | 4.48** |
| **Female maturity** | - | - | - | - | 1.53** |

Table 5: Covariance coefficient between effective population size ($N_e$), mutation rate per site per unit of time ($\mu$), and life-history traits (maximum longevity, adult weight and female maturity) were computed in placental mammals. Asterisks indicate strength of support (*$pp > 0.95$, **$pp > 0.975$).

| **Partial coefficient** | $N_e$ | $\mu$ | **Maximum longevity** | **Adult weight** | **Female maturity** |
|---|---|---|---|---|---|
| $N_e$ | - | −0.146 | −0.177 | −0.265* | −0.0223 |
| $\mu$ | - | - | −0.283* | −0.396** | −0.327** |
| **Maximum longevity** | - | - | - | 0.236* | 0.383** |
| **Adult weight** | - | - | - | - | 0.179 |
| **Female maturity** | - | - | - | - | - |

Table 6: Partial correlation coefficient between effective population size ($N_e$), mutation rate per site per unit of time ($\mu$), and life-history traits (maximum longevity, adult weight and female maturity) were computed in placental mammals. Asterisks indicate strength of support (*$pp > 0.95$, **$pp > 0.975$).

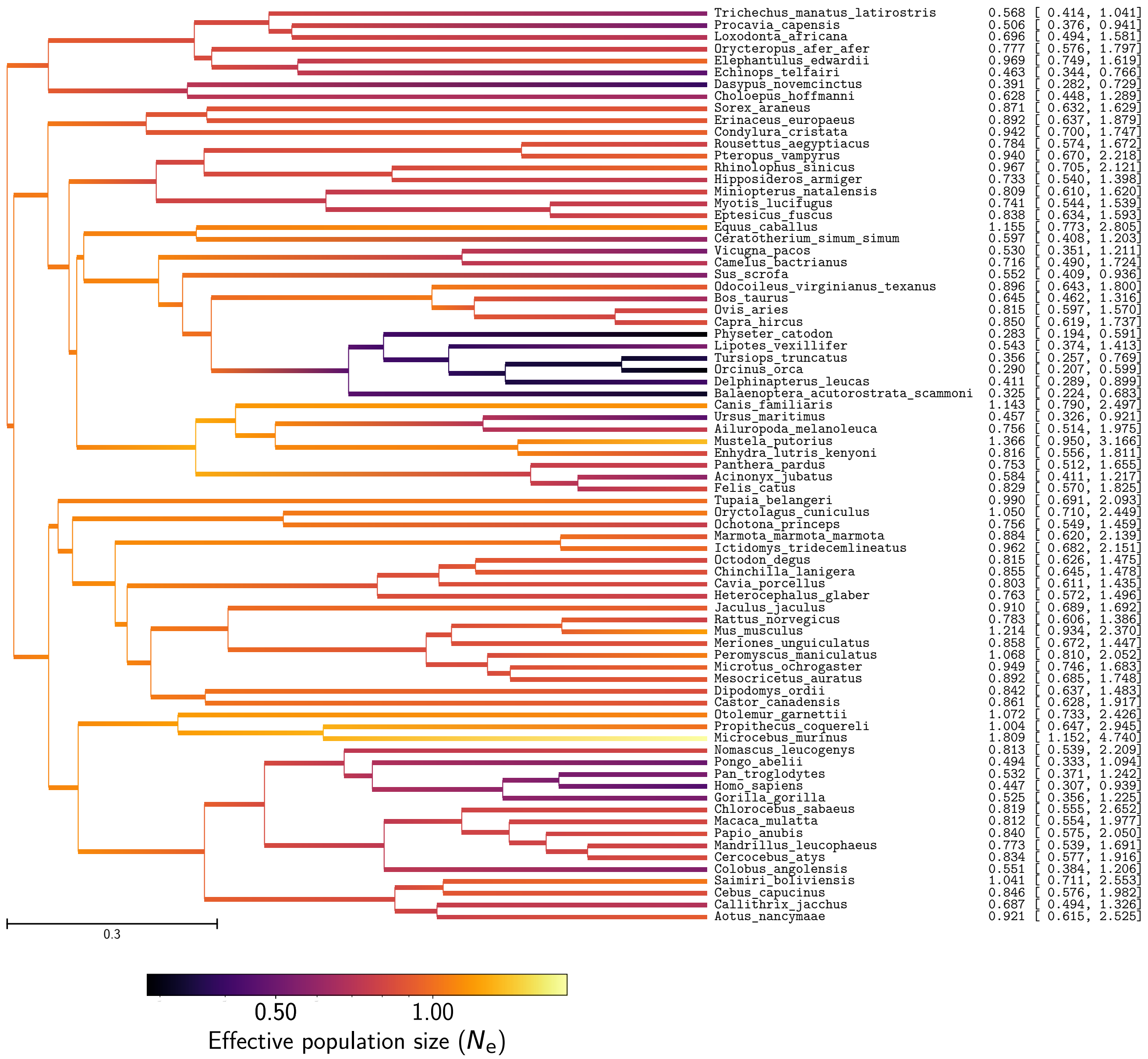


Figure 8: Effective population size ($N_{\mathrm{e}}$) estimation in mammals

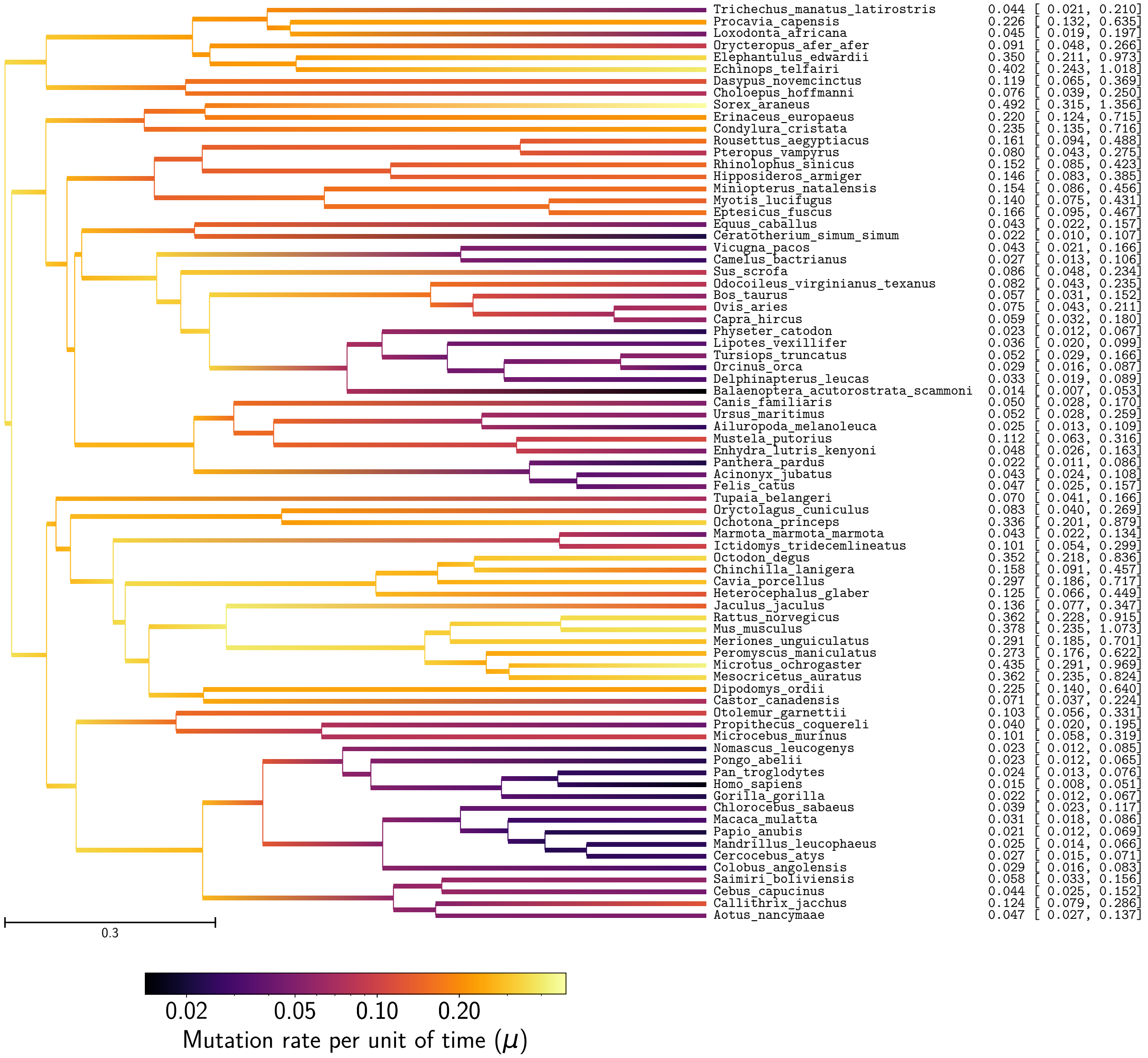


Figure 9: Mutation rate ($\mu$) estimation in mammals

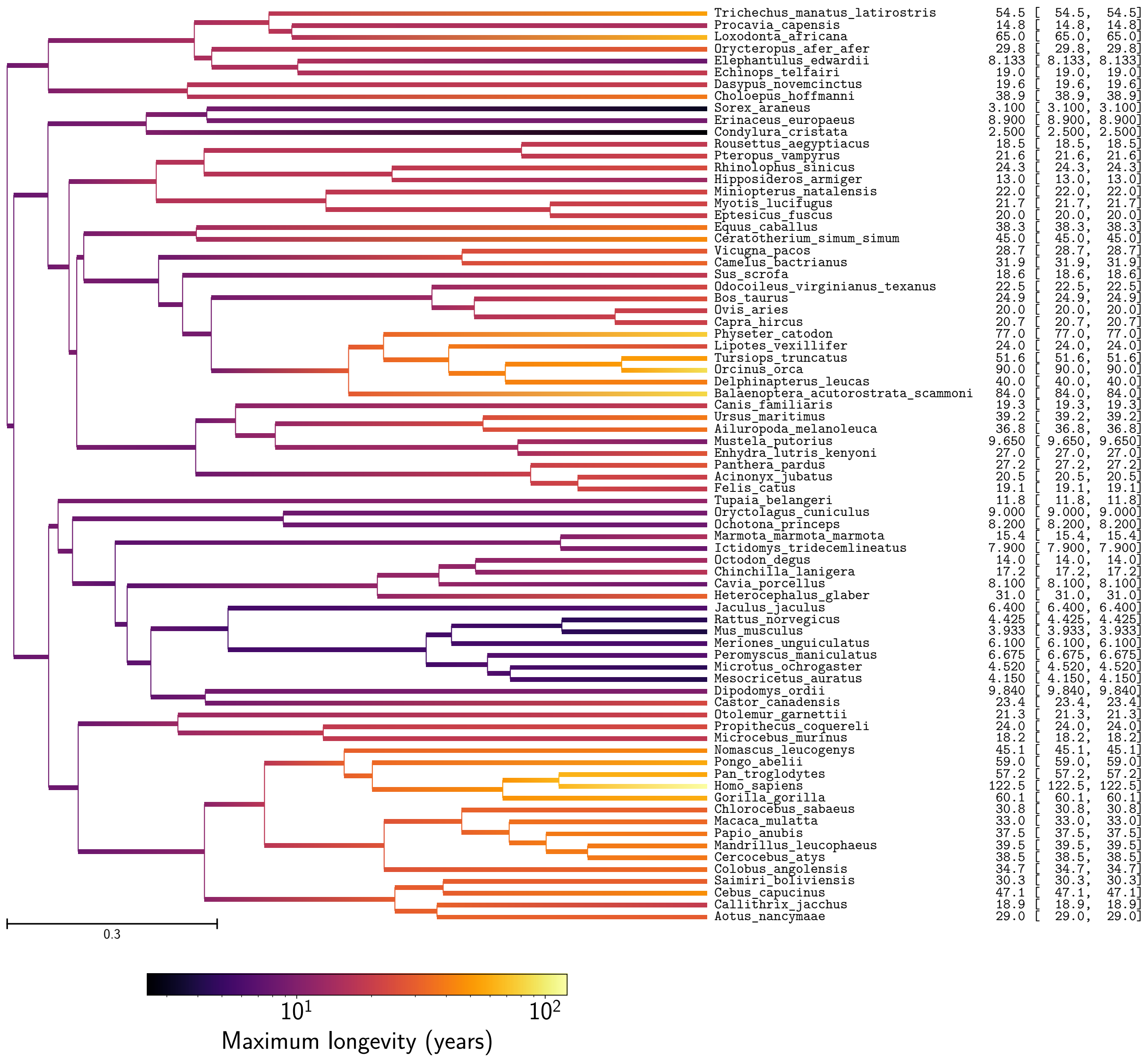


Figure 10: Maximum longevity estimation in mammals

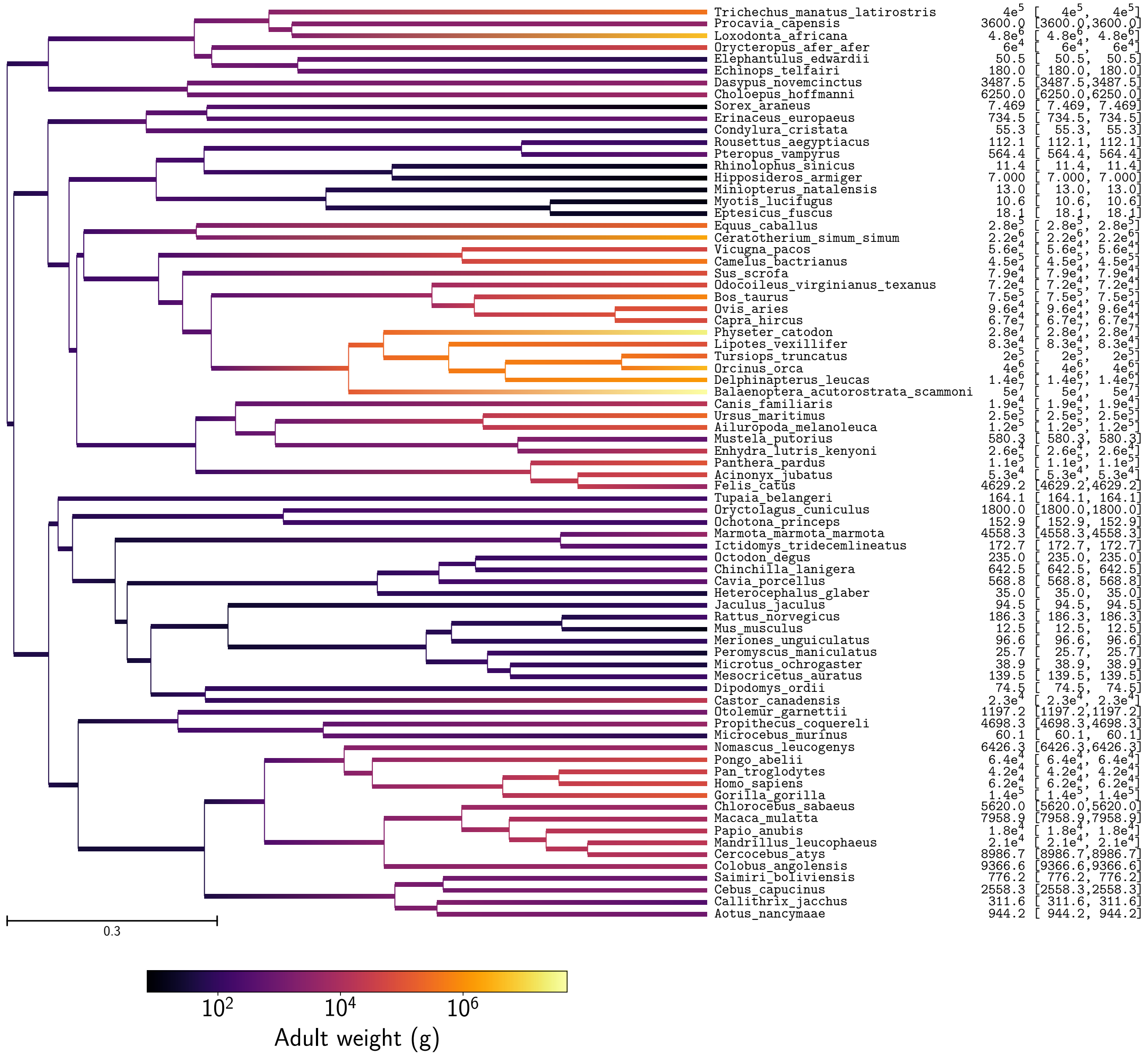


Figure 11: Adult weight estimation in mammals

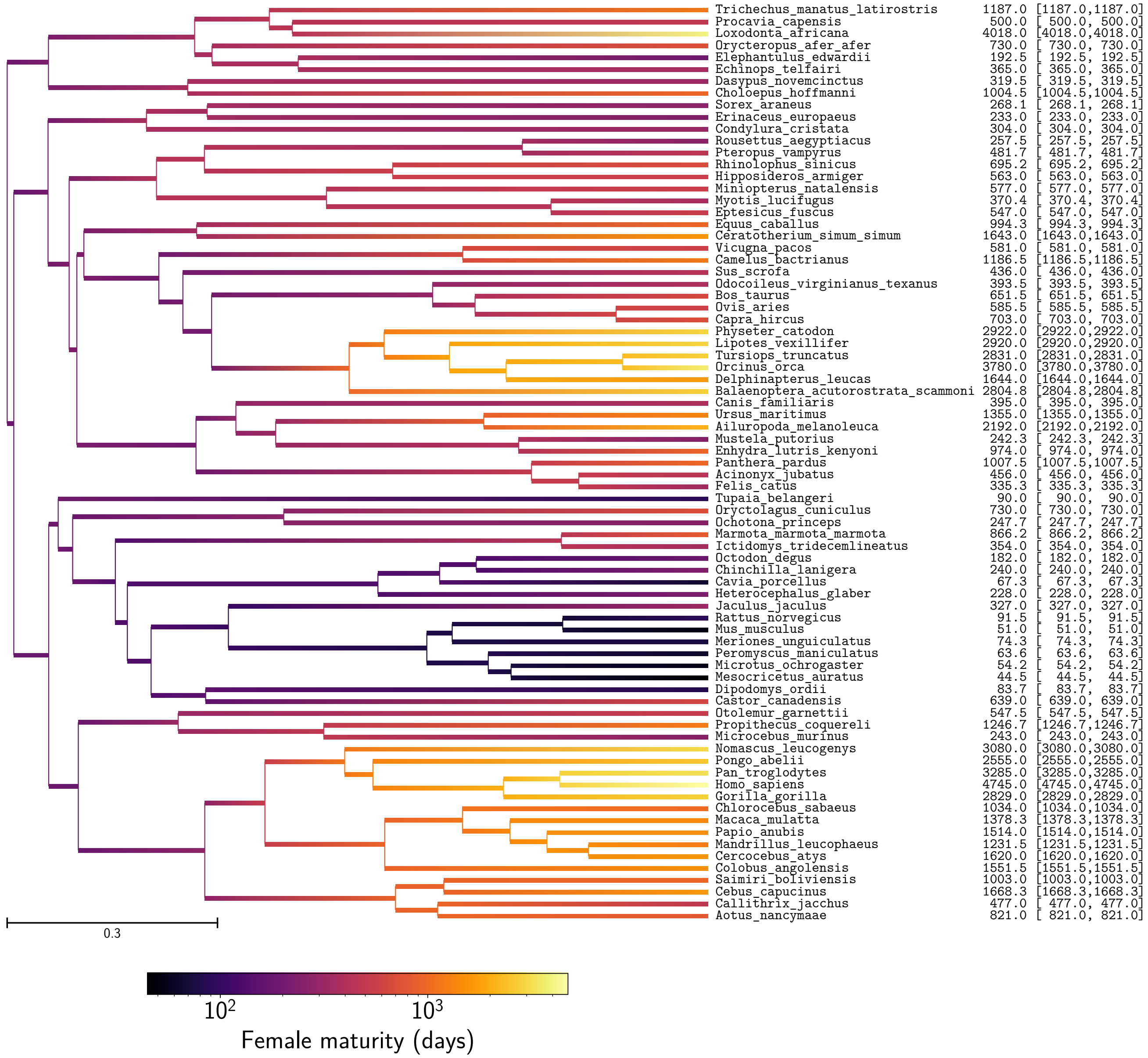


Figure 12: Female maturity estimation in mammals

#### 3.3 Repeatability of experiments

4 independent inferences were performed on a randomly chosen set of 18 coding sequences (CDS) out of 226. Obtained with the mechanistic inference model developed in this paper of site-specific amino-acid fitness profiles and log-Brownian process for $N_e$, $\mu$ and life-history traits. Each plot is a correlation between a pair of experiments for a given parameter. For each node (or branch) of the tree, the mean posterior of the parameter over the MCMC (after burn-in) is represented in blue dots, green solid lines are the 90% confidence interval of the MCMC. Solid red line is the regression line between replicates.

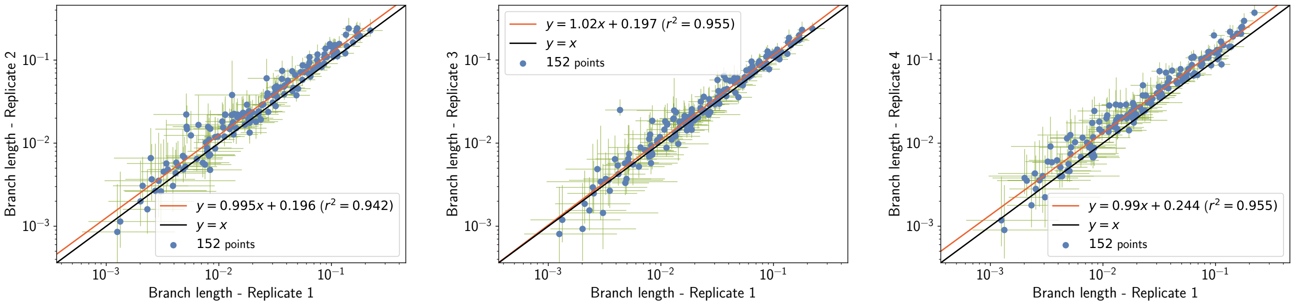


Figure 13: Repeatability of branch length ($l$) estimation in mammals

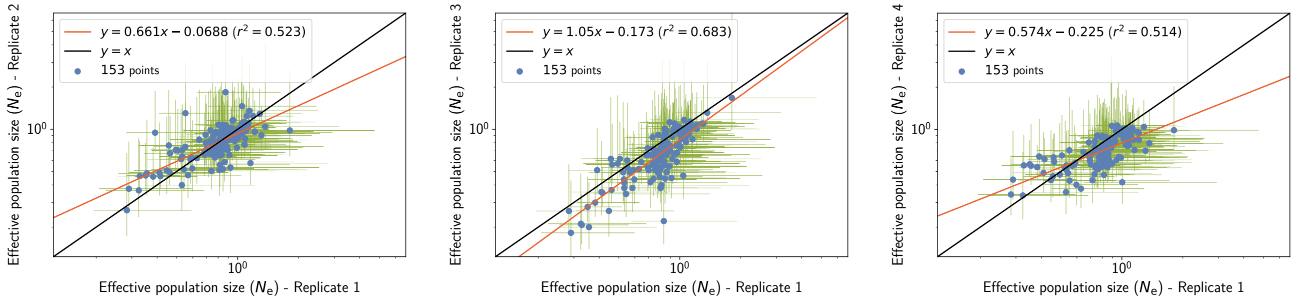


Figure 14: Repeatability of effective population size ($N_{\mathrm{e}}$) estimation in mammals

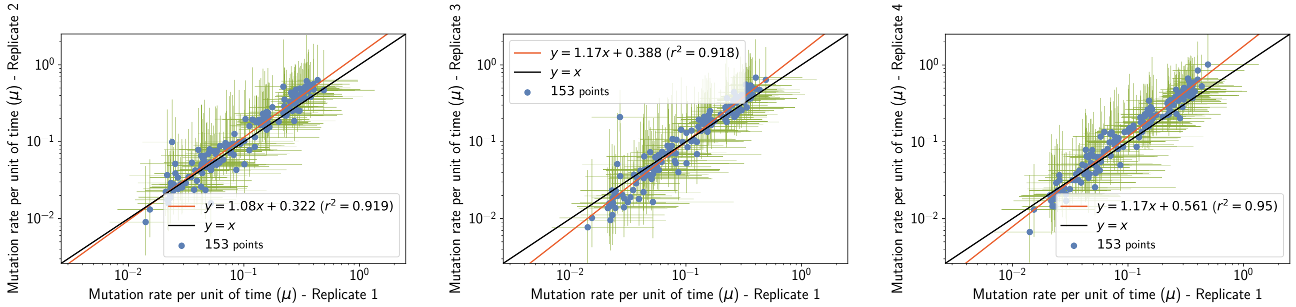


Figure 15: Repeatability of mutation rate ($\mu$) estimation in mammals

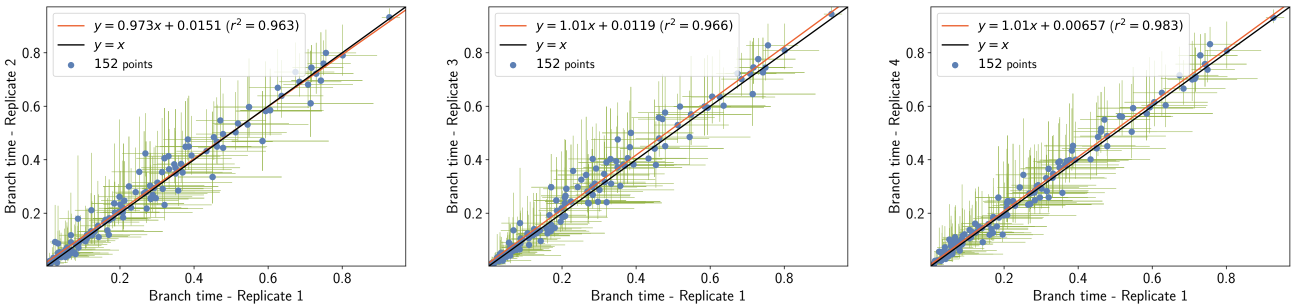


Figure 16: Repeatability of branch time ($\Delta T$) estimation in mammals

| Rep. 1 | Rep. 2 | Rep. 3 | Rep. 4 | Taxon |
|---|---|---|---|---|
| 0.568 | 0.469 | 0.489 | 0.548 | *Trichechus manatus latirostris* |
| 0.506 | 0.706 | 0.615 | 0.65 | *Procavia capensis* |
| 0.696 | 0.799 | 0.532 | 0.595 | *Loxodonta africana* |
| 0.777 | 0.812 | 0.651 | 0.717 | *Orycteropus afer afer* |
| 0.969 | 0.904 | 0.68 | 0.949 | *Elephantulus edwardii* |
| 0.463 | 0.673 | 0.56 | 0.586 | *Echinops telfairi* |
| 0.391 | 0.945 | 0.51 | 0.639 | *Dasypus novemcinctus* |
| 0.628 | 0.621 | 0.52 | 0.376 | *Choloepus hoffmanni* |
| 0.871 | 1.84 | 0.745 | 0.819 | *Sorex araneus* |
| 0.892 | 0.833 | 1.12 | 1.06 | *Erinaceus europaeus* |
| 0.942 | 1.15 | 1.1 | 0.916 | *Condylura cristata* |
| 0.784 | 0.679 | 0.488 | 0.535 | *Rousettus aegyptiacus* |
| 0.94 | 0.838 | 0.662 | 0.604 | *Pteropus vampyrus* |
| 0.967 | 0.823 | 0.586 | 0.636 | *Rhinolophus sinicus* |
| 0.733 | 0.98 | 0.876 | 0.746 | *Hipposideros armiger* |
| 0.809 | 0.934 | 0.742 | 0.738 | *Miniopterus natalensis* |
| 0.741 | 0.504 | 0.442 | 0.53 | *Myotis lucifugus* |
| 0.838 | 0.849 | 0.588 | 0.753 | *Eptesicus fuscus* |
| 1.16 | 0.573 | 0.846 | 0.711 | *Equus caballus* |
| 0.597 | 0.524 | 0.438 | 0.402 | *Ceratotherium simum simum* |
| 0.53 | 0.399 | 0.438 | 0.356 | *Vicugna pacos* |
| 0.716 | 0.68 | 0.418 | 0.432 | *Camelus bactrianus* |
| 0.552 | 1.3 | 0.531 | 0.43 | *Sus scrofa* |
| 0.896 | 0.861 | 0.761 | 0.568 | *Odocoileus virginianus texanus* |
| 0.645 | 0.844 | 0.583 | 0.69 | *Bos taurus* |
| 0.815 | 0.649 | 0.747 | 0.473 | *Ovis aries* |
| 0.85 | 0.723 | 0.742 | 0.538 | *Capra hircus* |
| 0.283 | 0.264 | 0.261 | 0.342 | *Physeter catodon* |
| 0.543 | 0.517 | 0.345 | 0.486 | *Lipotes vexillifer* |
| 0.356 | 0.484 | 0.2 | 0.549 | *Tursiops truncatus* |
| 0.29 | 0.376 | 0.182 | 0.437 | *Orcinus orca* |
| 0.411 | 0.491 | 0.356 | 0.488 | *Delphinapterus leucas* |
| 0.325 | 0.366 | 0.211 | 0.337 | *Balaenoptera acutorostrata scammoni* |
| 1.14 | 1.35 | 1 | 0.842 | *Canis familiaris* |
| 0.457 | 0.766 | 0.615 | 0.461 | *Ursus maritimus* |
| 0.756 | 0.716 | 0.569 | 0.53 | *Ailuropoda melanoleuca* |
| 1.37 | 1.04 | 1.31 | 0.795 | *Mustela putorius* |
| 0.816 | 0.557 | 0.7 | 0.657 | *Enhydra lutris kenyoni* |
| 0.753 | 0.881 | 0.683 | 0.556 | *Panthera pardus* |
| 0.584 | 0.61 | 0.381 | 0.561 | *Acinonyx jubatus* |
| 0.829 | 0.761 | 0.655 | 0.602 | *Felis catus* |
| 0.99 | 0.738 | 1.04 | 0.627 | *Tupaia belangeri* |
| 1.05 | 1.46 | 1.17 | 0.897 | *Oryctolagus cuniculus* |
| 0.756 | 0.751 | 0.514 | 0.902 | *Ochotona princeps* |
| 0.884 | 0.746 | 0.541 | 0.862 | *Marmota marmota marmota* |
| 0.962 | 0.933 | 0.773 | 0.977 | *Ictidomys tridecemlineatus* |
| 0.815 | 0.733 | 0.688 | 0.874 | *Octodon degus* |
| 0.855 | 0.979 | 0.691 | 0.645 | *Chinchilla lanigera* |
| 0.803 | 1.08 | 0.684 | 0.898 | *Cavia porcellus* |
| 0.763 | 0.76 | 0.702 | 0.518 | *Heterocephalus glaber* |
| 0.91 | 0.655 | 0.449 | 0.865 | *Jaculus jaculus* |
| 0.783 | 0.956 | 0.91 | 0.883 | *Rattus norvegicus* |
| 1.21 | 0.963 | 1.01 | 0.839 | *Mus musculus* |
| 0.858 | 0.856 | 0.818 | 0.828 | *Meriones unguiculatus* |
| 1.07 | 1.11 | 0.877 | 0.757 | *Peromyscus maniculatus* |
| 0.949 | 1.16 | 1.01 | 1.06 | *Microtus ochrogaster* |
| 0.892 | 0.949 | 1.12 | 0.788 | *Mesocricetus auratus* |
| 0.842 | 1.07 | 1.01 | 0.695 | *Dipodomys ordii* |
| 0.861 | 0.583 | 0.494 | 0.575 | *Castor canadensis* |
| 1.07 | 1.13 | 0.812 | 0.821 | *Otolemur garnettii* |
| 1 | 0.945 | 0.741 | 0.418 | *Propithecus coquereli* |
| 1.81 | 0.98 | 1.67 | 0.985 | *Microcebus murinus* |
| 0.813 | 0.512 | 0.399 | 0.582 | *Nomascus leucogenys* |
| 0.494 | 0.71 | 0.568 | 0.475 | *Pongo abelii* |
| 0.532 | 0.713 | 0.381 | 0.513 | *Pan troglodytes* |
| 0.447 | 0.508 | 0.261 | 0.433 | *Homo sapiens* |
| 0.525 | 0.611 | 0.493 | 0.528 | *Gorilla gorilla* |
| 0.819 | 0.754 | 0.782 | 0.71 | *Chlorocebus sabaeus* |
| 0.812 | 0.816 | 0.538 | 0.679 | *Macaca mulatta* |
| 0.84 | 0.8 | 0.555 | 0.676 | *Papio anubis* |
| 0.773 | 0.813 | 0.501 | 0.628 | *Mandrillus leucophaeus* |
| 0.834 | 0.823 | 0.221 | 0.631 | *Cercocebus atys* |
| 0.551 | 0.749 | 0.599 | 0.706 | *Colobus angolensis* |
| 1.04 | 0.93 | 0.466 | 0.859 | *Saimiri boliviensis* |
| 0.846 | 0.519 | 0.444 | 0.667 | *Cebus capucinus* |
| 0.687 | 0.658 | 0.659 | 0.805 | *Callithrix jacchus* |
| 0.921 | 0.532 | 0.614 | 0.794 | *Aotus nancymaae* |
| **6.38** | **6.96** | **9.19** | **3.16** | **Maximum range** |

Table 7: Repeatability of effective population size ($N_e$) estimation in mammals, for the extant taxa.

| Rep. 1 | Rep. 2 | Rep. 3 | Rep. 4 | Taxon |
|---|---|---|---|---|
| 0.0436 | 0.039 | 0.0645 | 0.0422 | *Trichechus manatus latirostris* |
| 0.226 | 0.281 | 0.369 | 0.229 | *Procavia capensis* |
| 0.0455 | 0.0656 | 0.0435 | 0.0342 | *Loxodonta africana* |
| 0.0909 | 0.0995 | 0.144 | 0.0862 | *Orycteropus afer afer* |
| 0.35 | 0.614 | 0.434 | 0.457 | *Elephantulus edwardii* |
| 0.402 | 0.478 | 0.684 | 0.848 | *Echinops telfairi* |
| 0.119 | 0.194 | 0.104 | 0.0861 | *Dasypus novemcinctus* |
| 0.0764 | 0.0713 | 0.109 | 0.0915 | *Choloepus hoffmanni* |
| 0.492 | 0.468 | 0.64 | 1.01 | *Sorex araneus* |
| 0.22 | 0.521 | 0.281 | 0.317 | *Erinaceus europaeus* |
| 0.235 | 0.228 | 0.282 | 0.352 | *Condylura cristata* |
| 0.161 | 0.173 | 0.186 | 0.266 | *Rousettus aegyptiacus* |
| 0.08 | 0.0942 | 0.0565 | 0.103 | *Pteropus vampyrus* |
| 0.152 | 0.222 | 0.157 | 0.158 | *Rhinolophus sinicus* |
| 0.146 | 0.232 | 0.208 | 0.164 | *Hipposideros armiger* |
| 0.154 | 0.166 | 0.135 | 0.23 | *Miniopterus natalensis* |
| 0.14 | 0.164 | 0.174 | 0.158 | *Myotis lucifugus* |
| 0.166 | 0.154 | 0.166 | 0.217 | *Eptesicus fuscus* |
| 0.0428 | 0.0657 | 0.0345 | 0.0404 | *Equus caballus* |
| 0.0217 | 0.0368 | 0.0158 | 0.017 | *Ceratotherium simum simum* |
| 0.0431 | 0.0512 | 0.0335 | 0.0788 | *Vicugna pacos* |
| 0.027 | 0.0244 | 0.0357 | 0.0244 | *Camelus bactrianus* |
| 0.0859 | 0.0431 | 0.0402 | 0.0497 | *Sus scrofa* |
| 0.0818 | 0.0677 | 0.0525 | 0.0805 | *Odocoileus virginianus texanus* |
| 0.057 | 0.0696 | 0.0575 | 0.0424 | *Bos taurus* |
| 0.0753 | 0.0883 | 0.0886 | 0.101 | *Ovis aries* |
| 0.0586 | 0.0764 | 0.0653 | 0.0641 | *Capra hircus* |
| 0.0234 | 0.0291 | 0.0161 | 0.0181 | *Physeter catodon* |
| 0.0355 | 0.0534 | 0.0373 | 0.038 | *Lipotes vexillifer* |
| 0.0524 | 0.0474 | 0.0481 | 0.0393 | *Tursiops truncatus* |
| 0.0292 | 0.0254 | 0.0184 | 0.0171 | *Orcinus orca* |
| 0.0331 | 0.0347 | 0.019 | 0.0318 | *Delphinapterus leucas* |
| 0.0141 | 0.00903 | 0.00772 | 0.0067 | *Balaenoptera acutorostrata scammoni* |
| 0.0505 | 0.0795 | 0.0329 | 0.065 | *Canis familiaris* |
| 0.0518 | 0.0158 | 0.0165 | 0.0256 | *Ursus maritimus* |
| 0.0255 | 0.0517 | 0.0513 | 0.0368 | *Ailuropoda melanoleuca* |
| 0.112 | 0.0774 | 0.101 | 0.163 | *Mustela putorius* |
| 0.0479 | 0.0513 | 0.0456 | 0.0528 | *Enhydra lutris kenyoni* |
| 0.0218 | 0.0175 | 0.0144 | 0.0194 | *Panthera pardus* |
| 0.0426 | 0.0261 | 0.039 | 0.0486 | *Acinonyx jubatus* |
| 0.0465 | 0.0235 | 0.038 | 0.038 | *Felis catus* |
| 0.0703 | 0.0841 | 0.0683 | 0.127 | *Tupaia belangeri* |
| 0.0829 | 0.127 | 0.107 | 0.112 | *Oryctolagus cuniculus* |
| 0.336 | 0.364 | 0.28 | 0.398 | *Ochotona princeps* |
| 0.0429 | 0.0489 | 0.0222 | 0.0388 | *Marmota marmota marmota* |
| 0.101 | 0.102 | 0.131 | 0.136 | *Ictidomys tridecemlineatus* |
| 0.352 | 0.5 | 0.416 | 0.447 | *Octodon degus* |
| 0.158 | 0.143 | 0.195 | 0.183 | *Chinchilla lanigera* |
| 0.297 | 0.39 | 0.357 | 0.283 | *Cavia porcellus* |
| 0.125 | 0.114 | 0.069 | 0.116 | *Heterocephalus glaber* |
| 0.136 | 0.193 | 0.139 | 0.18 | *Jaculus jaculus* |
| 0.362 | 0.466 | 0.366 | 0.401 | *Rattus norvegicus* |
| 0.378 | 0.383 | 0.409 | 0.451 | *Mus musculus* |
| 0.291 | 0.256 | 0.271 | 0.443 | *Meriones unguiculatus* |
| 0.273 | 0.186 | 0.224 | 0.343 | *Peromyscus maniculatus* |
| 0.435 | 0.633 | 0.473 | 0.528 | *Microtus ochrogaster* |
| 0.362 | 0.561 | 0.541 | 0.538 | *Mesocricetus auratus* |
| 0.225 | 0.221 | 0.181 | 0.336 | *Dipodomys ordii* |
| 0.0708 | 0.0903 | 0.0716 | 0.0659 | *Castor canadensis* |
| 0.103 | 0.197 | 0.0849 | 0.151 | *Otolemur garnettii* |
| 0.0403 | 0.0785 | 0.0421 | 0.0504 | *Propithecus coquereli* |
| 0.101 | 0.0509 | 0.0524 | 0.144 | *Microcebus murinus* |
| 0.0229 | 0.0197 | 0.0232 | 0.0201 | *Nomascus leucogenys* |
| 0.0231 | 0.0187 | 0.0112 | 0.0145 | *Pongo abelii* |
| 0.0239 | 0.0992 | 0.0197 | 0.0292 | *Pan troglodytes* |
| 0.0154 | 0.0131 | 0.0102 | 0.0131 | *Homo sapiens* |
| 0.0222 | 0.0158 | 0.00956 | 0.0145 | *Gorilla gorilla* |
| 0.0392 | 0.0311 | 0.018 | 0.0295 | *Chlorocebus sabaeus* |
| 0.0314 | 0.029 | 0.0164 | 0.0242 | *Macaca mulatta* |
| 0.0211 | 0.0224 | 0.0137 | 0.0201 | *Papio anubis* |
| 0.0245 | 0.0231 | 0.0138 | 0.0232 | *Mandrillus leucophaeus* |
| 0.0269 | 0.0343 | 0.209 | 0.0233 | *Cercocebus atys* |
| 0.0291 | 0.0259 | 0.014 | 0.0208 | *Colobus angolensis* |
| 0.0581 | 0.0503 | 0.0673 | 0.117 | *Saimiri boliviensis* |
| 0.0445 | 0.0498 | 0.038 | 0.039 | *Cebus capucinus* |
| 0.124 | 0.0856 | 0.0996 | 0.114 | *Callithrix jacchus* |
| 0.0472 | 0.0469 | 0.0366 | 0.0677 | *Aotus nancymaae* |
| **34.9** | **70.1** | **88.6** | **151** | **Maximum range** |

Table 8: Repeatability of mutation rate ($\mu$) estimation in mammals, for the extant taxa.

| Correlation ($\rho$) | $N_e$ | $\mu$ | Maximum longevity | Adult weight | Female maturity |
|---|---|---|---|---|---|
| $N_e$ | - | $0.439^{**}$ | $-0.523^{**}$ | $-0.544^{**}$ | $-0.47^{**}$ |
| $\mu$ | - | - | $-0.832^{**}$ | $-0.835^{**}$ | $-0.833^{**}$ |
| Maximum longevity | - | - | - | $0.827^{**}$ | $0.845^{**}$ |
| Adult weight | - | - | - | - | $0.809^{**}$ |
| Female maturity | - | - | - | - | - |

| Correlation ($\rho$) | $N_e$ | $\mu$ | Maximum longevity | Adult weight | Female maturity |
|---|---|---|---|---|---|
| $N_e$ | - | $0.51^{**}$ | $-0.591^{**}$ | $-0.496^{**}$ | $-0.465^{**}$ |
| $\mu$ | - | - | $-0.771^{**}$ | $-0.722^{**}$ | $-0.679^{**}$ |
| Maximum longevity | - | - | - | $0.802^{**}$ | $0.812^{**}$ |
| Adult weight | - | - | - | - | $0.764^{**}$ |
| Female maturity | - | - | - | - | - |

| Correlation ($\rho$) | $N_e$ | $\mu$ | Maximum longevity | Adult weight | Female maturity |
|---|---|---|---|---|---|
| $N_e$ | - | $0.497^{**}$ | $-0.643^{**}$ | $-0.577^{**}$ | $-0.627^{**}$ |
| $\mu$ | - | - | $-0.803^{**}$ | $-0.795^{**}$ | $-0.739^{**}$ |
| Maximum longevity | - | - | - | $0.836^{**}$ | $0.843^{**}$ |
| Adult weight | - | - | - | - | $0.805^{**}$ |
| Female maturity | - | - | - | - | - |

| Correlation ($\rho$) | $N_e$ | $\mu$ | Maximum longevity | Adult weight | Female maturity |
|---|---|---|---|---|---|
| $N_e$ | - | $0.707^{**}$ | $-0.687^{**}$ | $-0.638^{**}$ | $-0.611^{**}$ |
| $\mu$ | - | - | $-0.85^{**}$ | $-0.865^{**}$ | $-0.83^{**}$ |
| Maximum longevity | - | - | - | $0.839^{**}$ | $0.851^{**}$ |
| Adult weight | - | - | - | - | $0.817^{**}$ |
| Female maturity | - | - | - | - | - |

Table 9: In all four replicates, covariance coefficient between effective population size ($N_e$), mutation rate per site per unit of time ($\mu$), and life-history traits (maximum longevity, adult weight and female maturity) were computed in placental mammals. Asterisks indicate strength of support ($^*pp > 0.95$, $^{**}pp > 0.975$).

#### 3.4 Amino-acid preferences entropy

| **Experiment** | $\langle\Omega\rangle$ (**branch** $N_\mathrm{e}$) | $\langle\Omega\rangle$ (**constant** $N_\mathrm{e}$) |
|---|---|---|
| Mammals 18 CDS, replicate 1, Chain 1 | $1.07 \pm 0.10$ | $1.14 \pm 0.10$ |
| Mammals 18 CDS, replicate 2, Chain 2 | $1.07 \pm 0.09$ | $1.14 \pm 0.10$ |
| Mammals 18 CDS, replicate 2, Chain 1 | $1.06 \pm 0.10$ | $1.12 \pm 0.09$ |
| Mammals 18 CDS, replicate 2, Chain 2 | $1.06 \pm 0.09$ | $1.11 \pm 0.10$ |
| Mammals 18 CDS, replicate 3, Chain 1 | $1.08 \pm 0.12$ | $1.15 \pm 0.11$ |
| Mammals 18 CDS, replicate 3, Chain 2 | $1.04 \pm 0.10$ | $1.18 \pm 0.11$ |
| Mammals 18 CDS, replicate 4, Chain 1 | $0.94 \pm 0.11$ | $1.02 \pm 0.12$ |
| Mammals 18 CDS, replicate 4, Chain 2 | $0.89 \pm 0.11$ | $1.02 \pm 0.11$ |
| Mammals 36 CDS, replicate 1, Chain 1 | $1.02 \pm 0.06$ | $1.07 \pm 0.10$ |
| Mammals 36 CDS, replicate 1, Chain 2 | $0.91 \pm 0.07$ | $1.03 \pm 0.07$ |
| Mammals 36 CDS, replicate 2, Chain 1 | $0.92 \pm 0.09$ | $0.96 \pm 0.09$ |
| Mammals 36 CDS, replicate 2, Chain 2 | $1.01 \pm 0.09$ | $1.02 \pm 0.11$ |
| Mammals 36 CDS, replicate 3, Chain 1 | $0.93 \pm 0.00$ | $1.05 \pm 0.09$ |
| Mammals 36 CDS, replicate 3, Chain 2 | $1.02 \pm 0.07$ | $1.05 \pm 0.11$ |
| Mammals 36 CDS, replicate 4, Chain 1 | $1.04 \pm 0.07$ | $1.10 \pm 0.08$ |
| Mammals 36 CDS, replicate 4, Chain 2 | $1.03 \pm 0.10$ | $1.08 \pm 0.08$ |
| Mammals 36 CDS, replicate 5, Chain 1 | $1.03 \pm 0.10$ | $1.03 \pm 0.08$ |
| Mammals 36 CDS, replicate 5, Chain 2 | $0.99 \pm 0.10$ | $1.04 \pm 0.08$ |
| Mammals 36 CDS, replicate 6, Chain 1 | $1.05 \pm 0.10$ | $1.10 \pm 0.08$ |
| Mammals 36 CDS, replicate 6, Chain 2 | $0.97 \pm 0.11$ | $1.10 \pm 0.10$ |

Table 10: Estimated amino-acid entropy in mammals. Obtained with the mechanistic inference model developed in this paper of site-specific amino-acid fitness profiles and log-Brownian process for $N_\mathrm{e}$, $\mu$ and life-history traits (in the left column), or under the assumption of constant $N_\mathrm{e}$ (in the right column).

#### 3.5 Traits estimation with branch $\omega$ (replicate 1, chain 1)

Obtained with the phenomenological inference model of log-Brownian process for the $\mu$ and the relative non-synonymous substitution rate ($\omega$), as in Lartillot and Poujol (2011).

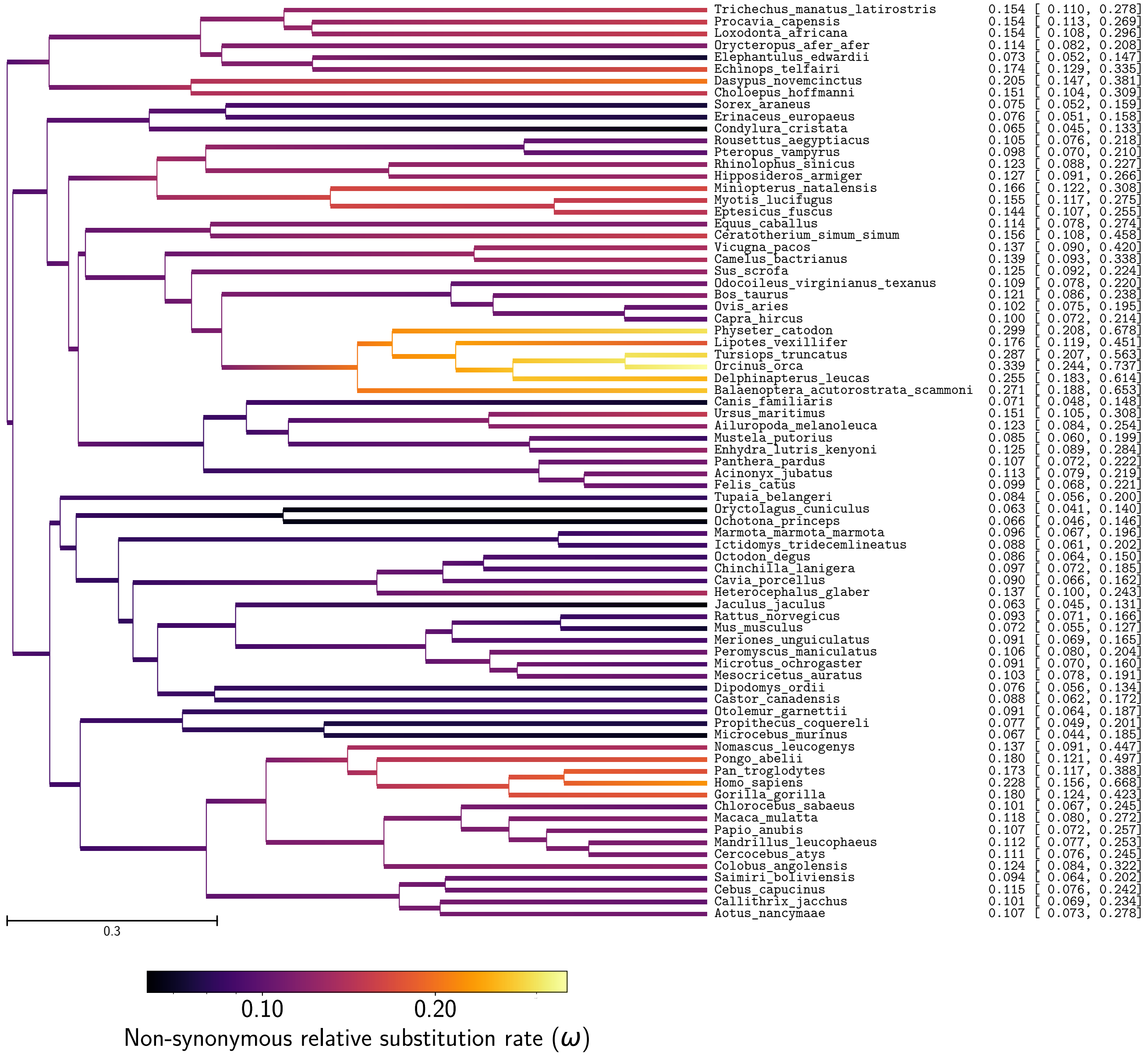


Figure 17: Non-synonymous substitution rate ($\omega$) estimation in mammals

| **Correlation ($\rho$)** | $\omega$ | $\mu$ | **Maximum longevity** | **Adult weight** | **Female maturity** |
|---|---|---|---|---|---|
| $\omega$ | - | $-0.374^{**}$ | $0.544^{**}$ | $0.43^{**}$ | $0.433^{**}$ |
| $\mu$ | - | - | $-0.807^{**}$ | $-0.781^{**}$ | $-0.824^{**}$ |
| **Maximum longevity** | - | - | - | $0.801^{**}$ | $0.83^{**}$ |
| **Adult weight** | - | - | - | - | $0.785^{**}$ |
| **Female maturity** | - | - | - | - | - |

Table 11: Correlation coefficient between non-synonymous substitution rate ($\omega$), mutation rate per site per unit of time ($\mu$), and life-history traits (maximum longevity, adult weight and female maturity) were computed in placental mammals. Asterisks indicate strength of support ($^{*}pp > 0.95$, $^{**}pp > 0.975$).

| Covariance ($\Sigma$) | $\omega$ | $\mu$ | Maximum longevity | Adult weight | Female maturity |
|---|---|---|---|---|---|
| $\omega$ | $0.215^{**}$ | $-0.236^{**}$ | $0.231^{**}$ | $0.828^{**}$ | $0.242^{**}$ |
| $\mu$ | - | $1.82^{**}$ | $-0.998^{**}$ | $-4.38^{**}$ | $-1.34^{**}$ |
| Maximum longevity | - | - | $0.837^{**}$ | $3.04^{**}$ | $0.917^{**}$ |
| Adult weight | - | - | - | $17.1^{**}$ | $3.93^{**}$ |
| Female maturity | - | - | - | - | $1.45^{**}$ |

Table 12: Correlation coefficient between non-synonymous substitution rate ($\omega$), mutation rate per site per unit of time ($\mu$), and life-history traits (maximum longevity, adult weight and female maturity) were computed in placental mammals. Asterisks indicate strength of support ($^{*}pp > 0.95$, $^{**}pp > 0.975$).

| Partial coefficient | $\omega$ | $\mu$ | Maximum longevity | Adult weight | Female maturity |
|---|---|---|---|---|---|
| $\omega$ | - | 0.15 | $0.369^{**}$ | 0.0468 | 0.0223 |
| $\mu$ | - | - | $-0.299^{*}$ | $-0.272$ | $-0.382^{**}$ |
| Maximum longevity | - | - | - | $0.283^{**}$ | $0.338^{**}$ |
| Adult weight | - | - | - | - | $0.21^{*}$ |
| Female maturity | - | - | - | - | - |

Table 13: Partial correlation coefficient between non-synonymous substitution rate ($\omega$), mutation rate per site per unit of time ($\mu$), and life-history traits (maximum longevity, adult weight and female maturity) were computed in placental mammals. Asterisks indicate strength of support ($^{*}pp > 0.95$, $^{**}pp > 0.975$).

### 4 Empirical data in Isopods

#### 4.1 Traits estimation (replicate 1, chain 1)

Obtained with the mechanistic inference model developed in this paper of site-specific amino-acid fitness profiles and log-Brownian process for $N_{\mathrm{e}}$, $\mu$ and life-history traits.

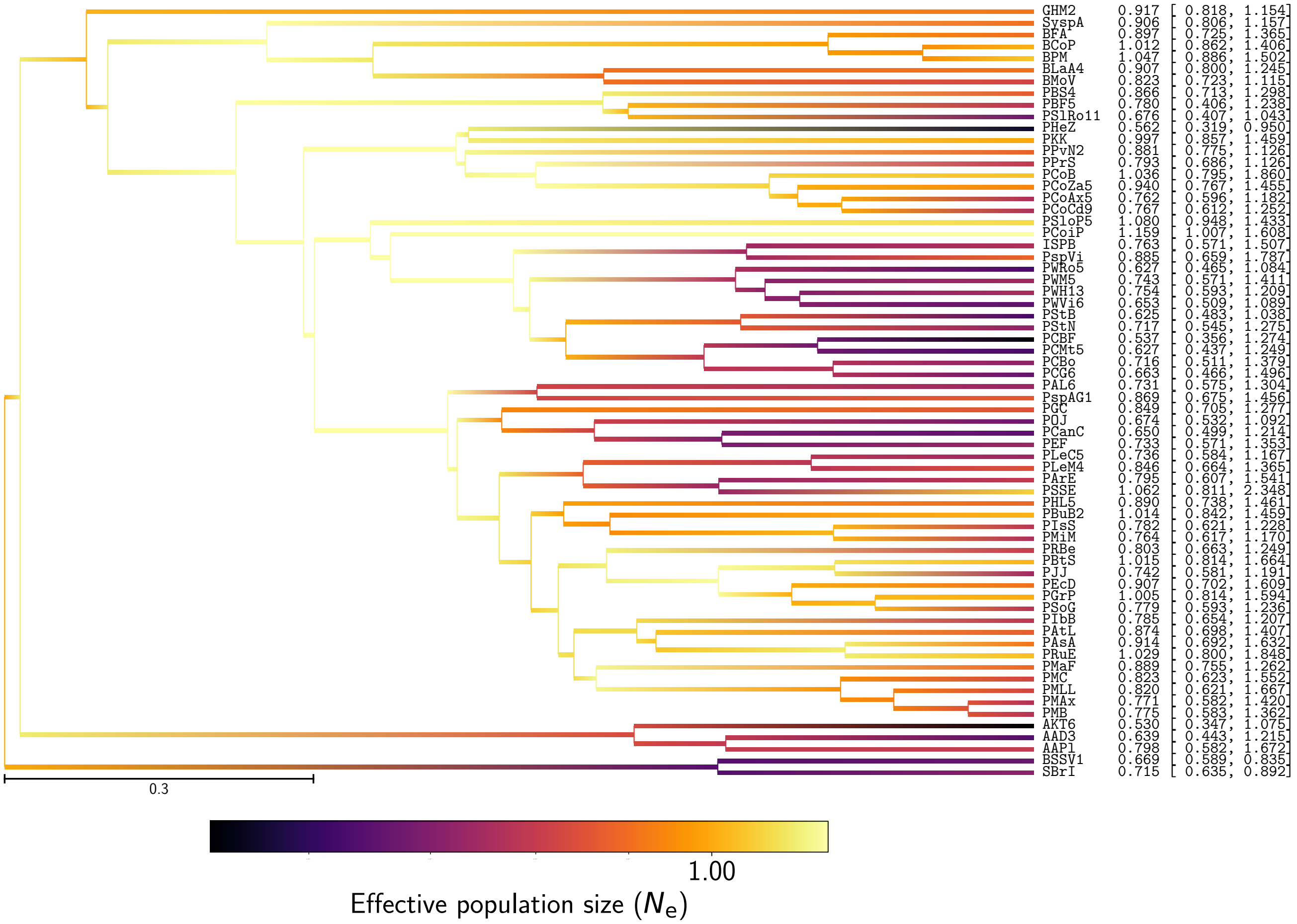


Figure 18: Effective population size ($N_{\mathrm{e}}$) estimation in isopods

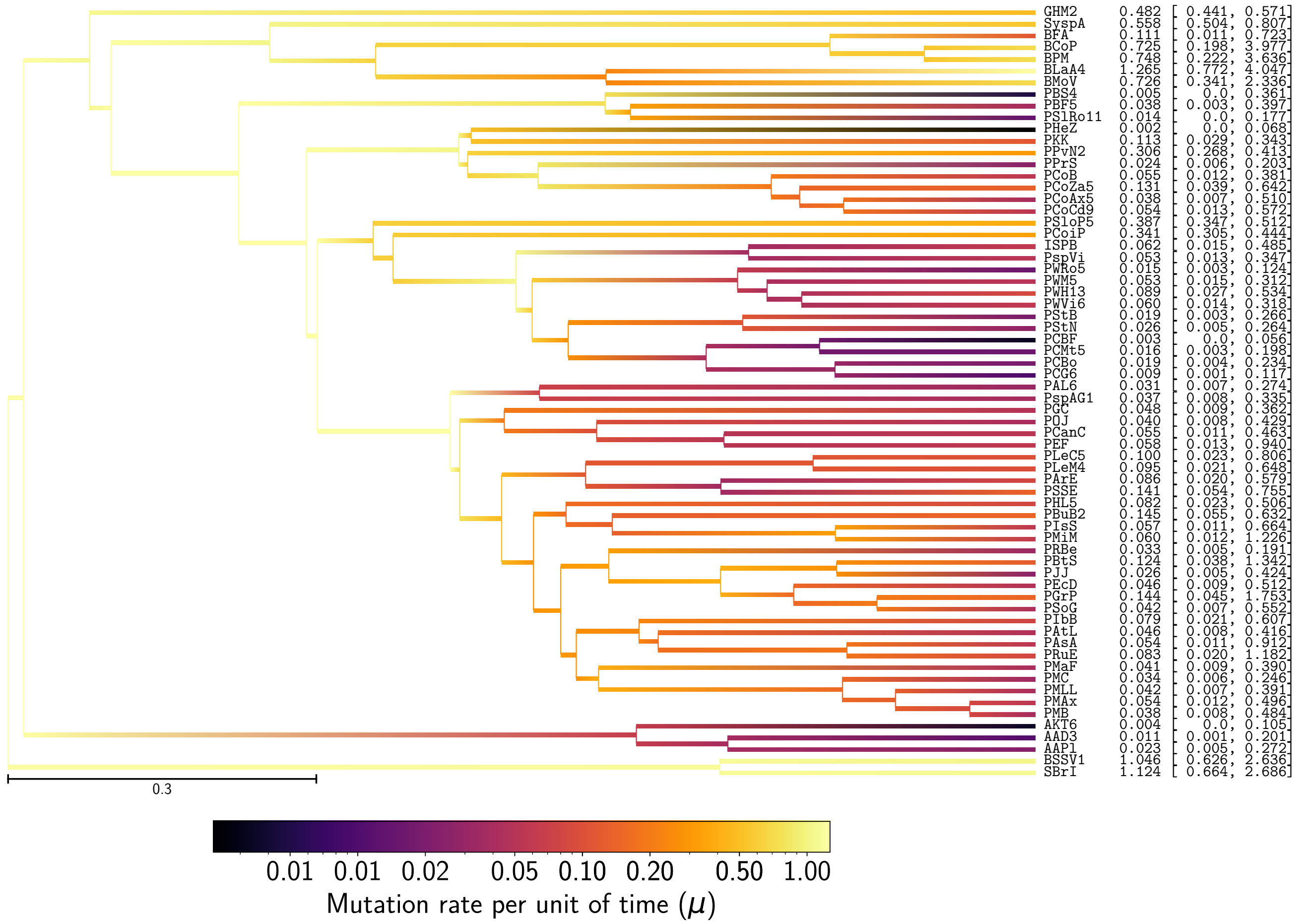


Figure 19: Mutation rate ($\mu$) estimation in isopods

#### 4.2 Repeatability of experiments

6 independent inferences were performed on a randomly chosen set of 12 coding sequences (CDS) out of 135. Obtained with the mechanistic inference model developed in this paper of site-specific amino-acid fitness profiles and log-Brownian process for $N_e$, $\mu$. Each plot is a correlation between a pair of experiments for a given parameter. For each node (or branch) of the tree, the mean posterior of the parameter over the MCMC (after burn-in) is represented in blue dots, green solid lines are the 90% confidence interval of the MCMC. Solid red line is the regression line between replicates.

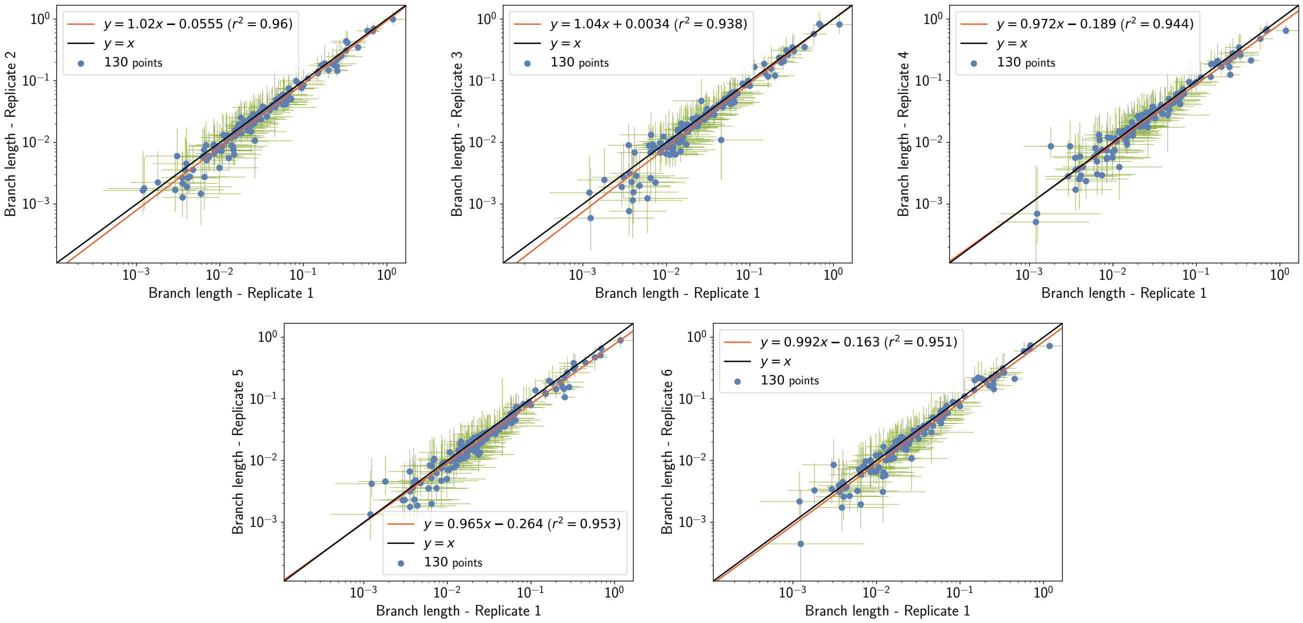


Figure 20: Repeatability of branch length ($l$) estimation in isopods

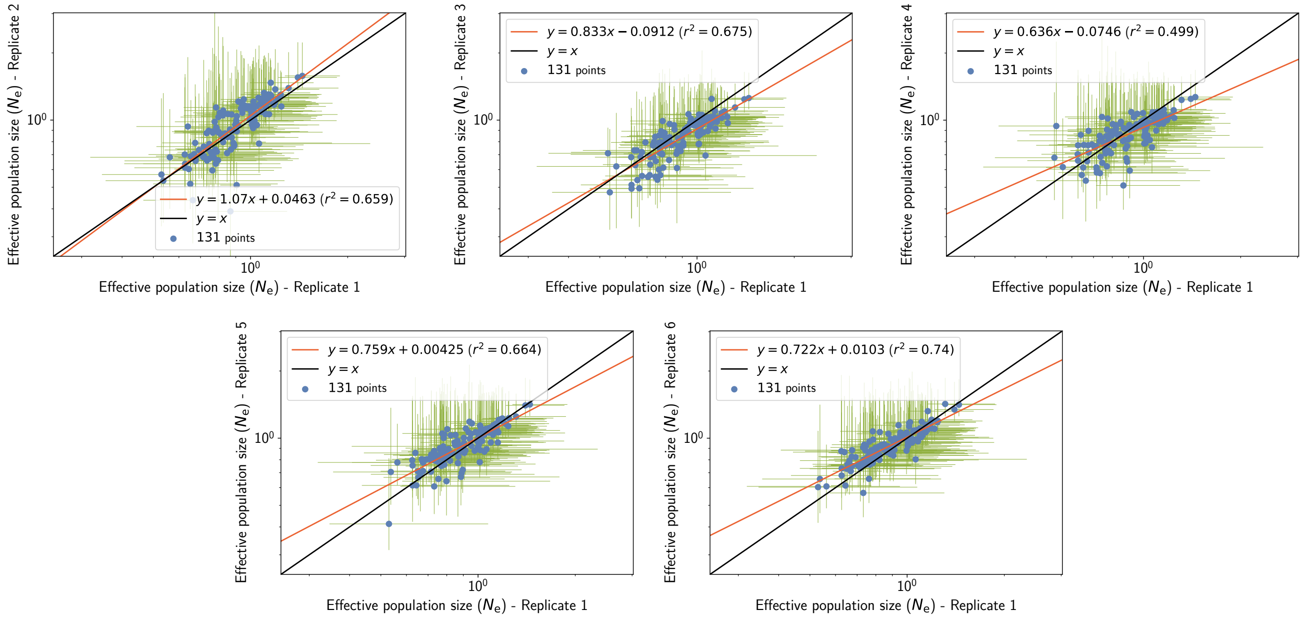


Figure 21: Repeatability of effective population size ($N_e$) estimation in isopods

| Rep. 1 | Rep. 2 | Rep. 3 | Rep. 4 | Rep. 5 | Rep. 6 | Habitat | Pigmentation | Ocular structure | Code | Taxon |
|---|---|---|---|---|---|---|---|---|---|---|
| 0.917 | 1.21 | 0.91 | 1.09 | 1.1 | 0.997 | Underground | Depigmented | Anophthalmia | GHM2 | *Gallasellus heyli* |
| 0.906 | 0.511 | 0.876 | 0.907 | 0.948 | 0.964 | Underground | Depigmented | Anophthalmia | SyspA | *Synasellus sp* |
| 0.897 | 0.954 | 0.838 | 0.819 | 0.721 | 0.859 | Underground | Depigmented | Anophthalmia | BFA | *Bragasellus frontellum* |
| 1.01 | 1.15 | 0.78 | 0.894 | 0.78 | 0.945 | Surface | Pigmented | Ocular | BCoP | *Bragasellus cortesi* |
| 1.05 | 0.95 | 0.881 | 0.815 | 0.87 | 1.08 | Surface | Pigmented | Ocular | BPM | *Bragasellus peltatus* |
| 0.907 | 0.891 | 0.769 | 0.831 | 0.873 | 0.907 | Underground | Depigmented | Anophthalmia | BLaA4 | - |
| 0.823 | 0.797 | 0.839 | 0.949 | 0.852 | 1.06 | Underground | Depigmented | Anophthalmia | BMoV | *Bragasellus molinai* |
| 0.866 | 0.39 | 0.61 | 0.51 | 0.773 | 0.871 | Underground | Depigmented | Anophthalmia | PBS4 | *Proasellus boui* |
| 0.78 | 0.637 | 0.702 | 0.808 | 0.919 | 0.884 | Underground | Depigmented | Anophthalmia | PBF5 | *Proasellus boui* |
| 0.676 | 0.657 | 0.718 | 0.895 | 0.777 | 0.884 | Underground | Depigmented | Anophthalmia | PSlRo11 | *Proasellus slavus* |
| 0.562 | 0.681 | 0.62 | 0.616 | 0.778 | 0.608 | Underground | Depigmented | Anophthalmia | PHeZ | *Proasellus hercegovinensis* |
| 0.997 | 1.15 | 0.962 | 1.08 | 0.907 | 1.06 | Surface | Pigmented | Ocular | PKK | *Proasellus karamani* |
| 0.881 | 0.897 | 0.753 | 0.746 | 0.877 | 0.901 | Underground | Depigmented | Anophthalmia | PPvN2 | *Proasellus pavani* |
| 0.793 | 0.66 | 0.876 | 0.711 | 0.813 | 0.808 | Underground | Depigmented | Anophthalmia | PPrS | *Proasellus parvulus* |
| 1.04 | 1.06 | 1.06 | 0.968 | 1.05 | 0.957 | Surface | Pigmented | Ocular | PCoB | *Proasellus coxalis* |
| 0.94 | 1.17 | 1.06 | 0.937 | 1.18 | 1.14 | Surface | Pigmented | Ocular | PCoZa5 | *Proasellus coxalis* |
| 0.762 | 0.595 | 0.792 | 0.731 | 0.902 | 0.657 | Underground | Depigmented | Anophthalmia | PCoAx5 | *Proasellus coxalis* |
| 0.767 | 0.705 | 0.711 | 0.848 | 0.847 | 0.761 | Underground | Depigmented | Microphthalmia | PCoCd9 | *Proasellus coxalis* |
| 1.08 | 1.09 | 0.893 | 0.975 | 1.1 | 0.968 | Underground | Depigmented | Anophthalmia | PSloP5 | *Proasellus slovenicus* |
| 1.16 | 1.28 | 1.02 | 1.07 | 1.21 | 1.32 | Surface | Pigmented | Ocular | PCoiP | *Proasellus coiffaiti* |
| 0.763 | 0.888 | 0.753 | 0.709 | 0.766 | 0.86 | Underground | Depigmented | Anophthalmia | ISPB | *Proasellus nsp* |
| 0.885 | 0.789 | 0.765 | 0.675 | 0.67 | 0.857 | Underground | Depigmented | Anophthalmia | PspVi | *Proasellus nsp* |
| 0.627 | 0.636 | 0.493 | 0.771 | 0.754 | 0.732 | Underground | Depigmented | Anophthalmia | PWRo5 | *Proasellus walteri* |
| 0.743 | 0.671 | 0.558 | 1.03 | 0.834 | 0.761 | Underground | Depigmented | Anophthalmia | PWM5 | *Proasellus walteri* |
| 0.754 | 0.718 | 0.54 | 0.79 | 0.656 | 0.824 | Underground | Depigmented | Anophthalmia | PWH13 | *Proasellus walteri* |
| 0.653 | 0.689 | 0.538 | 0.774 | 0.667 | 0.764 | Underground | Depigmented | Anophthalmia | PWVi6 | *Proasellus walteri* |
| 0.625 | 0.701 | 0.682 | 0.703 | 0.796 | 0.875 | Underground | Depigmented | Anophthalmia | PStB | *Proasellus strouhali* |
| 0.717 | 0.693 | 0.682 | 0.582 | 0.839 | 0.734 | Underground | Depigmented | Anophthalmia | PStN | *Proasellus strouhali* |
| 0.537 | 0.535 | 0.475 | 0.941 | 0.705 | 0.656 | Underground | Depigmented | Anophthalmia | PCBF | *Proasellus cavaticus* |
| 0.627 | 0.611 | 0.512 | 0.659 | 0.614 | 0.68 | Underground | Depigmented | Anophthalmia | PCMt5 | *Proasellus cavaticus* |
| 0.716 | 0.761 | 0.632 | 0.603 | 0.844 | 0.747 | Underground | Depigmented | Anophthalmia | PCBo | *Proasellus cavaticus* |
| 0.663 | 0.437 | 0.495 | 0.535 | 0.689 | 0.793 | Underground | Depigmented | Anophthalmia | PCG6 | *Proasellus cavaticus* |
| 0.731 | 0.668 | 0.778 | 0.805 | 0.608 | 0.568 | Underground | Depigmented | Anophthalmia | PAL6 | *Proasellus albigensis* |
| 0.869 | 0.737 | 0.729 | 0.92 | 0.827 | 0.896 | Underground | Depigmented | Anophthalmia | PspAG1 | *Proasellus n* |
| 0.849 | 0.839 | 0.95 | 0.966 | 0.931 | 0.94 | Underground | Depigmented | Anophthalmia | PGC | *Proasellus grafi* |
| 0.674 | 0.79 | 0.719 | 0.772 | 0.784 | 0.826 | Surface | Part. dep. | Microphthalmia | POJ | *Proasellus ortizi* |
| 0.65 | 0.517 | 0.732 | 0.613 | 0.711 | 0.729 | Underground | Depigmented | Anophthalmia | PCanC | *Proasellus cantabricus* |
| 0.733 | 0.659 | 0.684 | 0.579 | 0.795 | 0.802 | Surface | Part. dep. | Microphthalmia | PEF | *Proasellus ebrensis* |
| 0.736 | 0.743 | 0.8 | 0.685 | 0.805 | 0.935 | Underground | Depigmented | Anophthalmia | PLeC5 | - |
| 0.846 | 0.711 | 0.785 | 0.864 | 0.957 | 0.889 | Underground | Depigmented | Anophthalmia | PLeM4 | - |
| 0.795 | 1.03 | 0.846 | 0.87 | 0.869 | 0.851 | Surface | Part. dep. | Microphthalmia | PArE | *Proasellus aragonensis* |
| 1.06 | 0.785 | 0.694 | 0.756 | 0.893 | 0.804 | Underground | Depigmented | Anophthalmia | PSSE | *Proasellus spelaeus* |
| 0.89 | 0.774 | 0.727 | 0.766 | 0.926 | 0.992 | Underground | Depigmented | Anophthalmia | PHL5 | - |
| 1.01 | 0.994 | 0.783 | 0.789 | 1.14 | 1.17 | Underground | Depigmented | Anophthalmia | PBuB2 | - |
| 0.782 | 1.14 | 0.991 | 0.853 | 1 | 1.07 | Surface | Pigmented | Ocular | PIsS | *Proasellus istrianus* |
| 0.764 | 0.878 | 0.853 | 0.736 | 0.887 | 0.966 | Surface | Pigmented | Ocular | PMiM | *Proasellus micropectinatus* |
| 0.803 | 1.08 | 0.819 | 0.96 | 1.1 | 0.823 | Surface | Part. dep. | Microphthalmia | PRBe | *Proasellus racovitzai* |
| 1.01 | 1.1 | 0.905 | 0.931 | 0.884 | 1.01 | Surface | Pigmented | Ocular | PBtS | *Proasellus beticus* |
| 0.742 | 0.896 | 0.842 | 0.826 | 0.84 | 0.882 | Underground | Depigmented | Microphthalmia | PJJ | *Proasellus jaloniacus* |
| 0.907 | 1.04 | 0.792 | 0.594 | 0.859 | 0.836 | Underground | Depigmented | Anophthalmia | PEcD | *Proasellus escolai* |
| 1.01 | 1.02 | 0.86 | 0.786 | 1.06 | 0.922 | Surface | Part. dep. | Microphthalmia | PGrP | *Proasellus granadensis* |
| 0.779 | 0.738 | 0.707 | 0.731 | 0.812 | 0.808 | Underground | Depigmented | Anophthalmia | PSoG | *Proasellus solanasi* |
| 0.785 | 0.918 | 0.854 | 0.788 | 1.04 | 0.986 | Surface | Pigmented | Ocular | PIbB | *Proasellus ibericus* |
| 0.874 | 0.836 | 0.764 | 0.815 | 0.866 | 1.05 | Underground | Depigmented | Anophthalmia | PAtL | *Proasellus arthrodilus* |
| 0.914 | 0.951 | 0.888 | 0.834 | 0.881 | 0.886 | Surface | Part. dep. | Microphthalmia | PAsA | *Proasellus assaforensis* |
| 1.03 | 0.994 | 0.939 | 0.854 | 0.971 | 1 | Underground | Depigmented | Anophthalmia | PRuE | *Proasellus rectus* |
| 0.889 | 0.764 | 0.761 | 0.651 | 0.698 | 0.847 | Underground | Depigmented | Anophthalmia | PMaF | *Proasellus margalefi* |
| 0.823 | 1.1 | 0.961 | 0.914 | 1.03 | 0.838 | Surface | Pigmented | Ocular | PMC | *Proasellus meridianus* |
| 0.82 | 1.05 | 0.806 | 0.914 | 0.856 | 0.799 | Surface | Pigmented | Ocular | PMLL | *Proasellus meridianus* |
| 0.771 | 0.889 | 0.796 | 0.907 | 0.839 | 0.863 | Underground | Part. dep. | Microphthalmia | PMAx | *Proasellus meridianus* |
| 0.775 | 0.982 | 0.892 | 1.02 | 1.05 | 0.882 | Surface | Pigmented | Ocular | PMB | *Proasellus meridianus* |
| 0.53 | 0.57 | 0.71 | 0.679 | 0.413 | 0.603 | Underground | Depigmented | Anophthalmia | AKT6 | *Asellus kosswigi* |
| 0.639 | 0.936 | 0.732 | 0.859 | 0.859 | 0.866 | Surface | Pigmented | Ocular | AAD3 | *Asellus aquaticus* |
| 0.798 | 0.882 | 0.795 | 0.89 | 0.641 | 0.875 | Surface | Pigmented | Ocular | AAPl | *Asellus aquaticus* |
| 0.669 | 0.704 | 0.684 | 0.655 | 0.805 | 0.711 | Underground | Depigmented | Anophthalmia | BSSV1 | *Balkanostenasellus skopljensis* |
| 0.715 | 0.682 | 0.685 | 0.614 | 0.707 | 0.77 | Underground | Depigmented | Anophthalmia | SBrI | *Stenasellus breuili* |
| **2.19** | **3.29** | **2.24** | **2.13** | **2.93** | **2.33** | - | - | - | - | **Maximum range** |

Table 14: Repeatability of effective population size ($N_e$) estimation in isopods, for the extant taxa.

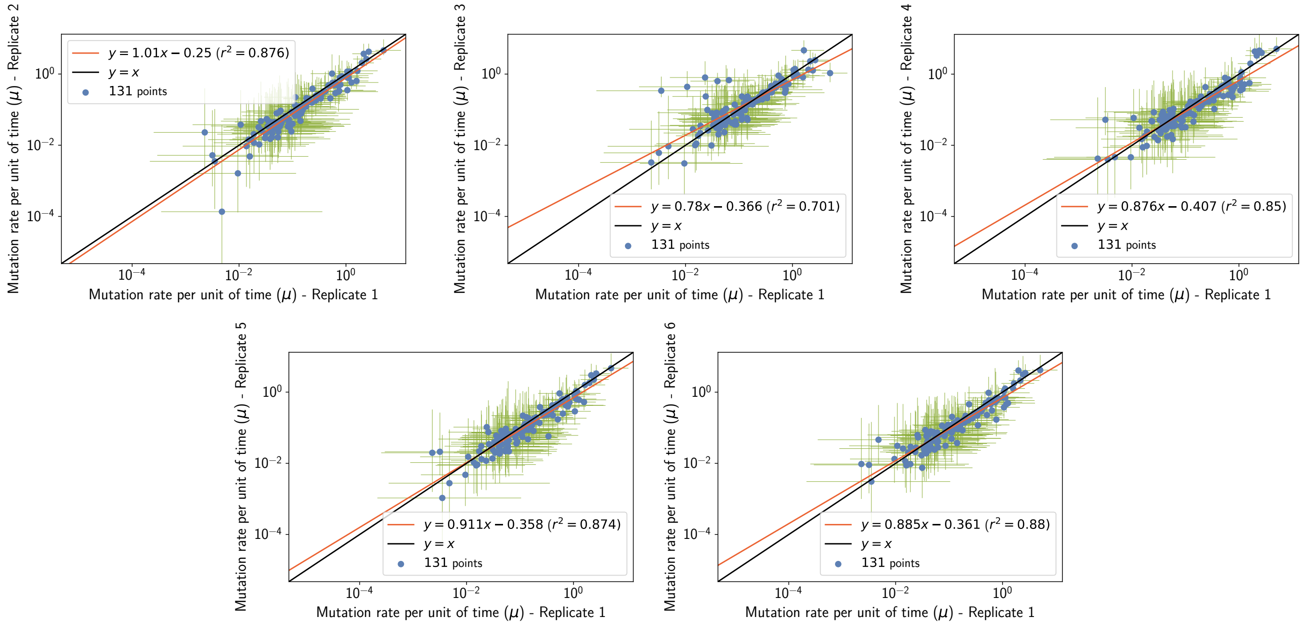


Figure 22: Repeatability of mutation rate ($\mu$) estimation in isopods

| Rep. 1 | Rep. 2 | Rep. 3 | Rep. 4 | Rep. 5 | Rep. 6 | Habitat | Pigmentation | Ocular structure | Code | Taxon |
|---|---|---|---|---|---|---|---|---|---|---|
| 0.482 | 0.488 | 0.54 | 0.462 | 0.456 | 0.497 | Underground | Depigmented | Anophthalmia | GHM2 | *Gallasellus heyli* |
| 0.558 | 0.563 | 0.536 | 0.52 | 0.503 | 0.59 | Underground | Depigmented | Anophthalmia | SyspA | *Synasellus sp* |
| 0.111 | 0.0674 | 0.0703 | 0.0781 | 0.0555 | 0.0852 | Underground | Depigmented | Anophthalmia | BFA | *Bragasellus frontellum* |
| 0.725 | 0.32 | 0.3 | 0.377 | 0.421 | 0.579 | Surface | Pigmented | Ocular | BCoP | *Bragasellus cortesi* |
| 0.748 | 0.337 | 0.364 | 0.331 | 0.427 | 0.692 | Surface | Pigmented | Ocular | BPM | *Bragasellus peltatus* |
| 1.27 | 0.579 | 0.893 | 0.516 | 0.607 | 0.49 | Underground | Depigmented | Anophthalmia | BLaA4 | - |
| 0.726 | 0.462 | 0.442 | 0.463 | 0.428 | 0.519 | Underground | Depigmented | Anophthalmia | BMoV | *Bragasellus molinai* |
| 0.00478 | 0.000136 | 0.00937 | 0.00462 | 0.00277 | 0.046 | Underground | Depigmented | Anophthalmia | PBS4 | *Proasellus boui* |
| 0.038 | 0.0347 | 0.0822 | 0.0873 | 0.048 | 0.0628 | Underground | Depigmented | Anophthalmia | PBF5 | *Proasellus boui* |
| 0.0137 | 0.00942 | 0.0281 | 0.031 | 0.0105 | 0.0259 | Underground | Depigmented | Anophthalmia | PSlRo11 | *Proasellus slavus* |
| 0.00228 | 0.0229 | 0.00329 | 0.00422 | 0.0197 | 0.00957 | Underground | Depigmented | Anophthalmia | PHeZ | *Proasellus hercegovinensis* |
| 0.113 | 0.224 | 0.235 | 0.103 | 0.218 | 0.202 | Surface | Pigmented | Ocular | PKK | *Proasellus karamani* |
| 0.306 | 0.25 | 0.257 | 0.225 | 0.229 | 0.23 | Underground | Depigmented | Anophthalmia | PPvN2 | *Proasellus pavani* |
| 0.0238 | 0.0105 | 0.235 | 0.0472 | 0.104 | 0.0275 | Underground | Depigmented | Anophthalmia | PPrS | *Proasellus parvulus* |
| 0.0545 | 0.0359 | 0.0712 | 0.0458 | 0.073 | 0.0453 | Surface | Pigmented | Ocular | PCoB | *Proasellus coxalis* |
| 0.131 | 0.121 | 0.213 | 0.149 | 0.198 | 0.141 | Surface | Pigmented | Ocular | PCoZa5 | *Proasellus coxalis* |
| 0.0381 | 0.015 | 0.0518 | 0.0305 | 0.0522 | 0.0356 | Underground | Depigmented | Anophthalmia | PCoAx5 | *Proasellus coxalis* |
| 0.0544 | 0.0332 | 0.0673 | 0.0728 | 0.0931 | 0.0672 | Underground | Depigmented | Microphthalmia | PCoCd9 | *Proasellus coxalis* |
| 0.387 | 0.328 | 0.364 | 0.348 | 0.3 | 0.317 | Underground | Depigmented | Anophthalmia | PSloP5 | *Proasellus slovenicus* |
| 0.341 | 0.28 | 0.296 | 0.246 | 0.0573 | 0.253 | Surface | Pigmented | Ocular | PCoiP | *Proasellus coiffaiti* |
| 0.062 | 0.0427 | 0.0495 | 0.0556 | 0.0217 | 0.0281 | Underground | Depigmented | Anophthalmia | ISPB | *Proasellus nsp* |
| 0.0533 | 0.0419 | 0.0549 | 0.0516 | 0.0228 | 0.0245 | Underground | Depigmented | Anophthalmia | PspVi | *Proasellus nsp* |
| 0.0151 | 0.0144 | 0.0101 | 0.015 | 0.00874 | 0.0115 | Underground | Depigmented | Anophthalmia | PWRo5 | *Proasellus walteri* |
| 0.0531 | 0.0167 | 0.0231 | 0.0911 | 0.0218 | 0.0243 | Underground | Depigmented | Anophthalmia | PWM5 | *Proasellus walteri* |
| 0.0886 | 0.0317 | 0.0384 | 0.0491 | 0.0402 | 0.0934 | Underground | Depigmented | Anophthalmia | PWH13 | *Proasellus walteri* |
| 0.0597 | 0.093 | 0.0527 | 0.106 | 0.0297 | 0.0383 | Underground | Depigmented | Anophthalmia | PWVi6 | *Proasellus walteri* |
| 0.019 | 0.0117 | 0.0353 | 0.0242 | 0.0131 | 0.0291 | Underground | Depigmented | Anophthalmia | PStB | *Proasellus strouhali* |
| 0.0263 | 0.0262 | 0.026 | 0.0173 | 0.0345 | 0.0534 | Underground | Depigmented | Anophthalmia | PStN | *Proasellus strouhali* |
| 0.00317 | 0.00521 | 0.00612 | 0.0523 | 0.0211 | 0.00902 | Underground | Depigmented | Anophthalmia | PCBF | *Proasellus cavaticus* |
| 0.0159 | 0.00485 | 0.0112 | 0.00879 | 0.00919 | 0.00876 | Underground | Depigmented | Anophthalmia | PCMt5 | *Proasellus cavaticus* |
| 0.0188 | 0.0205 | 0.0221 | 0.00963 | 0.0302 | 0.00951 | Underground | Depigmented | Anophthalmia | PCBo | *Proasellus cavaticus* |
| 0.0095 | 0.00162 | 0.00309 | 0.00461 | 0.00475 | 0.0132 | Underground | Depigmented | Anophthalmia | PCG6 | *Proasellus cavaticus* |
| 0.0314 | 0.021 | 0.0791 | 0.0263 | 0.0238 | 0.00749 | Underground | Depigmented | Anophthalmia | PAL6 | *Proasellus albigensis* |
| 0.0372 | 0.0313 | 0.073 | 0.0375 | 0.0464 | 0.0191 | Underground | Depigmented | Anophthalmia | PspAG1 | *Proasellus n* |
| 0.0477 | 0.0403 | 0.0817 | 0.0666 | 0.0594 | 0.0425 | Underground | Depigmented | Anophthalmia | PGC | *Proasellus grafi* |
| 0.0404 | 0.03 | 0.0467 | 0.0547 | 0.0285 | 0.0343 | Surface | Part. dep. | Microphthalmia | POJ | *Proasellus ortizi* |
| 0.0551 | 0.0385 | 0.0367 | 0.0343 | 0.0369 | 0.052 | Underground | Depigmented | Anophthalmia | PCanC | *Proasellus cantabricus* |
| 0.0578 | 0.0337 | 0.0446 | 0.0149 | 0.0349 | 0.0499 | Surface | Part. dep. | Microphthalmia | PEF | *Proasellus ebrensis* |
| 0.1 | 0.0399 | 0.0968 | 0.0322 | 0.0743 | 0.104 | Underground | Depigmented | Anophthalmia | PLeC5 | - |
| 0.095 | 0.0243 | 0.0793 | 0.0904 | 0.0706 | 0.12 | Underground | Depigmented | Anophthalmia | PLeM4 | - |
| 0.0856 | 0.0426 | 0.0472 | 0.0317 | 0.0361 | 0.0311 | Surface | Part. dep. | Microphthalmia | PArE | *Proasellus aragonensis* |
| 0.141 | 0.0482 | 0.0583 | 0.0461 | 0.0474 | 0.0372 | Underground | Depigmented | Anophthalmia | PSSE | *Proasellus spelaeus* |
| 0.0822 | 0.0338 | 0.0387 | 0.0355 | 0.0522 | 0.059 | Underground | Depigmented | Anophthalmia | PHL5 | - |
| 0.145 | 0.0777 | 0.052 | 0.0705 | 0.0868 | 0.132 | Underground | Depigmented | Anophthalmia | PBuB2 | - |
| 0.0573 | 0.0857 | 0.106 | 0.0794 | 0.0706 | 0.0643 | Surface | Pigmented | Ocular | PIsS | *Proasellus istrianus* |
| 0.0599 | 0.0387 | 0.0584 | 0.0471 | 0.0606 | 0.0728 | Surface | Pigmented | Ocular | PMiM | *Proasellus micropectinatus* |
| 0.0328 | 0.0543 | 0.0365 | 0.052 | 0.0391 | 0.0289 | Surface | Part. dep. | Microphthalmia | PRBe | *Proasellus racovitzai* |
| 0.124 | 0.0797 | 0.15 | 0.168 | 0.128 | 0.12 | Surface | Pigmented | Ocular | PBtS | *Proasellus beticus* |
| 0.0256 | 0.0475 | 0.0985 | 0.0846 | 0.0751 | 0.0621 | Underground | Depigmented | Microphthalmia | PJJ | *Proasellus jaloniacus* |
| 0.0455 | 0.03 | 0.0441 | 0.0175 | 0.0339 | 0.0279 | Underground | Depigmented | Anophthalmia | PEcD | *Proasellus escolai* |
| 0.144 | 0.102 | 0.234 | 0.163 | 0.185 | 0.161 | Surface | Part. dep. | Microphthalmia | PGrP | *Proasellus granadensis* |
| 0.0423 | 0.028 | 0.0529 | 0.0454 | 0.0455 | 0.0635 | Underground | Depigmented | Anophthalmia | PSoG | *Proasellus solanasi* |
| 0.0788 | 0.0552 | 0.0724 | 0.0547 | 0.0686 | 0.0646 | Surface | Pigmented | Ocular | PIbB | *Proasellus ibericus* |
| 0.0458 | 0.0447 | 0.0517 | 0.0443 | 0.0601 | 0.0791 | Underground | Depigmented | Anophthalmia | PAtL | *Proasellus arthrodilus* |
| 0.0543 | 0.0351 | 0.0573 | 0.0478 | 0.065 | 0.0503 | Surface | Part. dep. | Microphthalmia | PAsA | *Proasellus assaforensis* |
| 0.083 | 0.0536 | 0.139 | 0.0608 | 0.0918 | 0.0674 | Underground | Depigmented | Anophthalmia | PRuE | *Proasellus rectus* |
| 0.0413 | 0.0168 | 0.0202 | 0.0369 | 0.0292 | 0.0359 | Underground | Depigmented | Anophthalmia | PMaF | *Proasellus margalefi* |
| 0.0343 | 0.034 | 0.0559 | 0.0219 | 0.0199 | 0.0235 | Surface | Pigmented | Ocular | PMC | *Proasellus meridianus* |
| 0.0416 | 0.0215 | 0.0337 | 0.0312 | 0.0187 | 0.0231 | Surface | Pigmented | Ocular | PMLL | *Proasellus meridianus* |
| 0.0542 | 0.0241 | 0.044 | 0.0194 | 0.0247 | 0.0486 | Underground | Part. dep. | Microphthalmia | PMAx | *Proasellus meridianus* |
| 0.0385 | 0.0252 | 0.0807 | 0.0782 | 0.0541 | 0.06 | Surface | Pigmented | Ocular | PMB | *Proasellus meridianus* |
| 0.00351 | 0.00353 | 0.34 | 0.00401 | 0.00106 | 0.00307 | Underground | Depigmented | Anophthalmia | AKT6 | *Asellus kosswigi* |
| 0.0107 | 0.0378 | 0.438 | 0.0589 | 0.0152 | 0.031 | Surface | Pigmented | Ocular | AAD3 | *Asellus aquaticus* |
| 0.0232 | 0.0321 | 0.802 | 0.048 | 0.0118 | 0.0209 | Surface | Pigmented | Ocular | AAPl | *Asellus aquaticus* |
| 1.05 | 0.901 | 1.31 | 0.585 | 0.841 | 0.711 | Underground | Depigmented | Anophthalmia | BSSV1 | *Balkanostenasellus skopljensis* |
| 1.12 | 0.784 | 1.11 | 0.479 | 0.641 | 0.701 | Underground | Depigmented | Anophthalmia | SBrI | *Stenasellus breuili* |
| **554** | **6.63e+03** | **423** | **146** | **792** | **231** | - | - | - | - | **Maximum range** |

Table 15: Repeatability of mutation rate ($\mu$) estimation in isopods, for the extant taxa.

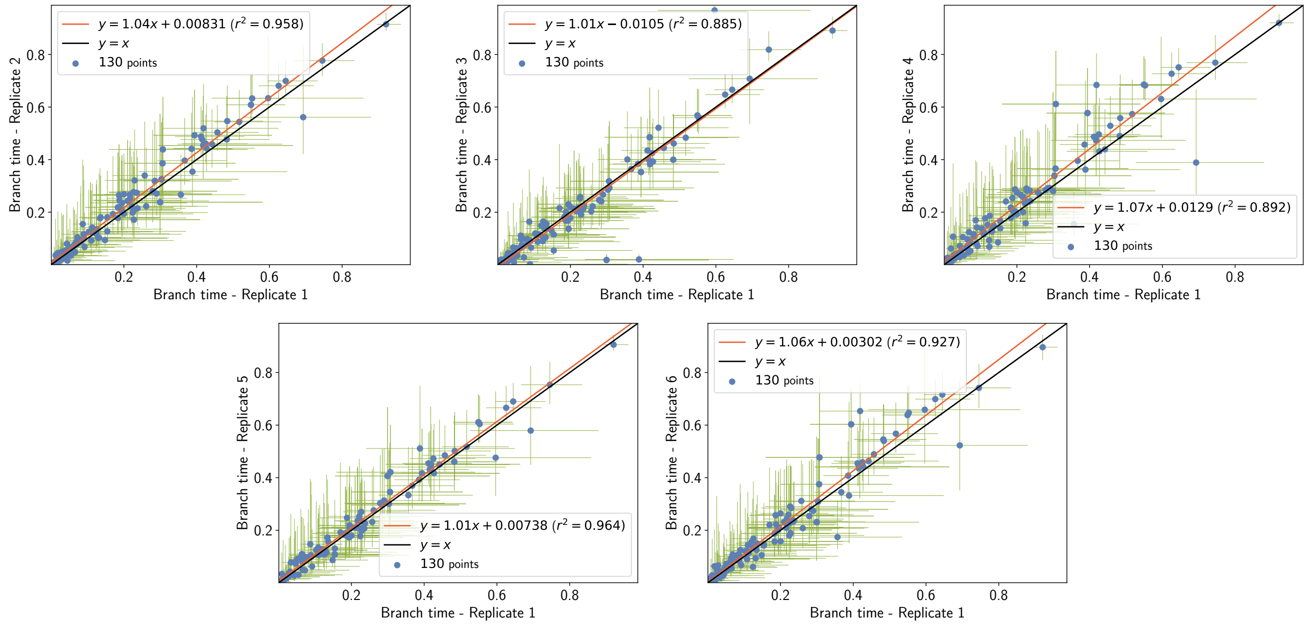


Figure 23: Repeatability of branch time ($\Delta T$) estimation in isopods

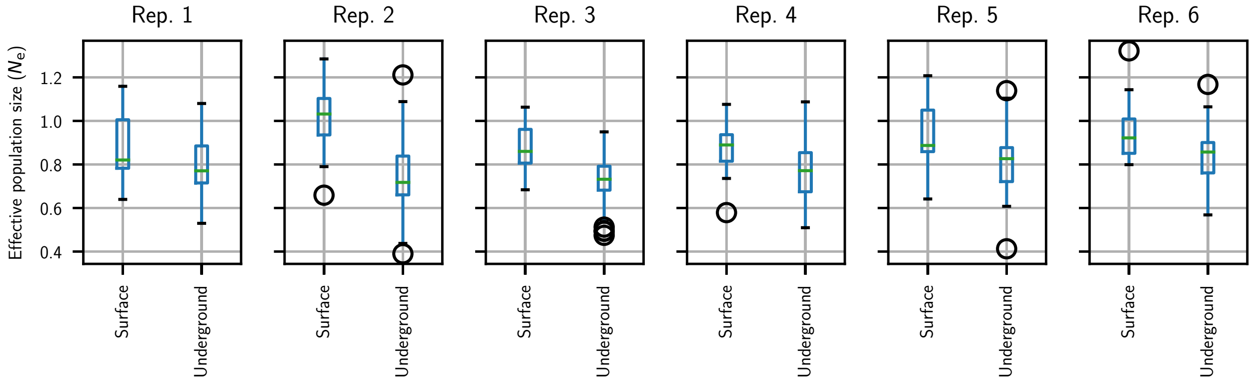


Figure 24: $N_{\mathrm{e}}$ as a function of habitat in isopods.

```
Analysis of Variance Table

Response: PopulationSize
           Df Sum Sq Mean Sq F value    Pr(>F)
Habitat     1 1.6777 1.67769  89.506 < 2.2e-16 ***
rep         5 0.4226 0.08452   4.509 0.0005236 ***
Residuals 389 7.2913 0.01874
---
Signif. codes:  0 ‘***’ 0.001 ‘**’ 0.01 ‘*’ 0.05 ‘.’ 0.1 ‘ ’ 1
```

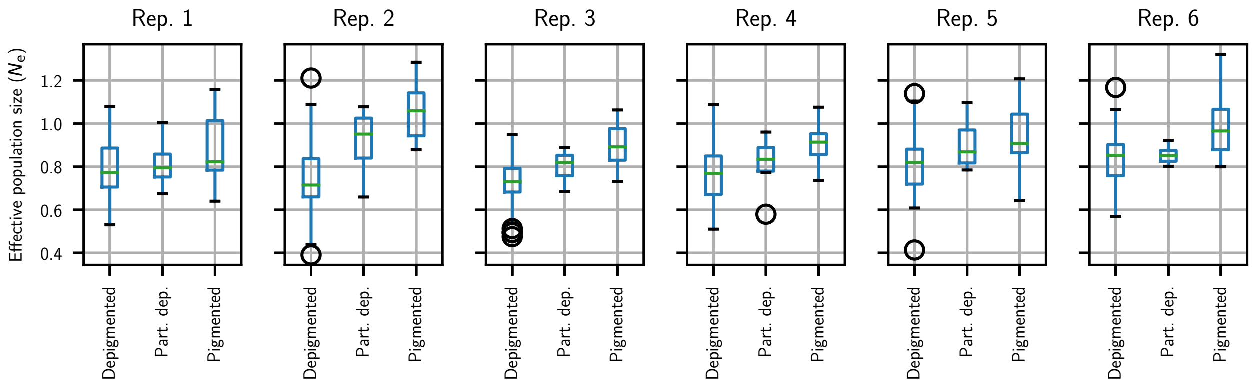


Figure 25: $N_\mathrm{e}$ as a function of pigmentation in isopods

```
Analysis of Variance Table

Response: PopulationSize
              Df Sum Sq Mean Sq F value    Pr(>F)
Pigmentation   2 1.9442 0.97210 53.6917 < 2.2e-16 ***
rep            5 0.4226 0.08452  4.6681 0.0003764 ***
Residuals    388 7.0248 0.01811
---
Signif. codes:  0 ‘***’ 0.001 ‘**’ 0.01 ‘*’ 0.05 ‘.’ 0.1 ‘ ’ 1
```

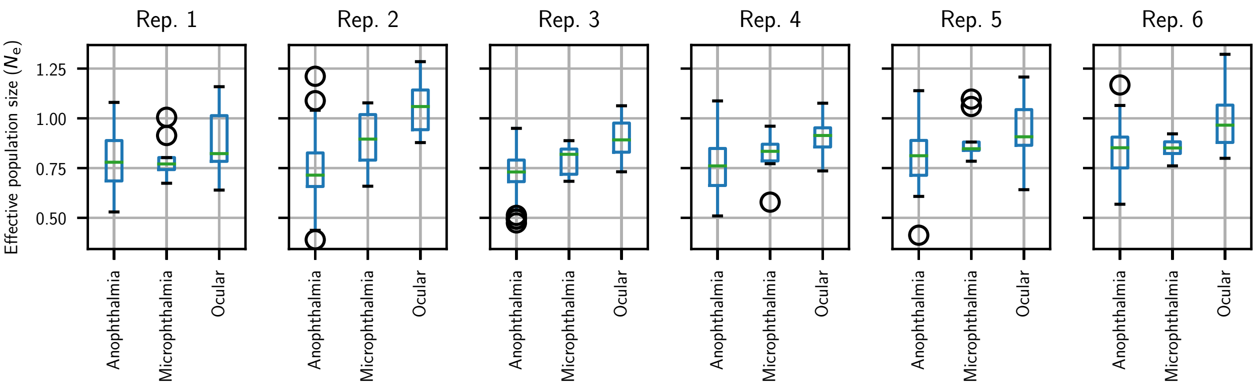


Figure 26: $N_\mathrm{e}$ as a function of ocular structure in isopods

```
Analysis of Variance Table

Response: PopulationSize
                  Df Sum Sq Mean Sq F value    Pr(>F)
Ocular.structure   2 1.9335 0.96676  53.316 < 2.2e-16 ***
rep                5 0.4226 0.08452   4.661  0.000382 ***
Residuals        388 7.0355 0.01813
---
Signif. codes:  0 ‘***’ 0.001 ‘**’ 0.01 ‘*’ 0.05 ‘.’ 0.1 ‘ ’ 1
```

### 5 Empirical data in Primates

#### 5.1 Chain convergence

Obtained with the mechanistic inference model developed in this paper of site-specific amino-acid fitness profiles and log-Brownian process for $N_\mathrm{e}$, $\mu$ and life-history traits.

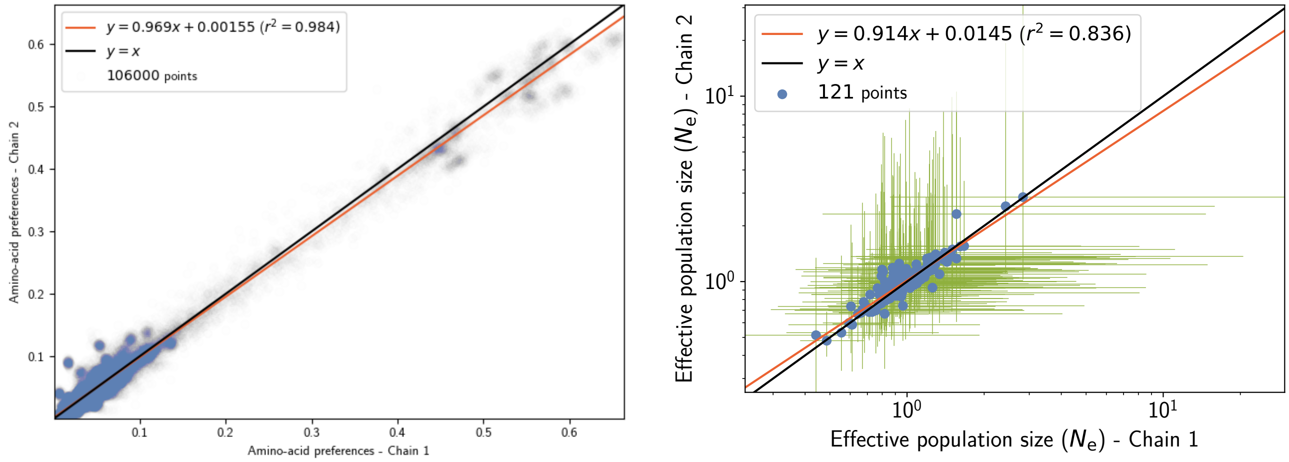


Figure 27: Chain convergence of site amino-acid preferences (left panel) and branch $N_\mathrm{e}$ (right panel).

#### 5.2 Traits estimation (chain 1)

Obtained with the mechanistic inference model developed in this paper of site-specific amino-acid fitness profiles and log-Brownian process for $N_\mathrm{e}$, $\mu$ and life-history traits.

| **Correlation ($\rho$)** | $N_\mathrm{e}$ | $\mu$ | **maturity** | **mass** | **longevity** | $\pi_S$ | $\pi_N/\pi_S$ | **generation time** |
|---|---|---|---|---|---|---|---|---|
| $N_\mathrm{e}$ | - | $-0.433^{**}$ | 0.155 | 0.166 | 0.157 | $-0.133$ | 0.104 | 0.16 |
| $\mu$ | - | - | $-0.792^{**}$ | $-0.791^{**}$ | $-0.773^{**}$ | $0.62^{**}$ | $-0.59$ | $-0.78^{**}$ |
| **maturity** | - | - | - | $0.986^{**}$ | $0.985^{**}$ | $-0.8^{**}$ | 0.746 | $0.991^{**}$ |
| **mass** | - | - | - | - | $0.977^{**}$ | $-0.737^{**}$ | 0.695 | $0.981^{**}$ |
| **longevity** | - | - | - | - | - | $-0.819^{**}$ | 0.752 | $0.999^{**}$ |
| $\pi_S$ | - | - | - | - | - | - | $-0.86^{**}$ | $-0.816^{**}$ |
| $\pi_N/\pi_S$ | - | - | - | - | - | - | - | 0.752 |
| **generation time** | - | - | - | - | - | - | - | - |

Table 16: Correlation coefficient between effective population size ($N_\mathrm{e}$), mutation rate per site per unit of time ($\mu$), and life-history traits (maximum longevity, adult weight and female maturity) were computed in primates. Asterisks indicate strength of support ($^{*}pp > 0.95$, $^{**}pp > 0.975$).

| Covariance ($\Sigma$) | $N_e$ | $\mu$ | maturity | mass | longevity | $\pi_S$ | $\pi_N/\pi_S$ | generation time |
|---|---|---|---|---|---|---|---|---|
| $N_e$ | 1.08** | −1.39** | 0.66 | 1.18 | 0.414 | −0.251 | 0.0898 | 0.452 |
| $\mu$ | - | 9.86** | −10.1** | −17.5** | −6.44** | 3.42** | −1.28 | −6.96** |
| maturity | - | - | 16.9** | 28.4** | 10.6** | −5.39** | 1.9 | 11.5** |
| mass | - | - | - | 49.8** | 18.1** | −8.89** | 3.29 | 19.5** |
| longevity | - | - | - | - | 6.99** | −3.75** | 1.31 | 7.47** |
| $\pi_S$ | - | - | - | - | - | 3.26** | −0.986** | −3.96** |
| $\pi_N/\pi_S$ | - | - | - | - | - | - | 0.419** | 1.39 |
| generation time | - | - | - | - | - | - | - | 8.02** |

Table 17: Correlation coefficient between effective population size ($N_e$), mutation rate per site per unit of time ($\mu$), and life-history traits (maximum longevity, adult weight and female maturity) were computed in primates. Asterisks indicate strength of support ($^*pp > 0.95$, $^{**}pp > 0.975$).

| Partial coefficient | $N_e$ | $\mu$ | maturity | mass | longevity | $\pi_S$ | $\pi_N/\pi_S$ | generation time |
|---|---|---|---|---|---|---|---|---|
| $N_e$ | - | −0.411 | −0.0622 | 0.0184 | −0.0436 | −0.0482 | −0.00476 | 0.0333 |
| $\mu$ | - | - | 0.0548 | −0.101 | 0.146 | −0.0134 | −0.102 | −0.124 |
| maturity | - | - | - | 0.292 | −0.793** | −0.167 | 0.0547 | 0.824** |
| mass | - | - | - | - | −0.0589 | 0.43 | −0.195 | 0.101 |
| longevity | - | - | - | - | - | −0.159 | −0.148 | 0.991** |
| $\pi_S$ | - | - | - | - | - | - | −0.573** | 0.11 |
| $\pi_N/\pi_S$ | - | - | - | - | - | - | - | 0.144 |
| generation time | - | - | - | - | - | - | - | - |

Table 18: Partial correlation coefficient between Neffective population size ($N_e$), mutation rate per site per unit of time ($\mu$), and life-history traits (maximum longevity, adult weight and female maturity) were computed in primates. Asterisks indicate strength of support ($^*pp > 0.95$, $^{**}pp > 0.975$).

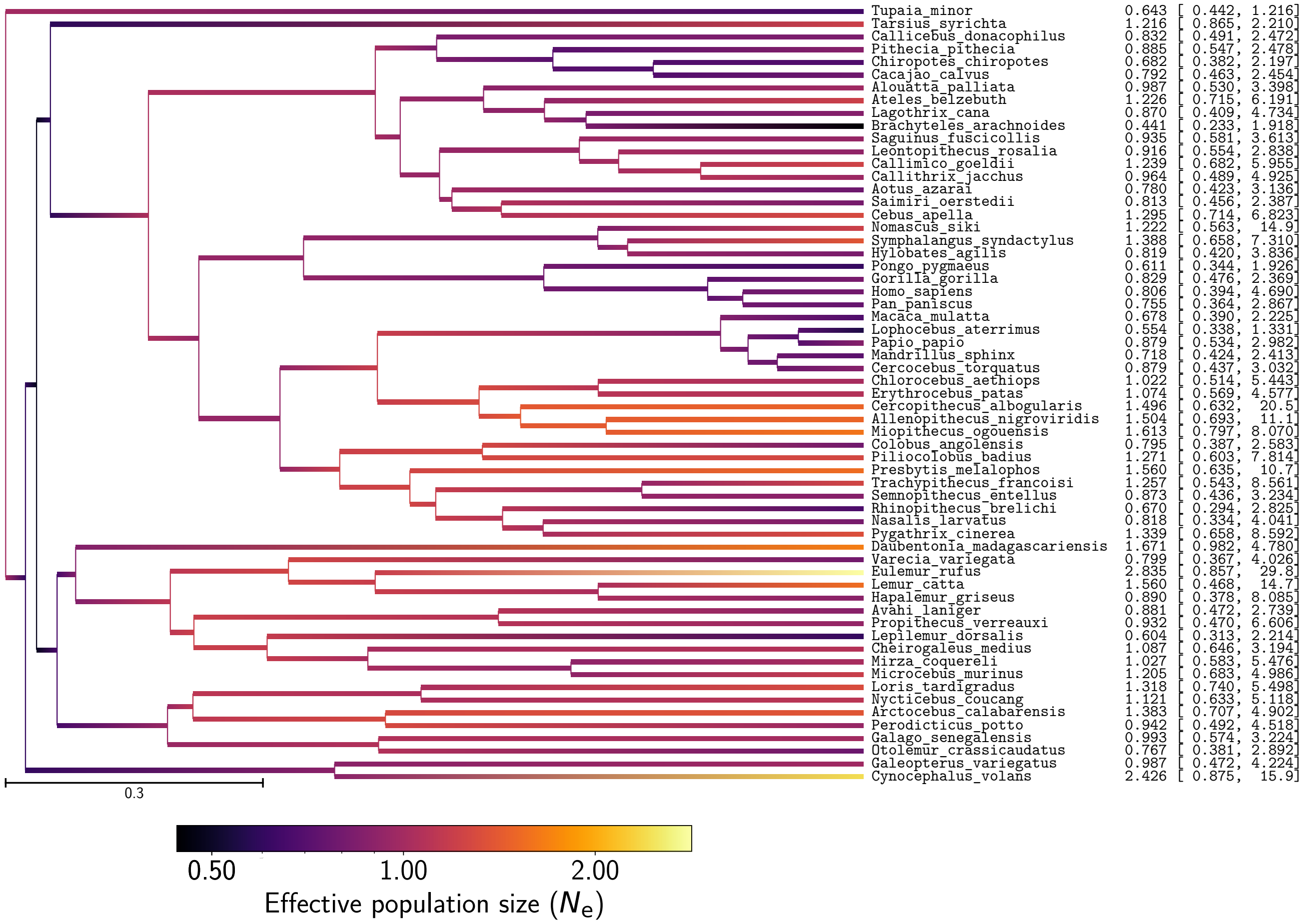


Figure 28: Effective population size ($N_e$) estimation in primates

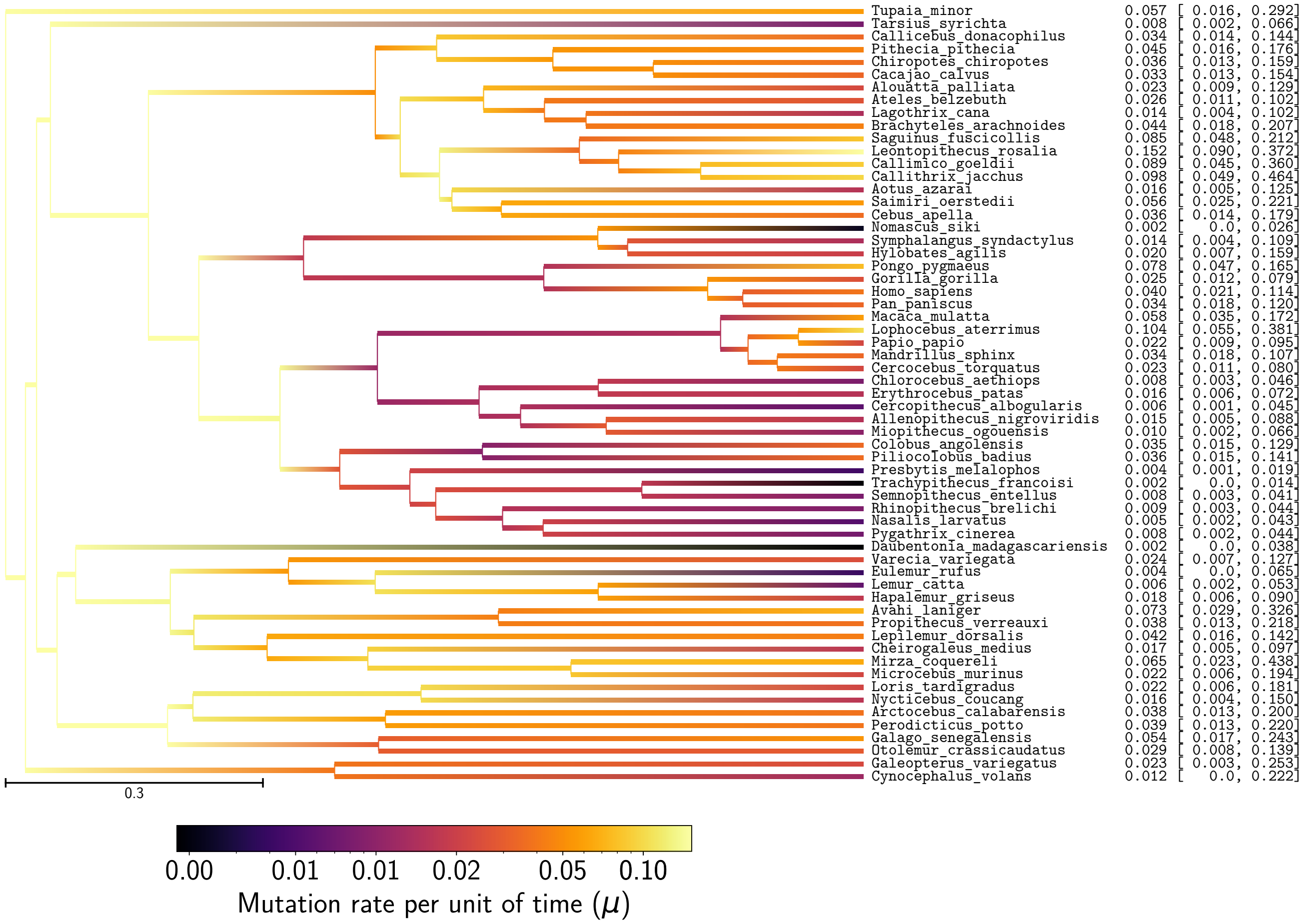


Figure 29: Mutation rate ($\mu$) estimation in primates

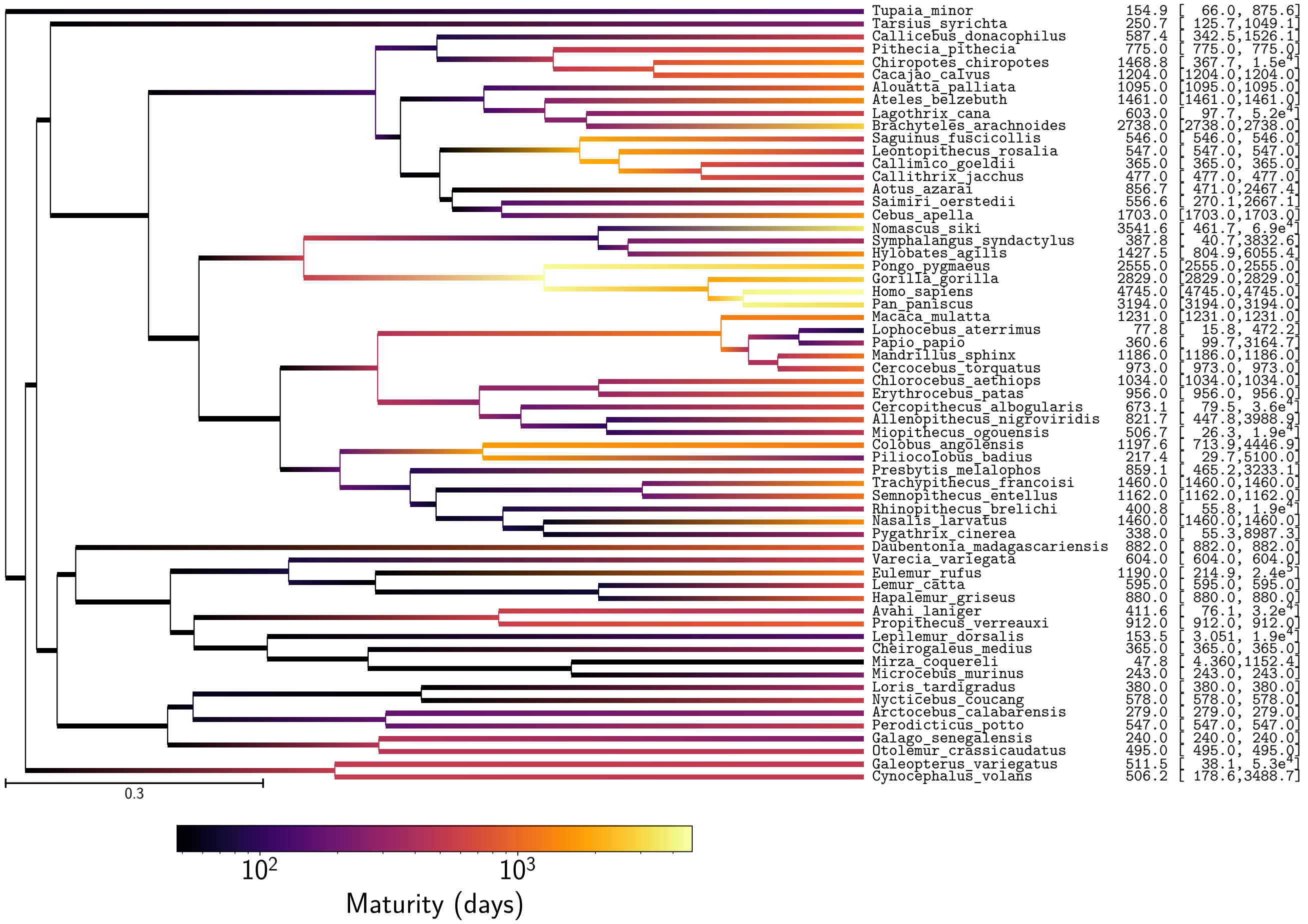


Figure 30: Female maturity estimation in primates

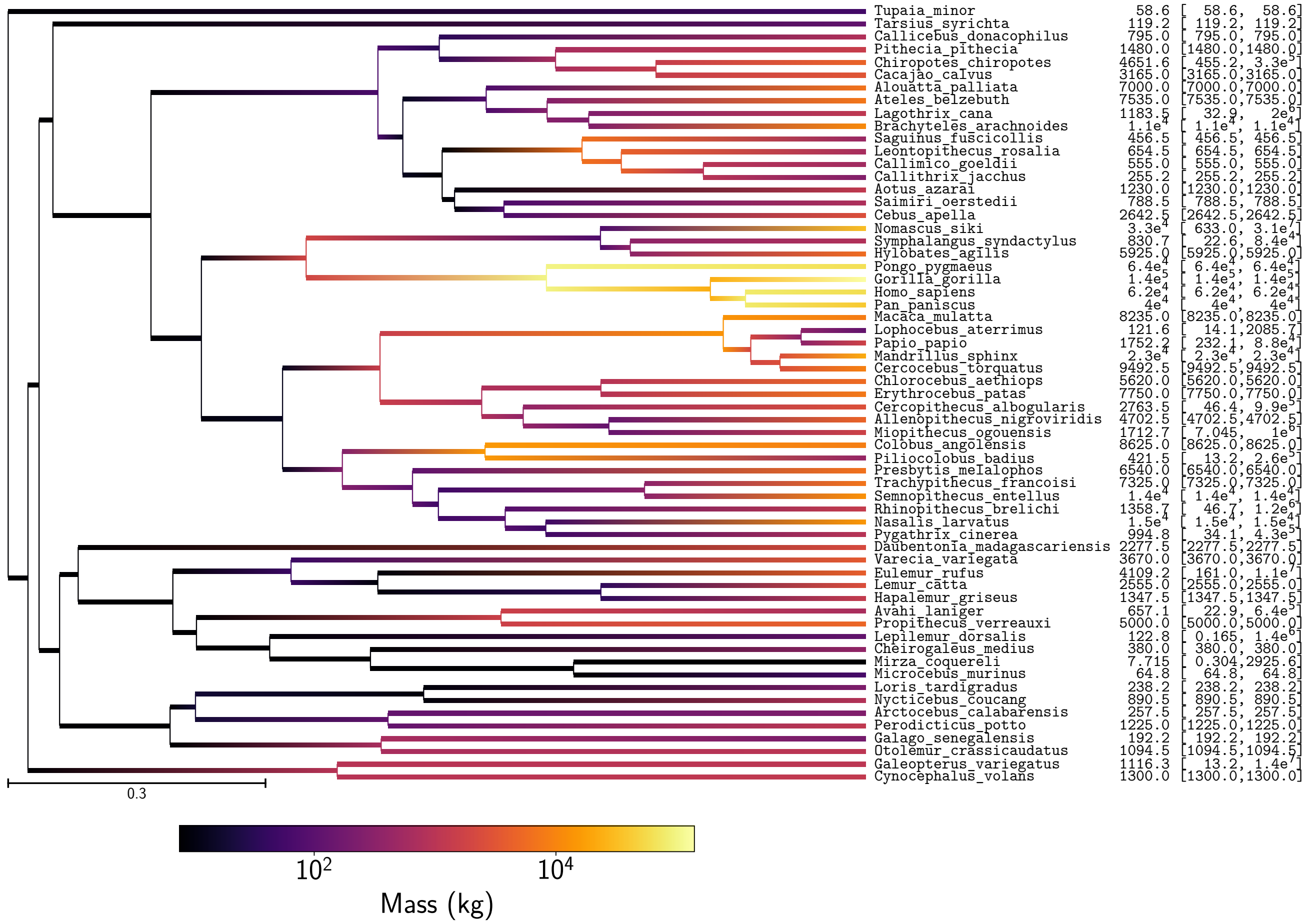


Figure 31: Mass estimation in primates

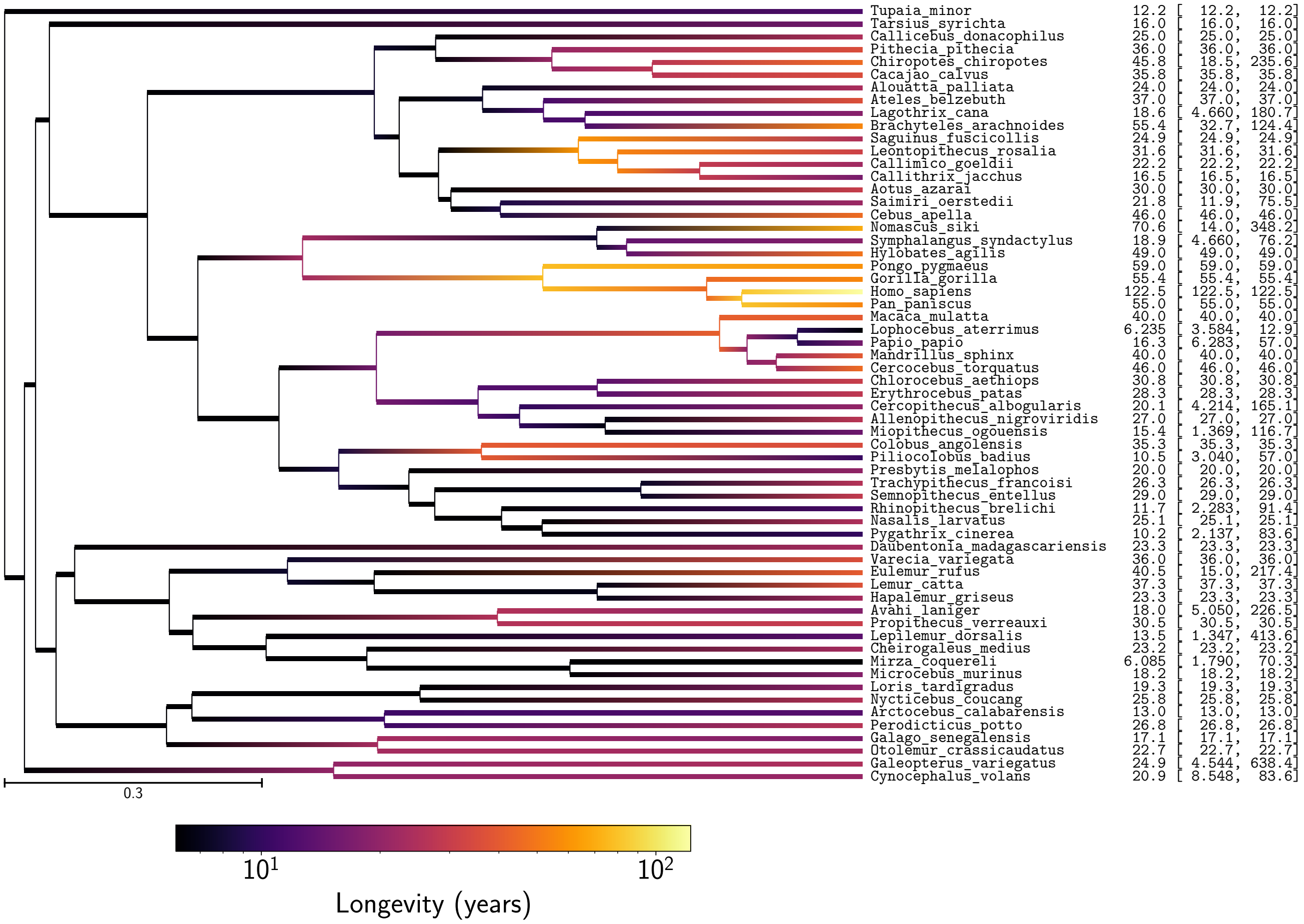


Figure 32: Longevity estimation in primates

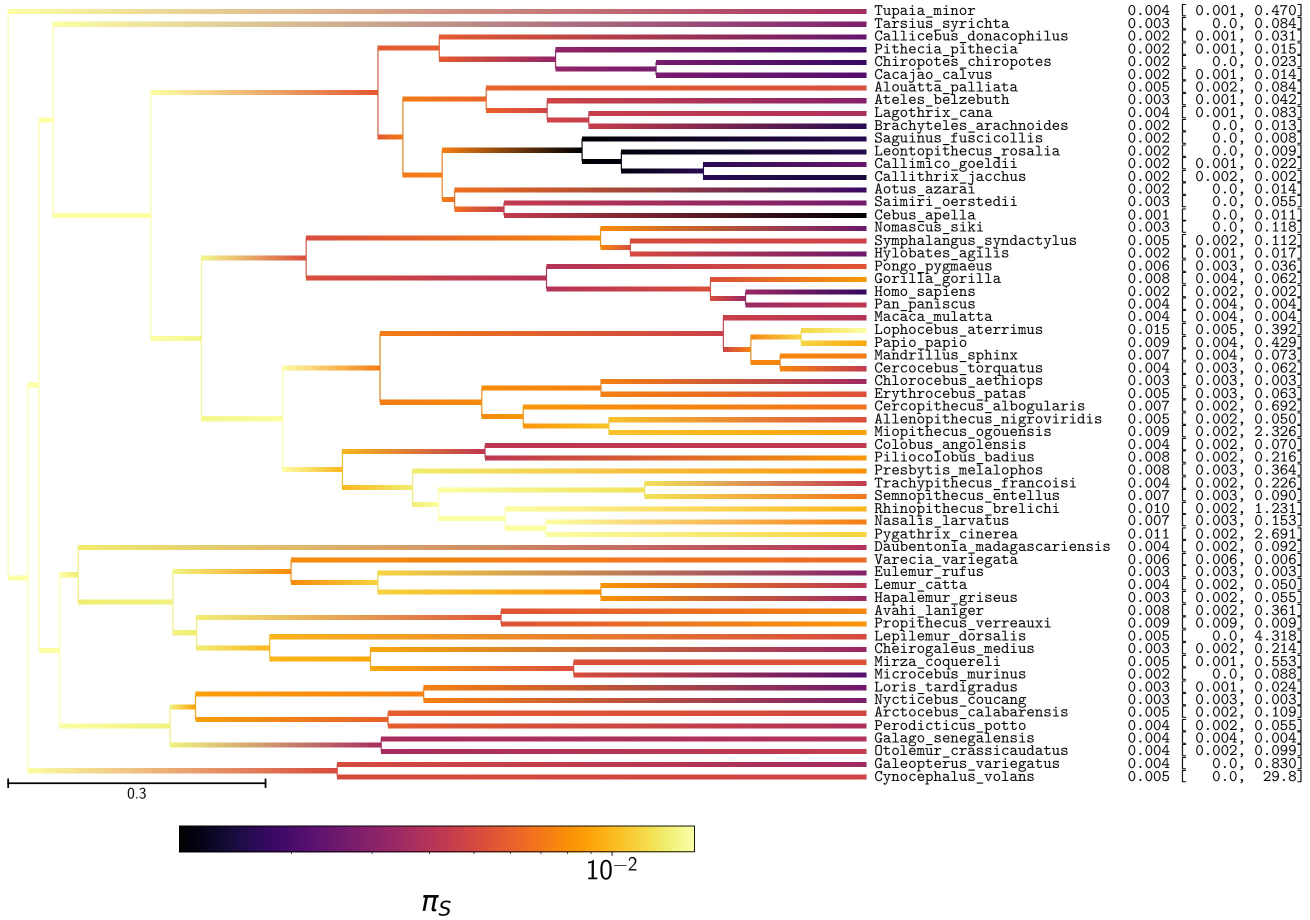


Figure 33: $\pi_S$ estimation in primates

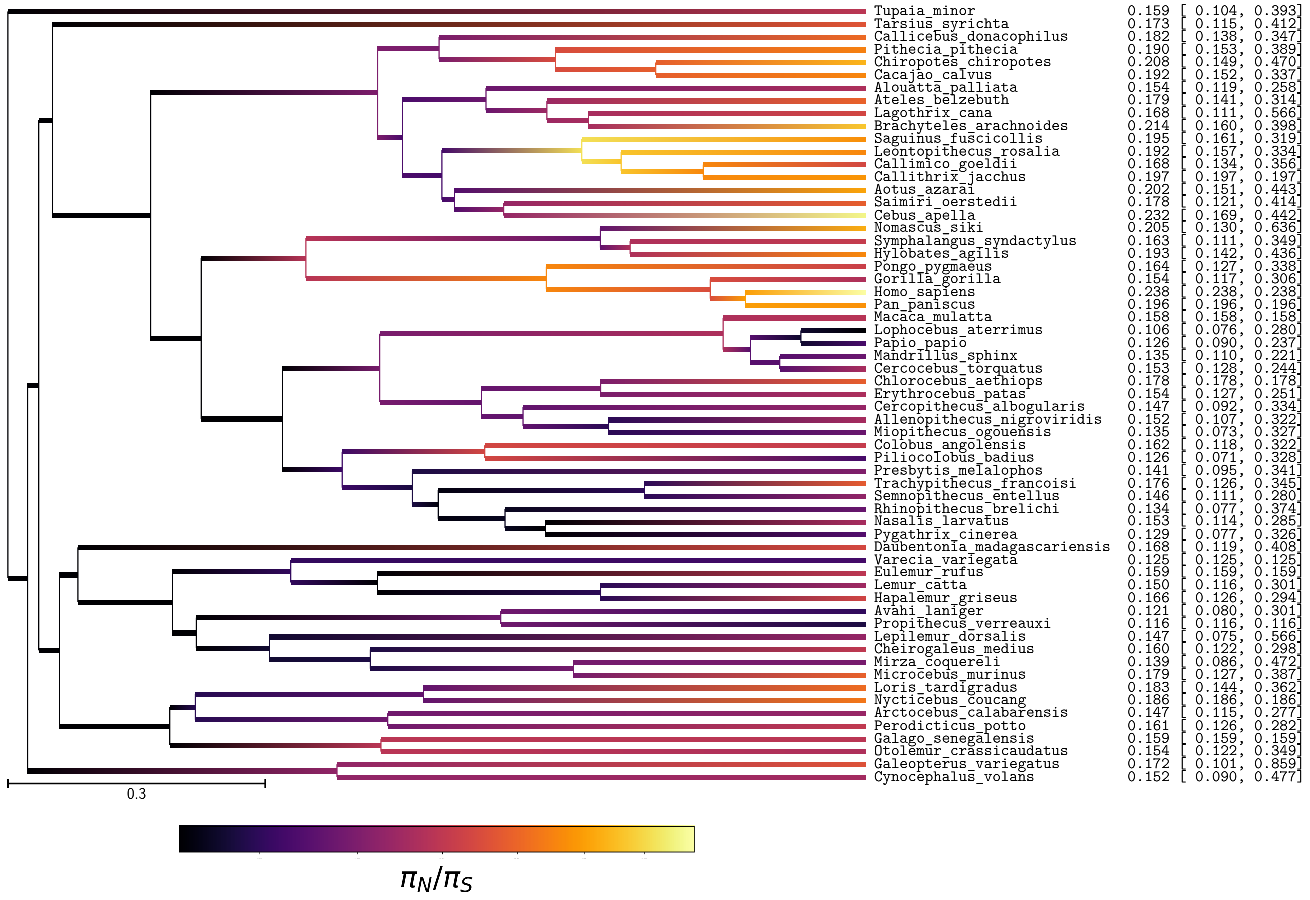


Figure 34: $\pi_N/\pi_S$ estimation in primates

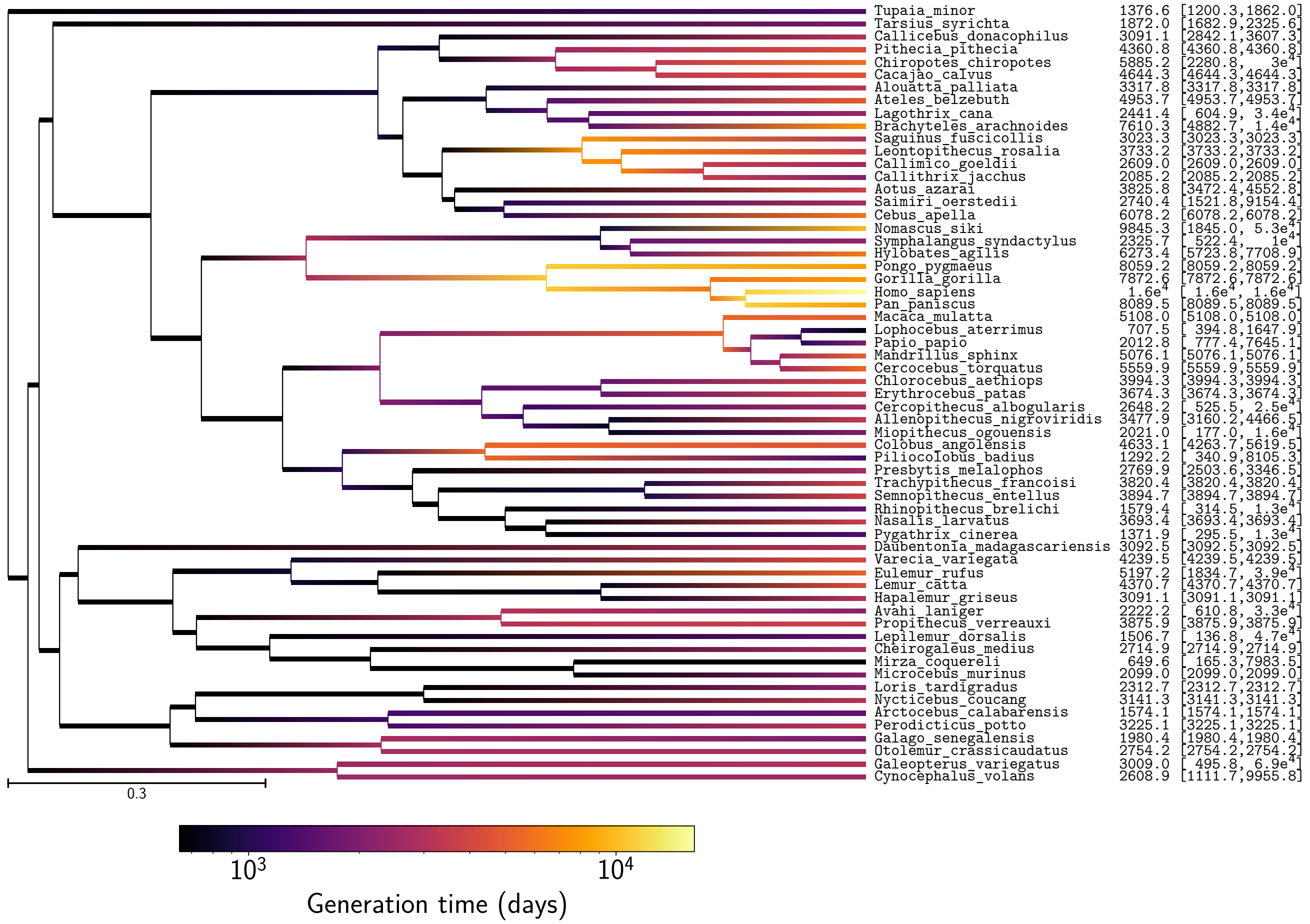


Figure 35: Generation time estimation in primates

#### 5.3 Amino-acid preferences entropy

| Experiment | $\langle \Omega \rangle$ (branch $N_e$) | $\langle \Omega \rangle$ (constant $N_e$) |
|---|---|---|
| Primates, chain 1 | $1.41 \pm 0.10$ | $1.49 \pm 0.08$ |
| Primates, chain 2 | $1.40 \pm 0.10$ | $1.48 \pm 0.08$ |

Table 19: Estimated amino-acid entropy in primates. Obtained with the mechanistic inference model developed in this paper of site-specific amino-acid fitness profiles and log-Brownian process for $N_e$, $\mu$ and life-history traits (in the left column), or under the assumption of constant $N_e$ (in the right column).

#### 5.4 Traits estimation with branch $\omega$ (chain 1)

Obtained with the phenomenological inference model of log-Brownian process for the $\mu$ and the relative non-synonymous substitution rate ($\omega$), as in Lartillot and Poujol (2011).

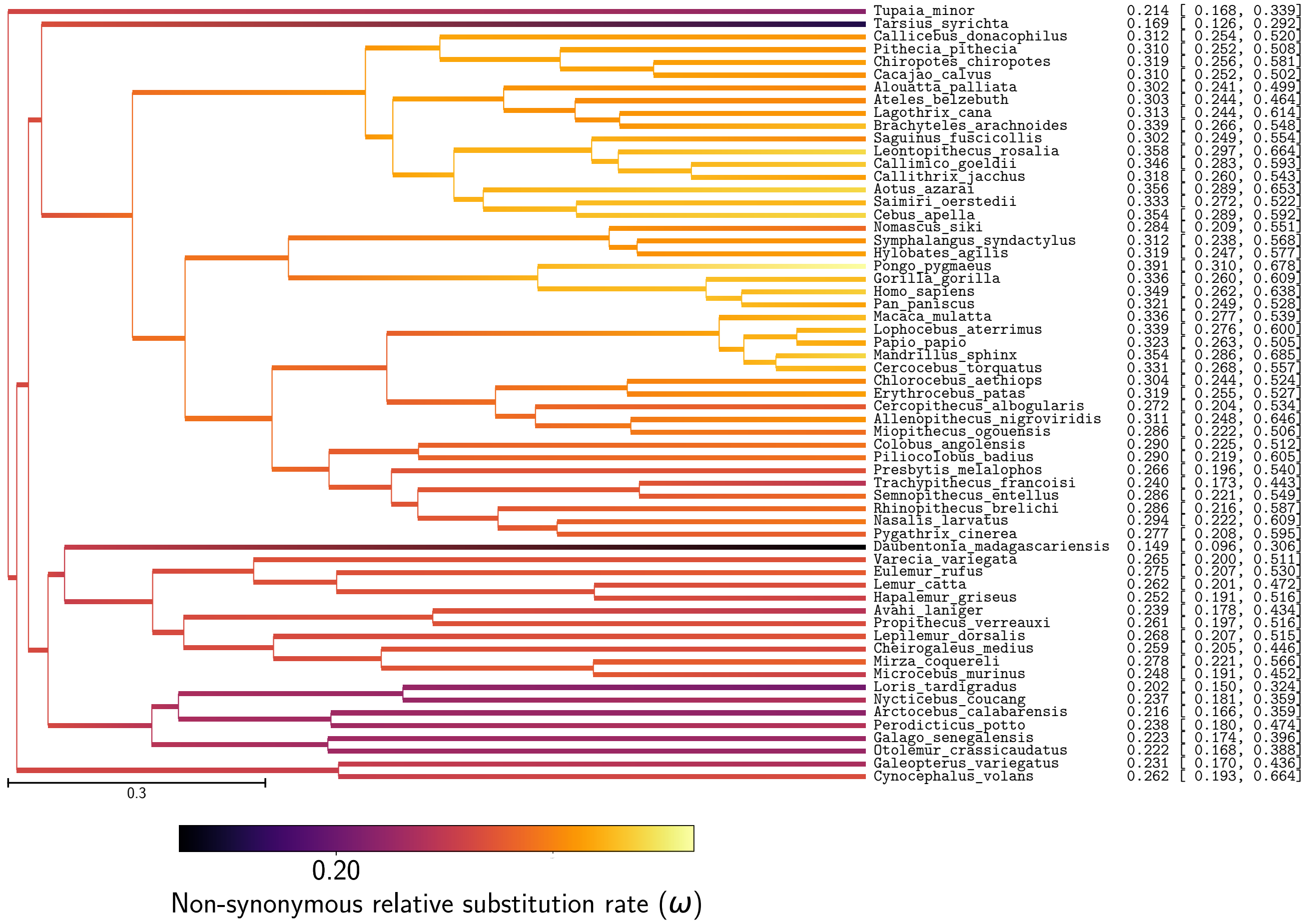


Figure 36: Non-synonymous substitution rate ($\omega$) estimation in primates

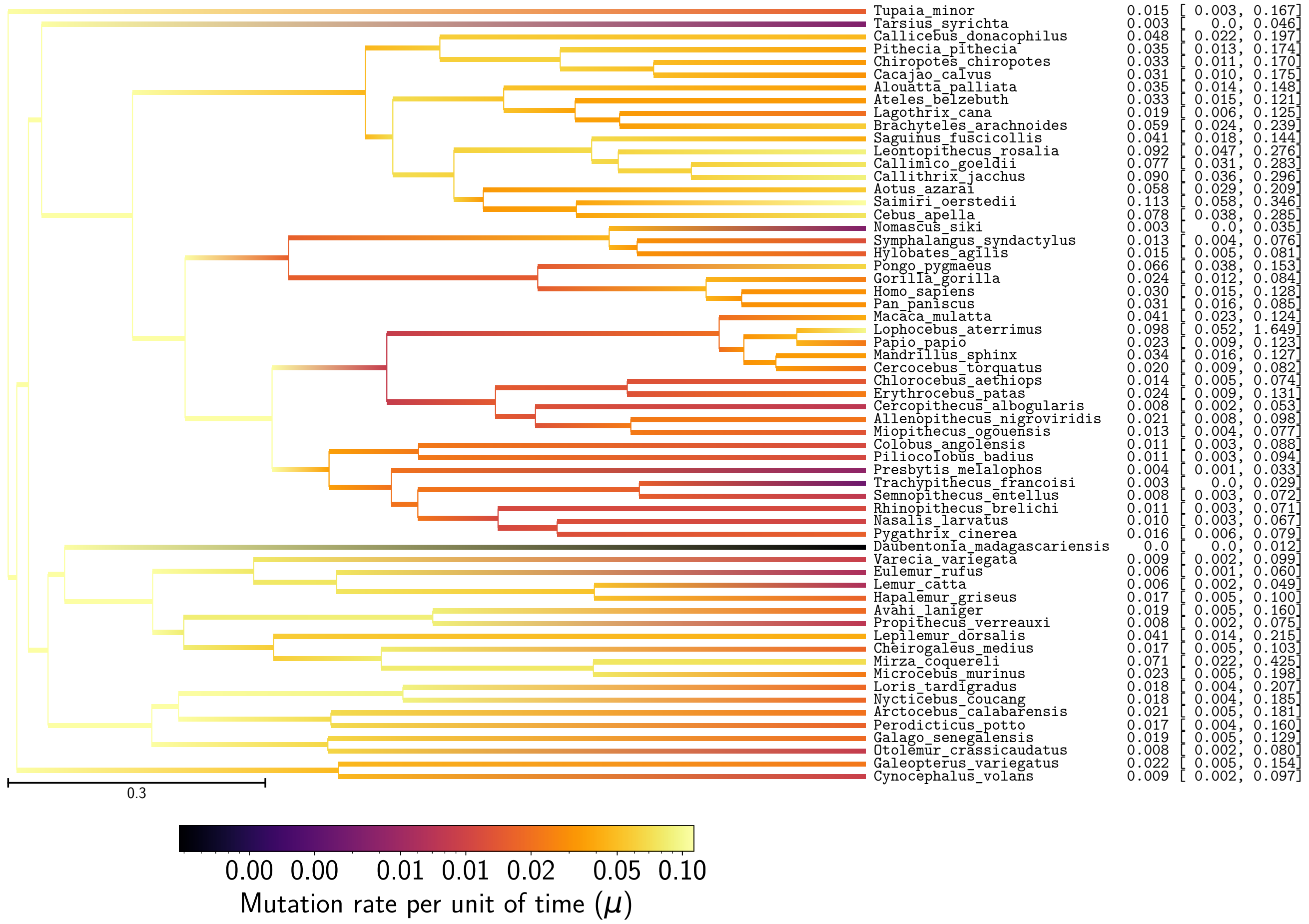


Figure 37: Mutation rate ($\mu$) estimation in primates

| **Correlation ($\rho$)** | $\omega$ | $\mu$ | **maturity** | **mass** | **longevity** | $\pi_S$ | $\pi_N/\pi_S$ | **generation time** |
|---|---|---|---|---|---|---|---|---|
| $\omega$ | - | 0.294 | 0.000316 | 0.0361 | 0.0155 | $-0.197$ | 0.145 | 0.0111 |
| $\mu$ | - | - | $-0.804^{**}$ | $-0.798^{**}$ | $-0.817^{**}$ | $-0.0201$ | 0.031 | $-0.823^{**}$ |
| **maturity** | - | - | - | $0.952^{**}$ | $0.957^{**}$ | $-0.166$ | 0.162 | $0.97^{**}$ |
| **mass** | - | - | - | - | $0.933^{**}$ | $-0.0437$ | 0.0427 | $0.943^{**}$ |
| **longevity** | - | - | - | - | - | $-0.223$ | 0.165 | $0.999^{**}$ |
| $\pi_S$ | - | - | - | - | - | - | $-0.664$ | $-0.212$ |
| $\pi_N/\pi_S$ | - | - | - | - | - | - | - | 0.162 |
| **generation time** | - | - | - | - | - | - | - | - |

Table 20: Correlation coefficient between non-synonymous substitution rate ($\omega$), mutation rate per site per unit of time ($\mu$), and life-history traits (maximum longevity, adult weight and female maturity) were computed in primates. Asterisks indicate strength of support ($^*pp > 0.95$, $^{**}pp > 0.975$).

| Covariance ($\boldsymbol{\Sigma}$) | $\omega$ | $\mu$ | maturity | mass | longevity | $\pi_S$ | $\pi_N/\pi_S$ | generation time |
|---|---|---|---|---|---|---|---|---|
| $\omega$ | $0.0674^{**}$ | 0.231 | −0.0106 | 0.0149 | −0.00138 | −0.0435 | 0.0101 | −0.00314 |
| $\mu$ | - | $8.71^{**}$ | $-4.8^{**}$ | $-9.22^{**}$ | $-3.97^{**}$ | 0.188 | 0.0483 | $-4.08^{**}$ |
| maturity | - | - | $4.95^{**}$ | $8.37^{**}$ | $3.29^{**}$ | −1.01 | 0.000924 | $3.53^{**}$ |
| mass | - | - | - | $16.3^{**}$ | $6.14^{**}$ | −0.932 | −0.0741 | $6.45^{**}$ |
| longevity | - | - | - | - | $2.76^{**}$ | −0.577 | 0.0919 | $2.82^{**}$ |
| $\pi_S$ | - | - | - | - | - | $1.3^{**}$ | −0.148 | −0.637 |
| $\pi_N/\pi_S$ | - | - | - | - | - | - | $0.182^{**}$ | 0.0775 |
| generation time | - | - | - | - | - | - | - | $2.92^{**}$ |

Table 21: Correlation coefficient between non-synonymous substitution rate ($\omega$), mutation rate per site per unit of time ($\mu$), and life-history traits (maximum longevity, adult weight and female maturity) were computed in primates. Asterisks indicate strength of support ($^*pp > 0.95$, $^{**}pp > 0.975$).

| Partial coefficient | $\omega$ | $\mu$ | maturity | mass | longevity | $\pi_S$ | $\pi_N/\pi_S$ | generation time |
|---|---|---|---|---|---|---|---|---|
| $\omega$ | - | 0.463 | −0.0461 | 0.248 | −0.027 | −0.193 | −0.0681 | 0.0319 |
| $\mu$ | - | - | 0.0649 | −0.000258 | 0.0374 | −0.128 | 0.115 | −0.075 |
| maturity | - | - | - | 0.228 | $-0.834^{**}$ | −0.0991 | 0.0491 | $0.854^{**}$ |
| mass | - | - | - | - | −0.038 | 0.435 | −0.123 | 0.0851 |
| longevity | - | - | - | - | - | −0.184 | −0.145 | $0.994^{**}$ |
| $\pi_S$ | - | - | - | - | - | - | $-0.553^{*}$ | 0.125 |
| $\pi_N/\pi_S$ | - | - | - | - | - | - | - | 0.136 |
| generation time | - | - | - | - | - | - | - | - |

Table 22: Partial correlation coefficient between non-synonymous substitution rate ($\omega$), mutation rate per site per unit of time ($\mu$), and life-history traits (maximum longevity, adult weight and female maturity) were computed in primates. Asterisks indicate strength of support ($^*pp > 0.95$, $^{**}pp > 0.975$).

### 6 Sufficient statistics

A sequence of length Z evolves by point substitutions, according to a random process defined by the substitution matrices $\boldsymbol{Q}^{(\mathrm{b,z})}$, over a phylogenetic tree. A realization of the random process along a branch b, and at a particular site z results in a detailed substitution history, which will be denoted by $\mathcal{H}^{(\mathrm{b,z})}$.

#### 6.1 Path sufficient statistics

All sites owning to the same category of fitness profile share the same substitution rate matrix. Hence, $\mathcal{H}^{(\mathrm{b,z})}$ can be gathered across all sites owing to a specific category k, denoted $\mathcal{H}^{(\mathrm{b})}$. If we express the probability of the substitution mapping ($\mathcal{H}^{(\mathrm{b,k})}$) as a function of the codon substitution process for this category k, we get the following expression:

$$\mathbb{P}(\mathcal{H}^{(\mathrm{b,k})}|l^{(\mathrm{b})}, \boldsymbol{Q}^{(\mathrm{b,k})}) \propto \left[\prod_{i=1}^{61}\left[\pi_i^{(\mathrm{b,k})}\right]^{n_i^{(\mathrm{b,k})}}\right] \cdot \left[\prod_{1\leq i,j\leq 61}\left[Q_{i,j}^{(\mathrm{b,k})}\right]^{m_{i,j}^{(\mathrm{b,k})}}\right] \cdot \left[\prod_{i=1}^{61} \mathrm{e}^{-\left|Q_{i,i}^{(\mathrm{b,k})}\right| a_i^{(\mathrm{b,k})}}\right], \tag{17}$$

where we define the sufficient statistics:

- $m_{i,j}^{(\mathrm{b,k})}$ is the total number of substitutions from codon $i$ to codon $j$
- $n_i^{(\mathrm{b,k})}$ is the number of sites starting with codon $i$ at the tip of the branch.
- $a_i^{(\mathrm{b,k})}$ is the total waiting time in codon $i$.

Once these sufficient statistics have been computed, the parameters of the substitution matrix $\boldsymbol{Q}^{(\mathrm{b,k})}$ can be resampled conditional on $\mathcal{H}^{(\mathrm{b,k})}$, using equation 17 each time the likelihood needs to be recomputed. This leads to relatively fast MCMC strategy.

#### 6.2 Length sufficient statistics

$\mathcal{H}^{(\mathrm{b,z})}$ can also be gathered across all sites along a specific branch, giving $\mathcal{H}^{(\mathrm{b})}$. Then the probability of the substitution history given the branch lengths ($l^{(\mathrm{b})} = \mu^{(\mathrm{b})}\Delta T^{(\mathrm{b})}$), takes a very simple form:

$$\mathbb{P}(\mathcal{H}^{(\mathrm{b})}|L^{(\mathrm{b})}) \propto \left[L^{(\mathrm{b})}\right]^{u^{(\mathrm{b})}} \mathrm{e}^{-r^{(\mathrm{b})}L^{(\mathrm{b})}}, \tag{18}$$

where we define the sufficient statistics:

- $u^{(\mathrm{b})}$ is the total number of substitutions over branch b, summed over all sites.
- $r^{(\mathrm{b})}$ is the mean rate away from current codon state (averaged over the entire substitution history).

Thus, formally, the probability of the substitution mapping can be summarized by saying that the total number of substitutions along a given branch over all sites, $u^{(\mathrm{b})}$, is Poisson distributed, of mean $r^{(\mathrm{b})}L^{(\mathrm{b})}$.

#### 6.3 Scatter sufficient statistics

From the independent contrast $\boldsymbol{C}^{(\mathrm{b})}$ of the Brownian process $\boldsymbol{B}^{(\mathrm{n})}$, we can define the $2 \times 2$ scatter sufficient statistic matrix, $\boldsymbol{A}$ as:

$$\boldsymbol{A} = \sum_{b=1}^{2P-2} \boldsymbol{C}^{(\mathrm{b})} \cdot \left[\boldsymbol{C}^{(\mathrm{b})}\right]^{\mathrm{T}} \tag{19}$$

By Bayes theorem, the posterior on $\boldsymbol{\Sigma}$, conditional on a particular realization of $B$ (and thus of $\boldsymbol{C}$) is an invert Wishart distribution, of parameter $\kappa\boldsymbol{I} + \boldsymbol{A}$ and with $2P - 2 + 3$ degrees of freedom.

$$\boldsymbol{\Sigma} \sim \text{Wishart}^{-1}\left(\kappa\boldsymbol{I} + \boldsymbol{A}, 2P - 2 + 3\right) \tag{20}$$

This invert Wishart distribution can be obtained by sampling $2P - 2 + 3$ independent and identically distributed multivariate normal random variables $\boldsymbol{Z}^{(a)}$ defined by

$$\boldsymbol{Z}^{(a)} \sim \mathcal{N}\left(\boldsymbol{0}, [\kappa\boldsymbol{I} + \boldsymbol{A}]^{-1}\right). \tag{21}$$

And from these multivariate samples, $\boldsymbol{\Sigma}$ is Gibbs sampled as:

$$\boldsymbol{\Sigma} = \left(\sum_{k=1}^{2P-2+3} \boldsymbol{Z}^{(a)} \cdot \left[\boldsymbol{Z}^{(a)}\right]^{\mathrm{T}}\right)^{-1} \tag{22}$$
